# A genome-scale drug discovery pipeline uncovers new therapeutic targets and a unique p97 allosteric binding site in *Schistosoma mansoni*

**DOI:** 10.1101/2025.03.14.643303

**Authors:** Dylon R Stephens, Ho Yee Joyce Fung, Yan Han, Jue Liang, Zhe Chen, Joseph Ready, James J Collins

## Abstract

Schistosomes are parasitic flatworms that infect more than 200 million people globally. However, there is a shortage of molecular tools that enable the discovery of potential drug targets within schistosomes. Thus, praziquantel has remained the frontline treatment for schistosomiasis despite known liabilities. Here, we have conducted a genome-wide study in *S. mansoni* using the human druggable genome as a bioinformatic template to identify essential genes within schistosomes bearing similarity to catalogued drug targets. Then, we assessed these candidate targets *in silico* using a set of unbiased criteria to determine which possess ideal characteristics for a ready-made drug discovery campaign. Following this prioritization, we pursued a parasite p97 ortholog as a bona-fide drug target for the development of therapeutics to treat schistosomiasis. From this effort, we identified a covalent inhibitor series that kills schistosomes through an on-target killing mechanism by disrupting the ubiquitin proteasome system. Fascinatingly, these inhibitors induce a conformational change in the conserved D2 domain P-loop of schistosome p97 upon modification of Cys519. This conformational change reveals an allosteric binding site adjacent to the D2 domain active site reminiscent of the ‘DFG’ flip in protein kinases. This allosteric binding site can potentially be utilized to generate new classes of species-selective p97 inhibitors. Furthermore, these studies provide a resource for the development of alternative therapeutics for schistosomiasis and a workflow to identify potential drug targets in similar systems with few available molecular tools.

**Significance Statement:** Schistosomes cause widespread infections in humans, leading to severe chronic illnesses in endemic regions. There is no vaccine for schistosomiasis, and there has been limited success using the current standard-of-care treatment, praziquantel. Therefore, it is essential to identify drug targets within these parasites. Here, we identify potential drug targets in schistosomes bearing similarity to established human therapeutic targets, evaluate their essentiality for parasite survival, then prioritize them using an unbiased set of criteria to uncover high-value targets for the treatment of schistosomiasis. We investigated one candidate as a proof-of concept, a p97 ortholog, to discover newly characterized inhibitors of the parasite enzyme. This study demonstrates that this workflow can lead to the identification of small molecules that kill schistosomes.

## Introduction

Schistosomiasis affects more than 200 million of the world’s poorest people^1^, claiming the lives of ~250,000 people yearly^2^, while placing a significant clinical and economic burden on endemic regions^1^. Among the most practical challenges facing eradication of schistosomiasis is that there is no vaccine, and treatment has relied on a single drug, praziquantel (PZQ), for over 40 years.

Despite its exclusive use in the treatment of schistosomiasis since the 1970s^3–5^, PZQ possesses prominent and persistent pharmaceutical and pharmacological liabilities. For instance, cure rates following PZQ treatment vary dramatically in regions with high infection rates^6–8^. Because of this, there are valid concerns that mechanisms of PZQ resistance will become widespread. Indeed, reduced sensitivity can be induced rapidly in a laboratory setting^9,10^, and loss of function mutations in a PZQ-sensitive ion channel have already been identified in the field^11–13^. As PZQ has proven unable to lead to the elimination of schistosomiasis in endemic regions on its own^14^, serious dedicated efforts are required identify drug targets for the development of alternative therapeutics to PZQ.

However, identification of novel schistosomicidals has proven difficult. The relatively large size of adult worms (~1cm) prevent them from being adapted to micronized volumes amenable to high-throughput phenotypic screens, and molecules derived from screening against larval parasites suffer from high rates of attrition when examined in adult worms^15–17^. Because of these limitations in identifying chemicals that can kill the adult worms that drive disease pathology, an attractive alternative is the discovery of lead compounds through target-based approaches. As yet, there are few validated drug targets within schistosomes^18–23^. Even with progress in the development of large-scale RNAi approaches^24^, a paucity of facile molecular tools has hindered the systematic identification and subsequent functional validation of target proteins in the worm. Deciphering the function of essential genes still requires a large investment in labor and time. And, while there is an obvious requirement that drug targets be essential, essentiality alone is not sufficient to anticipate which targets will lead to successful drug discovery efforts. Many target-based screening efforts failed because the selected targets were unable to bind drug-like molecules with reasonable affinity^25–27^.

This paper expands upon our previous work by 1) identifying essential genes within the *S. mansoni* genome that bear homology to well-defined drug targets in the context of human disease using bioinformatic and genetic approaches and 2) assessing these genetically essential targets for viability as ‘bona fide’ drug targets utilizing well-defined criteria. Combining our findings from RNAi experiments in this study with those performed previously^24^, we have prioritized 18 potential drug targets using an unbiased set of criteria. We investigated one of these targets in-depth, p97, a gene encoding a AAA-ATPase involved in the degradation of proteins via the Ubiquitin Proteasome System (UPS)^28^. Following high-throughput screening efforts, we identified a benzoxazole propiolamide covalent scaffold that inhibits the enzyme, exhibits selectivity for the parasite ortholog over its human counterpart, and displays on-target effects in adult parasites via accumulation of K48 polyubiquitinated proteins^24,29^. Furthermore, we obtained a structure of p97 bound to these covalent inhibitors using cryo-EM, identifying a previously unreported conformational change in the nucleotide (Walker A) binding motif of p97 within its D2 domain^30^. This conformational change creates a novel allosteric pocket between p97 monomers that may enable species-selective inhibitors to be developed. Together, these studies highlight a validated set of high-value targets that will serve as the foundation for future drug discovery campaigns.

## Results

### Large-scale RNAi screen reveals 63 genes essential for *in vitro* parasite survival

With the success of previous RNAi studies^24^, we revisited the *S. mansoni* genome to determine other potentially attractive targets that remain uncharacterized. To improve our chances of success in finding essential genes, as well as potential drug targets, we utilized known “druggable” targets catalogued in databases such as DrugBank^31^, ChEMBL^32^, and TTD^33^ (**Fig. 1*A* and SI Appendix, Table S1**).

**Figure 1.**
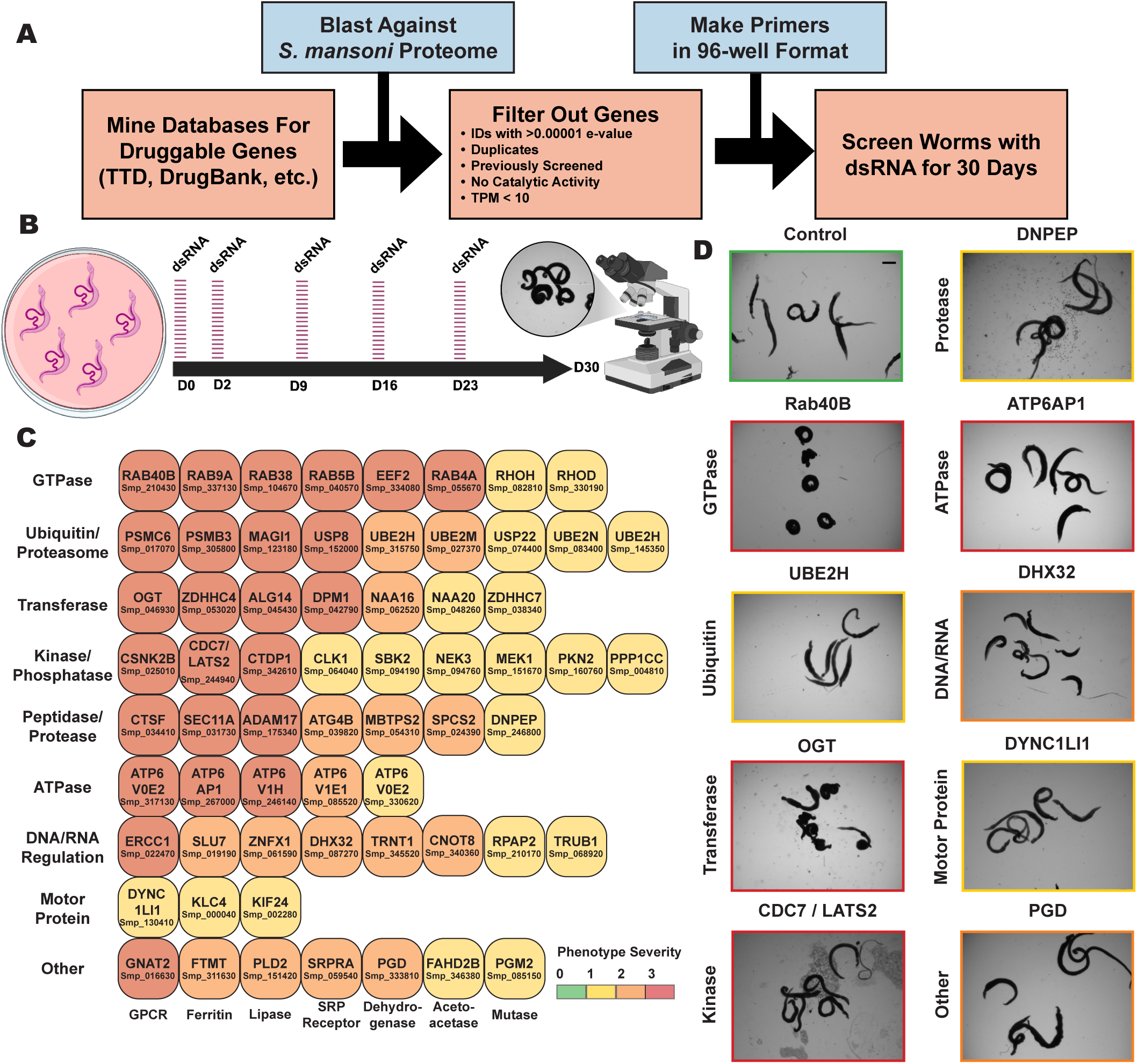
Large-scale RNAi screen targeting [Human] druggable genes. **(A)** Bioinformatic pipeline for the identification of potentially druggable genes. Target genes were identified using databases such as TTD, DrugBank, Chembl, etc. BLAST-p was used to identify similar proteins in the *S. mansoni* proteome, then passed through several preliminary qualifications before being included in the screen. Primers possessing a T7 promoter in addition to a 500-1000 bp target gene region were created to generate dsRNA for final candidate genes in 96-well format before treatment. **(B)** RNAi treatment schedule. Parasites were treated with dsRNA on D0 and D2, then every 7 days thereafter (D9, 16, and 23). Gross morphology was monitored over the course of 30 days by light microscopy. Observations on substrate attachment, movement, and morphology were taken every 1-2 days coinciding with media changes. Images were taken on Day 30 to assess final phenotypes. **(C)** Categorical arrangement of essential genes producing phenotypes according to enzymatic activity/biological function. Severity of phenotypes that appeared were scored based on how early the phenotype began to appear (D7-D15, D16-D25, and post-D25), as well as the number of visual phenotypes that manifested (tissue/gut edema, tegument/head degeneration, hypercontraction, death, etc.). Scoring follows description for RNAi Severity in Table 1. Red (3) represents a gene who presented more than one physical phenotype and appeared between D1-D15. Orange (2) represents a gene that presented more than one phenotype between D16-D25. Yellow (1) denotes a gene with an intermediate onset (D16-D25) of a modest phenotype. Green (0) represents a modest phenotype that appears after D25. **(D)** Representative images of phenotypes based on enzyme category compared to control dsRNA treatment (pJC53.2). GTPase RAB40B (Smp_210430), Ubiquitin transferase UBE2H (Smp_145350), Protein Kinase CDC7/LATS2 (Smp_244940), Glycosyltransferase OGT (Smp_046930), RNA helicase DHX32 (Smp_130410), V-type proton ATPase subunit ATP6AP1 (Smp_267000), Aminopeptidase DNPEP (Smp_246800), Dynein-light chain DYNC1LI1 (Smp_130410), and 6-Phosphogluconate dehydrogenase PGD (Smp_333810). Scale bar **(D)**, 1,000 μm.

**Table 1.** *In silico* target priority score categories. Scoring criteria rubric for the prioritization of essential schistosome genes identified by RNAi studies. There are five categories: RNAi Severity Score (RSS), Essentiality Score (ES) (Mouse and Human), Assayability Score (AS), Druggability Score (DS), and Parasite Selectivity Score (PSS). Targets were scored on a scale of ideal (3) to unideal (0) characteristics in each category. *Target genes were only assessed for parasite selectivity scoring through sequence alignments and 3D modelling if they had favorable scores in each of the other categories (≥10).

|  |  |
| --- | --- |
| **<u>RNAi Severity Score (RSS)</u>** |  |
| Modest phenotype, not seen until D25 | 0 |
| Intermediate Onset (D16-D25), modest phenotype | 1 |
| Intermediate Onset (D16-D25), more than one phenotype | 2 |
| Rapid Onset (D7-D15), more than one phenotype | 3 |
| **<u>Essentiality Score (ES)</u>** |  |
| ***Mouse Essentiality Score (Mouse ES)*** |  |
| Essential in mice | 0 |
| No information available for mice | 1 |
| Essential, not embryonic lethal | 2 |
| Non-essential in mice | 3 |
| ***Human Essentiality Score (Human ES)*** |  |
| Essential in human cells | 0 |
| No information available | 1 |
| Essential in some human cell lines | 2 |
| Non-essential in human cell lines | 3 |
| **<u>Assayability Score (AS)</u>** |  |
| No clear path to recombinant purification or assay development | 0 |
| Purification and assay possible, will require extensive optimization | 1 |
| Purification and assay possible, will require some optimization | 2 |
| Established purification schemes and assay available using commercial reagents | 3 |
| **<u>Druggability Score (DS)</u>** |  |
| Weakly related to DrugBank/TTD target or distinct predicted biochemical activity | 0 |
| Similar to DrugBank/TTD target, no inhibitors described | 1 |
| Similar to DrugBank/TTD target, inhibitors described, but non-drug-like | 2 |
| Similar to DrugBank/TTD target, one or more drug-like compound | 3 |
| **<u>Parasite Selectivity Score (PSS) \*If applicable</u>** |  |
| Active site conserved based on 3D structure and alignment | 0 |
| Active site residues known, conserved based on sequence alignment | 1 |
| No active site information/structure available, < 70% identify by sequence alignment | 2 |
| Active site residues known, unique based on structure and alignment | 3 |
| **<u>Total Target Priority Score = RSS + ES + AS + DS + PSS</u>** |  |

After removing genes already characterized by RNAi, we included additional stringencies to generate a list of high confidence genes likely to be essential. We found that the majority (~57%) of ‘hits’ from our initial RNAi study encoded either enzymes or large catalytic complexes such as the proteasome^24^. Because enzymes are likely to provide the best rate-of-return, we narrowed our list of genes to those predicted to encode proteins with catalytic activity based on Gene Ontology terms^34,35^ (**SI Appendix, Table S1**). We also refined this list for genes with an expression greater than 10 transcripts per million (TPM) (**SI Appendix, Table S1**). Retrospective analysis from Wang et al. found such genes are more likely to result in visible RNAi phenotypes than those with a lower expression level (<10 TPM). This filtering step enabled us to identify 576 genes with significant amino acid similarity (BLAST e-value < 1e-20) to catalogued druggable targets (**SI Appendix, Table S1**)^36–38^.

Out of the initial 576 genes, we were able to amplify and generate sufficient double-stranded RNAs (dsRNAs) for 507 genes (88%) (**SI Appendix, Table S1**). To evaluate RNAi knockdown phenotypes for these genes of interest, we treated adult male and female pairs with dsRNA over the course of 30 days (**Fig. 1*A* and *B***). During these experiments (**Fig. 1*B***), we monitored worm health under *in vitro* culture conditions. Healthy parasites attach to tissue culture substrate using both their oral and ventral suckers and can move inside of the culture vessel. Substrate attachment and normal movement were used as a preliminary readout of worm viability, identifying other previously-defined visible defects by light microscopy as they arose (tissue/gut edema, tegument/head degeneration, hypercontraction, death, *etc.*)^24^.

Genes producing phenotypes following RNAi treatment were validated by confirming gene identity using DNA sequencing and knockdown specificity through designing additional dsRNAs targeting a non-overlapping gene region when possible (**SI Appendix, Table S1**). These studies yielded 63 genes (~11% hit rate) which produced fully-penetrant phenotypes affecting attachment upon knockdown. Many worms displayed phenotypes in addition to detachment defects following depletion of individual genes of interest (**SI Appendix, Fig. S1 and Table S1**). **Focused RNAi screen yields potential targets that can be grouped according to enzymatic function**

Of the 63 genes with demonstrated fitness cost upon knockdown, we found that many fell into broad categories of enzyme activity (**Fig. 1*C* and *D* and SI Appendix, Table S2**). We identified essential genes with predicted functions in the categories of GTPases, ubiquitination/proteasomal degradation, transferases (acetyl, palmitoyl, glycosyl, *etc.*), kinases and phosphatases, peptidases/proteases, ATPases, DNA/RNA regulation, and motor proteins (**Fig. 1*C* and *D***). We also found additional genes that were predicted to perform functions in other pathways that did not fall into any of these categories (GPCR, mutase, dehydrogenase, lipase, *etc.*). This data reaffirms many of the trends that we observed in our previous work^24^. Particularly, that parasites are especially sensitive to knockdown of genes encoding proteins involved in proteostasis and RNAi targeting large protein complexes like the proteasome are likely to produce a phenotype. The V-type (vacuolar) ATP-dependent proton channel exemplifies another essential large protein complex represented in our data set, the knockdown of numerous individual subunits producing phenotypes following RNAi.

Two of the most abundant categories that produced severe phenotypes were GTPases and kinases, the former commonly leading to rapid death in adult parasites upon knockdown (**SI Appendix, Fig. S1**). GTPases present as molecular switches for numerous key cellular processes^39,40^, and kinases are frequently involved in vital signaling cascades^41,42^. Thus, it is understandable that depletion of these genes would result in deleterious effects. In addition, there were several proteases and peptidases whose knockdown produced phenotypes in parasites. Peptidases have often been proposed as potential drug targets for the development of therapeutics for schistosomiasis because of their involvement in critical cellular processes^43^, such as the proteolytic invasion machinery of cercariae that enable them to infect their mammalian hosts^44–46^. Another example are the cathepsins^47–51^, a family of cysteine proteases (B1, L1/F, L2, L3, C, and D) that are secreted into the schistosome gut and aid in the digestion of hemoglobin and its subsequent metabolism. Confirming the necessity of these enzymes for normal parasite function, we identified a putative cathepsin F ortholog (Smp_034410) that was essential for parasite survival *in vitro* (**SI Appendix, Table S1 and Fig. S1**). We also identified proteases that function as key regulators in cell signaling and developmental programs^52–58^. A putative ADAM17 ortholog was found to be essential for parasite survival (**SI Appendix, Table S1 and Fig. S1**). This metalloenzyme is expressed ubiquitously in mammals and contributes to important physiological processes^59^, such as the processing of tumor necrosis factor (TNF)-α, a critical factor in inflammation^60–62^.

### *In silico* prioritization of potential drug targets

Our previous study identified 181 genes producing a phenotype upon knockdown^24^, and the experiments performed here revealed an additional 63 (**Fig. 1*C* and *D* and SI Appendix, Table S1 and Fig. S1**). However, not all essential genes are equally targetable with drugs. Thus, it is unlikely the 244 potential targets from our RNAi studies are equally suited for the development of therapeutics. Our goal is not only to identify essential genes encoding potential drug targets, but also to provide data that enables us to decide which targets are best to proceed with for a ready-made drug discovery campaign. Therefore, we have decided upon a set of unbiased criteria that represents characteristics for an ideal drug target, thereby enabling us to distinguish amongst potential candidates. We selected prioritization criteria based on the rationale of identifying targets whose cellular processes are essential to the survival of parasites, provide an ample therapeutic window for the safe and effective treatment of disease, and have characteristics that are amenable to ready-made drug discovery campaigns. To enable us to prioritize targets for further in-depth experiments, we have assigned each target a numerical score based on the following criteria: RNAi phenotype severity, mammalian non-essentiality, assayability, druggability, and parasite selectivity (**Table 1 and Fig. 2*A***).

**Figure 2.**
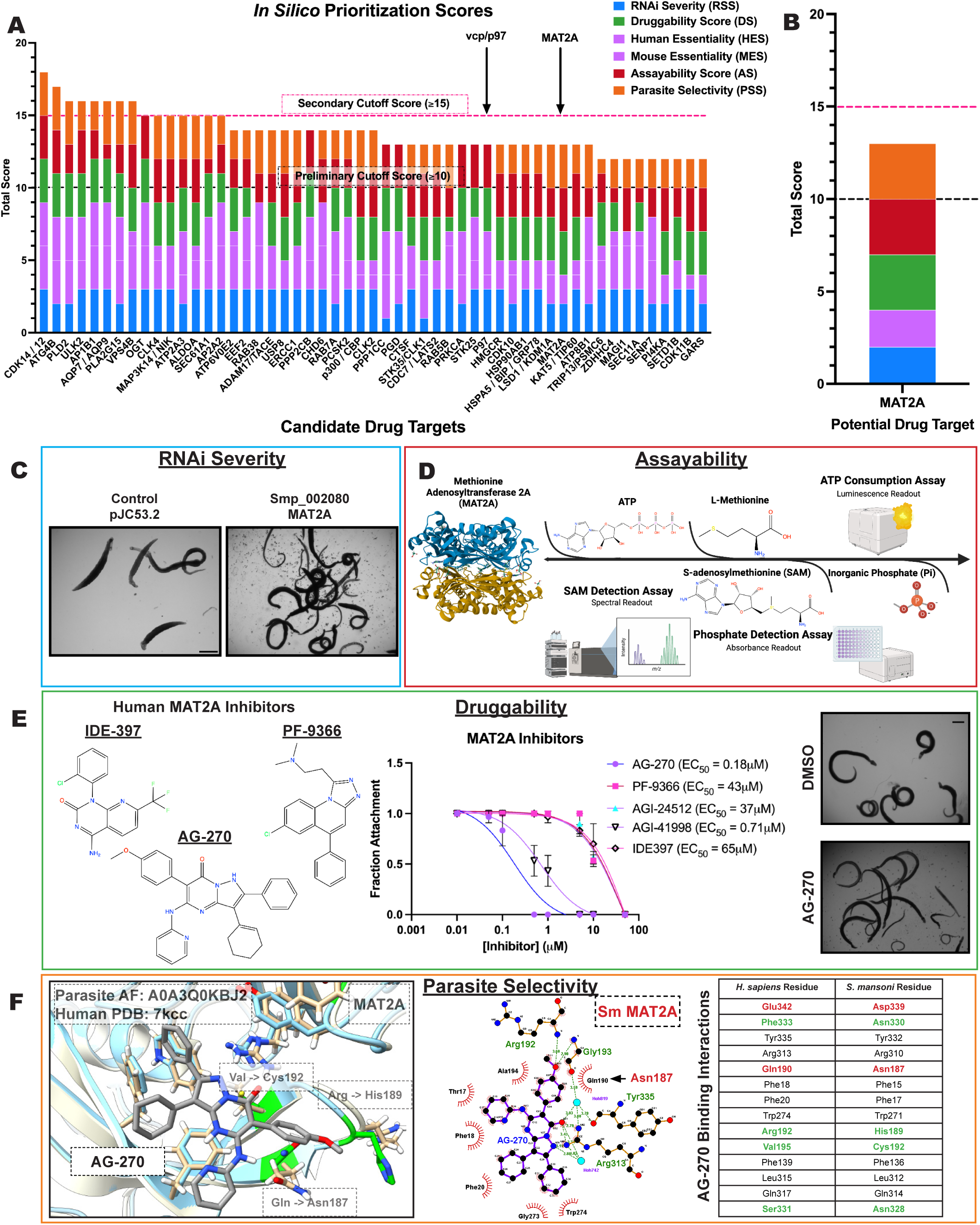
Prioritization scoring of essential schistosome genes reveals potential therapeutic targets. **(A)** *In silico* prioritization of essential genes identified by RNAi experiments. Representative targets shown were the top 53 highest scoring candidates. Potential targets were listed and prioritized according to scoring criteria detailed in Table 1. Those identifiers that possessed a score ≥ 10 in initial scoring were scored according to the criteria set in the parasite selectivity section in addition. Thresholds were set at 10 to prioritize targets with ideal characteristics in three or more categories, and ≥ 15 to represent candidates that scored well in each category. **(B)** Example of scoring for *Schistosoma mansoni* MAT2A (SmMAT2A; Smp_002080). **(C-F)** Visual representation of experimental data included in prioritization criteria. **(C)** RNAi Severity Score (RSS). Light microscopy images of adult worms treated with control dsRNA (pJC53.2) and dsRNA targeting SmMAT2A. **(D)** Assayability Score (AS). Outline of various techniques used to measure the biochemical activity of SmMAT2A (SAM Detection - RapidFire MS, ATP Consumption - Kinase Glo, Inorganic Phosphate Detection - PiColorLock). **(E)** Druggability Score (DS). Structures of known *Homo sapiens* MAT2A (HsMAT2A) allosteric inhibitors AG-270, PF-9366, and IDE-397. Light microscopy images of adult worms treated with DMSO control or HsMAT2A inhibitor AG-270 at 10 μM. Dose-response curve of adult parasites treated with different HsMAT2A inhibitors (AG-270, PF-9366, IDE397, AGI-41998, and AGI-24512). Compounds were tested from a range of 50 μM to 10 nM to determine EC_50_. Values were determined by Prism. **(F)** Parasite Selectivity Score (PSS). Alphafold structure of SmMAT2A (Blue; AF: A0A3Q0KBJ2) aligned with HsMAT2A (White; PDB: 7KCC) bound to a known inhibitor, AG-270. Unique schistosome residues are outlined in green. LigPlot schematic showing HsMAT2A (PDB: 7KCC) bound to AG-270. Tale depicting residues predicted to be involved in binding to AG-270. Conserved amino acid residues involved in binding are labeled black, while residues that are significantly different between the schistosome and human protein are colored green. Minor residues changes, such as small changes in size, are colored red. Scale bar (**C and E**), 1,000 μm.

We reason that an ideal drug target is one whose knockdown leads to rapid and severe effects within the parasite, so the worms are completely debilitated when modulated by a drug and subsequently cleared by the host (RNAi Severity Score). To allow for a wide therapeutic window, we have chosen to prioritize targets whose homologs are non-essential in mammals (human and mice) (Essentiality Score). An ideal target should also encode a soluble protein that can be expressed and purified in a recombinant system in sufficient quantity and whose enzymatic activity can be measured reliably in high-throughput format using an easily adaptable, robust, commercially-available assay (Assayability Score). Importantly, valuable drug targets need to bind drug-like molecules. Our prioritized drug targets should be closely related to human orthologs that possess commercially-available small molecule modulators with drug-like properties^63–67^ that show on-target activity *in cellulo* (Druggability Score). High-value targets should also have amino acid differences in critical binding regions, such as active sites, compared to their human counterpart to enable the development of species-selective drugs. This analysis was based on sequence and 3D structural alignments (Parasite Selectivity Score). One final consideration is that an ideal drug target should have a target-specific biological outcome that can be measured using well-established, robust methods to facilitate demonstration of on-target activity by a small molecule.

### Unbiased categorization reveals several attractive targets for the development of therapeutics

Using these metrics (**Table 1**), we have analyzed a total of 244 genes from both our initial RNAi screen^24^, as well as those uncovered in the present work (**Fig. 2*A* and SI Appendix, Table S3**). Based on these criteria, we found that there were many potential drug targets that possessed ideal characteristics in most of these categories. Conversely, we found a clear cut-off at a cumulative score of 10 in prerequisite categories (RNAi severity, mammalian essentiality, assayability, and druggability) (**Fig. 2*A***). Genes that fell below this threshold lacked ideal characteristics from two or more of these criteria. This does not mean that these are not valid drug targets, only that there are clear barriers in the path to drug discovery, whether that is a need for extensive purification or assay optimization, a lack of clear drug-like small molecule binding characteristics, or a limited therapeutic window.

Because of the intensive nature of structural studies, we utilized this cut-off score of ≥10 in the other composite categories to represent the targets with the most favorable properties for drug discovery. Using this cut-off, 85 potential targets qualified for additional investigation of selectivity using structural analysis (**SI Appendix, Table S3**). However, as a number of these potential targets encoded individual subunits of singular complexes, we combined these targets into a single representative, reducing the final candidate pool to 65. Utilizing resources such as the Protein Data Bank (PDB)^68,69^, AlphaFold^70^ and Clustal^71,72^, we have identified 51 potential drug targets that present favorable structural data (PSS ≥2) towards the development of selective drug-like molecules (**SI Appendix, Table S3**), supplying structural and unique residue differences between the parasite and orthologous mammalian structures (**SI Appendix, Fig. S3**) that can be leveraged for drug development.

During our analysis, we uncovered numerous reported small-molecule compounds that act upon the predicted human orthologs of schistosome proteins of interest (**SI Appendix, Table S3 and Table S4**). Most of them produced a phenotype in the worms at an initial concentration of 10 μM (**SI Appendix, Fig. S2*A* and *B***), but there were interesting trends that these studies revealed to prioritize future targets of interest. First, there were inhibitors that killed parasites quickly (within 24h), such as ULK-101, which targets ULK1/2 (**SI Appendix, Fig. S2*A* and *B***). The current standard of care, PZQ, paralyzes parasites within minutes following application. For a potential alternative therapy to compete with PZQ in a clinical setting, it is important that a drug act quickly, in a single dose. Therefore, drugs that display fast-killing kinetics on their cognate target are of great therapeutic interest^73^. Second, there were targets whose inhibitors were potent at sub-micromolar concentrations (**SI Appendix, Fig. S2*C***). Each of MEK1, ULK2, USP8, and MAT2A possessed commercial inhibitors that caused detachment of adult worms in culture in the nanomolar range (**Fig. 2*E* and SI Appendix, Fig. S2*C***). It is sufficient to note that the most attractive targets for further drug discovery are those whose related compounds engage their target with high potency. Lastly, it was often the case that several classes of inhibitors were available for a given target. This enabled us to test different chemical scaffolds, or perhaps binding sites, for a single target and compare potencies on worms to what is known in literature. There were several targets, such as MEK1, where one known compound (Trametinib) was profoundly effective (EC_50_ = 650 nM), yet the other (Mirdametinib) was ineffective (EC_50_ > 50 μM) (**SI Appendix, Fig. S2*C***). This suggests that there might be non-trivial differences in how these small molecules interact with their target in parasites compared to humans (*e.g.* permeability or target engagement). However, additional studies are needed to determine if these compounds are engaging the predicted target by investigating potential downstream biological outcomes following inhibition of each of these targets (i.e. substrate phosphorylation, detection of metabolite abundance, *etc.*).

This workflow led to prioritization of 18 schistosome protein targets (**SI Appendix, Table S5**) that we believe are the most promising to pursue in a drug discovery campaign. These final candidates represented ideal characteristics of our outlined criteria with the added benefit of a clear method to measure the biological outcome of pharmacologic modulation of the target by an inhibitor to determine on-target action. For instance, one hit, a schistosome ortholog of the methionine adenosyl transferase MAT2A (Smp_002080) embodied many of the ideal characteristics laid out in our prioritization criteria (**Fig. 2*B***). MAT2A is an essential metabolic enzyme that catalyzes the synthesis of the primary methyl donor in cells, S-adenosylmethionine (SAM), from ATP and methionine^74^. RNAi of the gene encoding schistosome MAT2A resulted in rapid detachment of parasites after roughly two weeks of dsRNA treatment (**Fig. 2*C***). Although MAT2A is essential in mice and various human cell lines, our own structural investigation revealed that this enzyme has an allosteric binding site that is targeted by many drugs and possesses residues distinct from its human counterpart (**Fig. 2*F***)^75^.

Allosteric inhibitors of MAT2A (**Fig. 2*E***), such as PF-9366^76^, IDE-397^77^, and AG-270^74,75^, bind at the interface between two MAT2A monomers. There are integral residues that are conserved within this binding pocket, which likely explains how we still see activity on worms (**Fig. 2*F***). But there are major changes within the surrounding pocket of AG-270, such as HsPhe333 to SmAsn330, HsArg192 to SmHis189, and HsSer331 to SmAsn328. In addition, a change from HsVal195 to SmCys192 creates an opportunity to target this allosteric binding site with covalent warheads, which would confer significant specificity to the schistosome protein over its human counterpart.

MAT2A is easily expressed in recombinant systems (*E. coli*) in soluble fractions with high yield and it catalyzes a simple biochemical reaction to convert ATP and L-methionine into SAM.^78^ This reaction can be monitored using numerous robust methods that are amenable to high-throughput format, whether it is ATP consumption (Kinase-Glo), inorganic phosphate detection (PiColorLock), or SAM formation (RapidFire) (**Fig. 2*D***). Lastly, the biological outcome of MAT2A inhibition within the context of the parasite can be easily measured through the detection of the common metabolite SAM using liquid chromatography followed by mass spectrometry (LC-MS). There is added credence to MAT2A as a ‘bona fide’ drug target, as inhibitors of MAT2A are being investigated in Phase I clinical trials for tumors with loss of the gene MTAP, constituting 15% of cancers^75^.

### p97 is a potential drug target for the treatment of schistosomiasis

As a proof-of-principle in the utility of our prioritization scheme, we conducted further studies with a schistosome protein encoding a putative valosin-containing protein/p97 (vcp/p97/cdc48) ortholog (Smp_018240). p97 is an abundant AAA-ATPase that serves a major part in the regulation of protein quality control, including mediating the degradation of ubiquitinated proteins via the endoplasmic reticulum associated degradation (ERAD) pathway^79,80^. Our lab previously described the rapid and severe phenotypes arising following p97 RNAi and the potent effects of inhibitors optimized for human p97 on adult parasites^24^. Treatment with CB-5083 (**Fig. 3*A* and *B* and SI Appendix, Fig. S4*A***)^79,81–83^, an ATP-competitive inhibitor, and NMS-873 (**Fig. 3*A* and *B* and SI Appendix, Fig. S4*A***)^82,84^, an allosteric inhibitor, led to parasite death *in vitro* at sub-micromolar concentrations^24^. However, there have been numerous inhibitors developed to target the human enzyme (**Fig. 3*A***)^85^ that remain untested on schistosomes, such as an additional allosteric inhibitor, UPCDC-30245^86–88^, and a physiological inhibitor, Eeyarestatin I (EerI)^89,90^. We revisited these compounds and tested them on adult parasites, finding that they were all effective, and produced similar physical phenotypes (**Fig. 3*A* and *B* and SI Appendix, Fig. S4*A***). We assessed on-target action as described previously^24^, performing a western blot on drug-treated worm lysate using an antibody that recognizes Lys48 (K48) polyubiquitinated proteins. Like before, we observed an accumulation of proteins targeted for proteasome-mediated degradation for known controls CB-5083 and NMS-873 as expected (**Fig. 3*C* and SI Appendix, Fig. S4*B***), meaning the function of p97 has been perturbed following drug treatment. However, we did not observe the same accumulation for CB-5083 analogs, DbeQ and ML240, despite seeing activity on the worms. ML240 killed parasites quickly, which could circumvent detectable ubiquitin accumulation. But this suggests that there might be another target in schistosomes for these inhibitors that could lead to parasite death, as is known to be true for human cells^91^ and other organisms^92^. In addition, we did not see significant increases in K48 abundance with UPCDC-30245 or Eer1 (**Fig. 3*C* and SI Appendix, Fig. S4*B***). p97 governs diverse pathways separate from protein degradation via the ubiquitin proteasome system^28,93^. Certainly, UPCDC-30245 has been shown to have a unique mechanism of action in blocking endo-lysosomal degradation^94,95^ and exerts only weak effects on protein ubiquitination.

**Figure 3.**
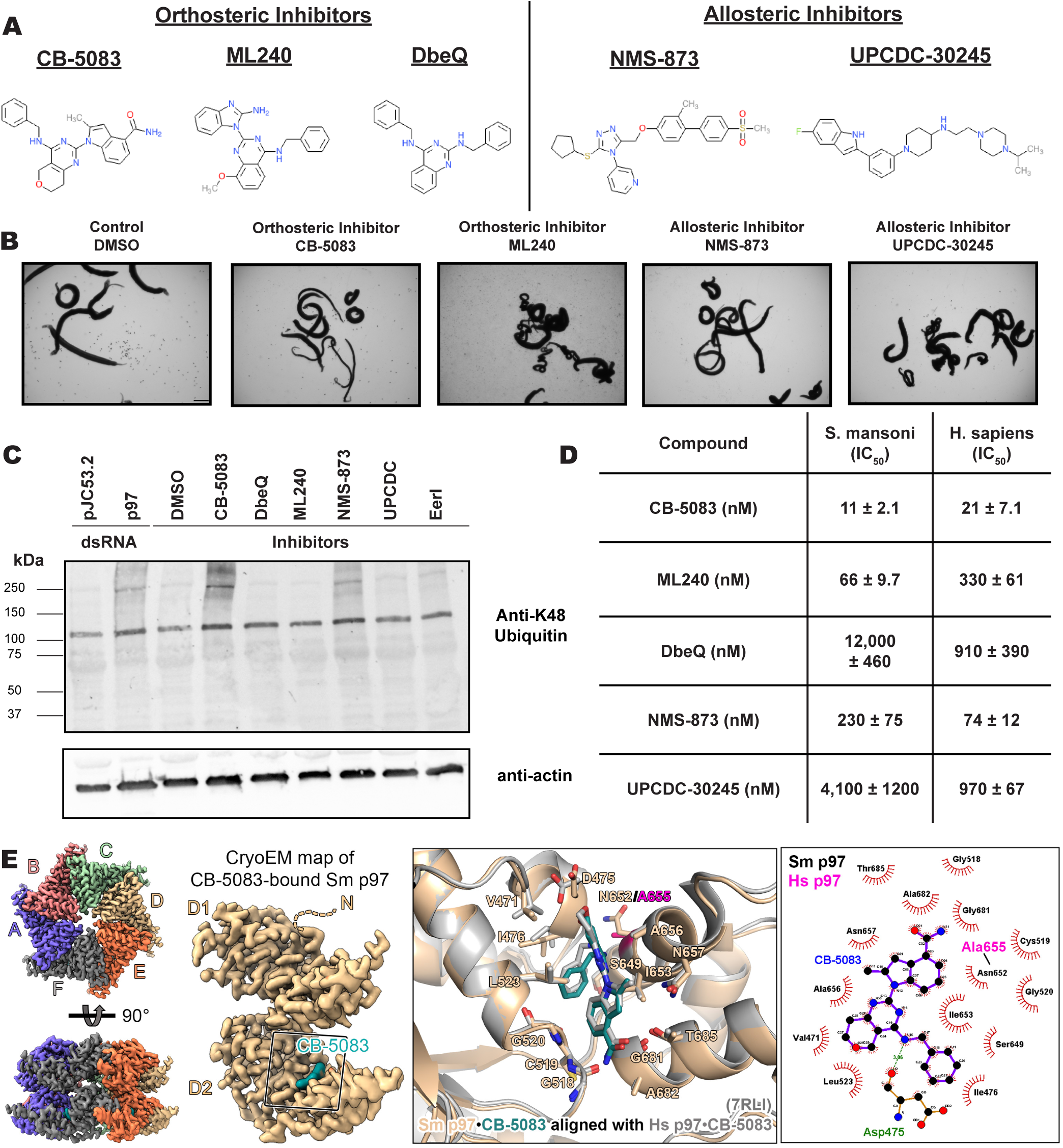
A parasite vcp/p97 homolog is essential for parasite survival and are affected by human inhibitors. **(A)** Chemical structures of established human p97 inhibitors classified as either active site (orthosteric) inhibitors (CB-5083, ML240, and DbeQ) or allosteric binding site inhibitors (NMS-873 or UPCDC-30245). **(B)** Light microscopy images of adult parasites treated with DMSO control or human p97 inhibitors at 10 μM. **(C)** Western blot depicting polyubiquitinated protein profile (K48 antibody) in worm lysate following treatment by dsRNA targeting p97 (pJC53.2 control) or established human p97 inhibitors (DMSO control). **(D)** Compared IC_50_ values of p97 inhibitors tested from 100 μM – 1 nM for *S. mansoni* and *H. sapiens* p97 recombinant enzyme using Kinase-Glo. Values were calculated in Prism. **(E)** Cryo-EM map and structure of the schistosome p97 hexamer in complex with the active site inhibitor CB-5083. Comparison of CB-5083 binding between the schistosome enzyme (Smp97; wheat) and human enzyme (Hsp97; grey, PDB: 7RLI). Ligplots were generated using LigPlot+ v2.2 using the structure of *S. mansoni* CB-5083-bound p97. Residues forming hydrophobic interactions are colored black, while other interactions are depicted in green. Unique human residues are colored pink. Scale bar **(B)**, 1,000 μm.

Since *S. mansoni* (804 amino acids) and *H. sapiens* (806 amino acids) p97 possess 82% identity in their amino acid sequence, we wanted to determine if there were any empirical differences in selectivity when treated with inhibitors. We purified both full-length *S. mansoni* and *H. sapiens* p97, then measured the IC_50_ of described inhibitors (**Fig. 3*A* and *D***) using a luminescence-based biochemical assay, Kinase-Glo, which measures ATP consumption. We determined that, for inhibitors related to the scaffold of CB-5083, there was some preference for the schistosome enzyme (**Fig. 3*D***). This selectivity was minor for CB-5083 (~2-fold), but more pronounced for ML240 (~5-fold). However, we were hesitant to repurpose this set of compounds for treatment of schistosomiasis, as treatment with CB-5083 in Phase I clinical trials for cancer led to undesired, off-target effects^91,96^. Unfortunately, both allosteric inhibitors, UPCDC-30245 and NMS-873 compounds were selective in favor of the human enzyme (**Fig. 3*D***). To further our ability to conduct target-based drug discovery, we obtained a structure of *S. mansoni* p97 using cryo-EM (**Fig. 3*E* and SI Appendix, Fig. S5 and Table S6**). Overall, we found minor changes in the active site of schistosome p97 compared to its human homolog. The main difference being an asparagine (Asn652) in the place of an alanine (Ala655) in the human enzyme that could contribute to selectivity (**Fig. 3*E***). To ascertain if there were any subtle differences in the conformational changes of schistosome p97 compared to its human ortholog, we determined the structure of the *S. mansoni* p97 apo-enzyme (**SI Appendix, Fig. S6*A*, Fig. S7*A*, and Table S6)** and its complex bound to ATPγS, a slow-hydrolyzing analog of ATP (**SI Appendix, Fig. S6*B* and Fig. S7*B***). We found that the binding modes and conformational changes observed in *S. mansoni* p97 following binding to active site inhibitors (CB-5083) (**Fig. 3*E***) and nucleotide substrates (ATPγS) were similar to the human enzyme^87,97^ (**SI Appendix, Fig. S7*A-C* and Fig. S8*A* and *B***). Indeed, there is a conserved mechanism in a pivot-like movement of the D2 domain and subsequent central pore constriction (~8 Å) following ATPγS nucleotide binding^87^ (**SI Appendix, Fig. S8*A* and *B***).

### High-throughput screen of *S. mansoni* p97 uncovers benzoxazole propiolamide covalent inhibitor scaffold

These findings inspired us to conduct a high-throughput screen of *S. mansoni* p97. This would provide the best opportunity to identify new scaffolds for enzyme inhibition that are specific for the parasite p97, rather than re-engineering existing compounds that are toxic or display no selectivity. So, we conducted a high-throughput screen of roughly 350,000 compounds (**SI Appendix, Table S7**). We were intrigued by one scaffold, as many compounds bearing a similar scaffold were included in the screen, but only four of the series showed marked inhibition of *S. mansoni* p97 function (**Fig. 4*A***). Each active compound possessed the same benzoxazole propiolamide moiety, while the corresponding alkenyl (Z - 324 and E - 135), or alkyl amide (326) were inactive (**Fig. 4*A***). We validated the activity of these compounds on the enzyme by performing dose-response analysis and found ~10-fold selectivity for the parasite enzyme over its human counterpart (**Fig. 4*A***). After determining their potency, we treated adult worms with these inhibitors, and three of the four parent compounds were found to cause at least detachment in the parasites after three days of treatment with 10 μM compound (**Fig. 4*B* and SI Appendix, Fig. S9*A***). Two compounds, 242 and 243, led to death in adult parasites (**Fig. 4*B* and SI Appendix, 2.** S9*A*), 243 being the more potent of the two (**SI Appendix, Fig. 9*C***). However, we did not see K48 polyubiquitinated protein accumulation by western blot following 48 hours of treatment with these initial inhibitors (**SI Appendix, Fig. S9*B***), which is expected for the majority of described p97 inhibitors.

**Figure 4.**
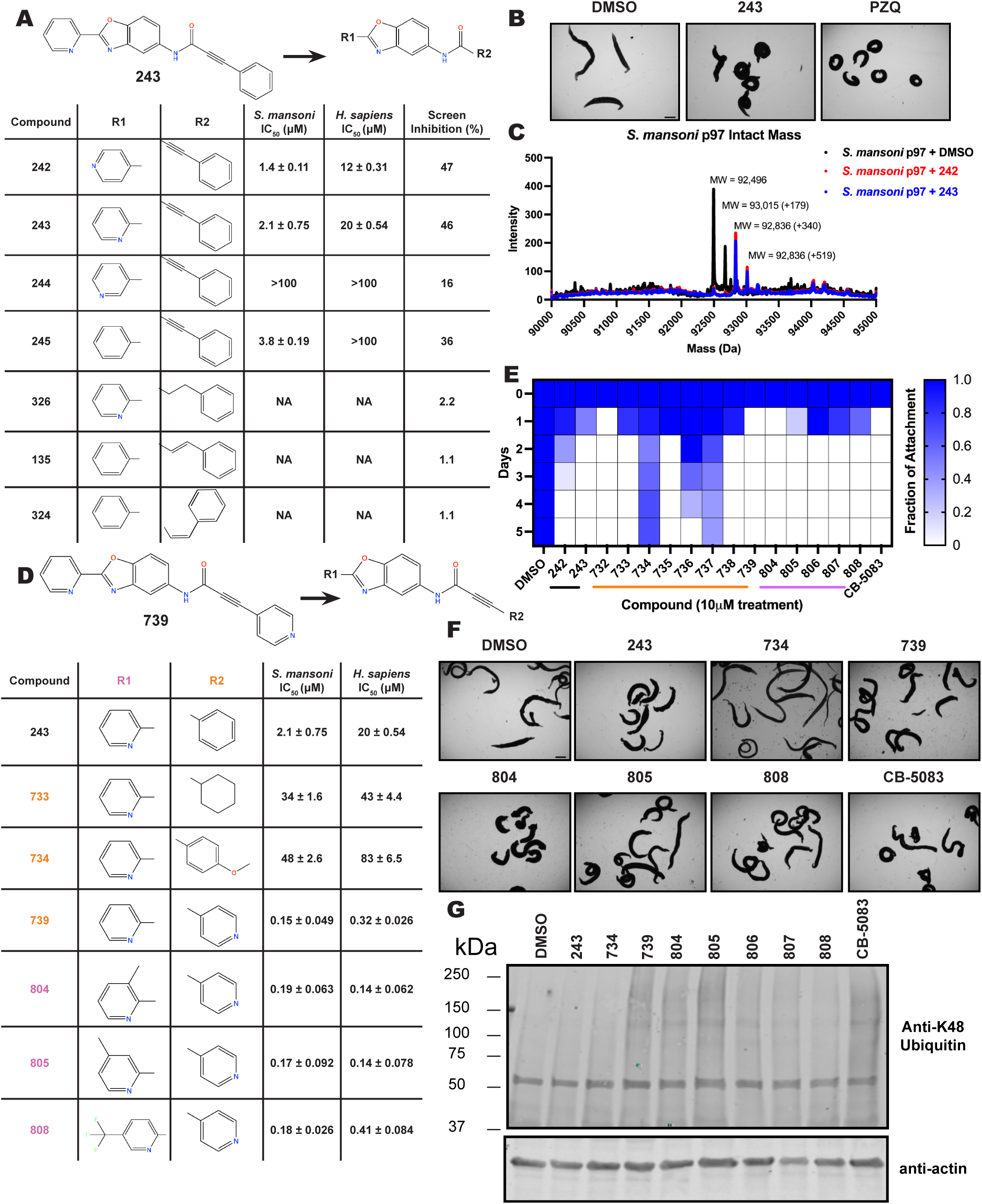
A covalent scaffold identified in high-throughput screen for inhibitors of schistosome p97 yields drug candidates that kill worms. **(A)** Chemical structure of schistosome p97 inhibitor benzoxazole propiolamide analog 243 and its related molecules identified from a high-throughput screen of the UT Southwestern 350k library. Structural activity relationship between analogous active and inactive compounds included in the high-throughput screen. Compounds were re-tested to determine the potency against parasite (Smp97) and human (Hsp97) enzyme and confirm selectivity. A value of > 100 signifies that the IC_50_ value was predicted to be greater than 100 μM or the maximal inhibition was not achieved by concentrations included in dose-response experiments (100 μM maximum). NA signifies that these compounds did not produce any notable effect on the enzyme at the tested concentrations (100 μM-1 nM). Values were calculated using Prism. **(B)** Light microscopy images of worms treated with compound 243 compared to DMSO (negative) and praziquantel (positive) control. **(C)** Mass spectrum of recombinant *S. mansoni* p97 in solution with DMSO control (black) or covalent scaffold compounds (242 - red and 243 - blue). Peak migrations account for a single p97 monomer (~92.5 kDa) and a mass shift following incubation with either covalent inhibitor (339.35 Da) or DMSO. Additional species are detected at 179 Da for either peak. **(D)** Chemical structure of compound 739. Structural activity relationship outlining **R1** (800 series) and **R2** (700 series) modifications made to the depicted scaffold. Comparative IC_50_ values display the potency of each compound on the recombinant parasite and human enzyme (100 μM −1 nM). Values were calculated using Prism. **(E)** Heat map showing attachment of a population of 10 adult worms in a culture well over time following treatment with compounds for 72 hours, refreshing drug and media every 24 hours, then monitoring for phenotypes until the end of the experiment at D5. A value of 1 (dark blue) means that all worms were attached to tissue culture substrate, and a value of 0 (white) denotes that all worms were detached from the plate. **(F)** Light microscopy images of worms treated with inhibitors from this chemical series compared to DMSO negative control and positive control CB-5083, a known p97 inhibitor that is active on adult schistosomes. **(G)** Western blot depicting polyubiquitinated protein profile (K48 antibody) in worm lysate following treatment by DMSO control or p97 covalent inhibitor analogs (actin loading control). Scale bar (**B and F**), 1,000 μm.

As the electrophilic propiolamide group is an essential part of the pharmacophore for p97 inhibition, we hypothesized that these molecules were covalently modifying p97. We confirmed this suspicion by performing intact mass spectrometry and found a shift in abundance at the expected sizes of these molecules (~339 Da) (**Fig. 4*C* and SI Appendix, Fig. S10*A* and *B***). A trypsin digest followed by mass spectrometry revealed that the dominant species modified by this scaffold was a peptide ([K].GVLFYGPPG**<u>C</u>**GK.[T]) containing Cys519, which resides within the ATP binding pocket of p97 (**SI Appendix, Fig. 10*C* and Fig. S7*D***). However, other cysteines along p97’s structure do appear to be modified by this chemical series, albeit at much lower levels (>100-fold). Covalent modification of this residue would explain the inhibition of p97’s ATPase function, as other described inhibitors that modify this residue are effective inhibitors of p97’s capacity for ATP hydrolysis^82^.

### Structure-activity relationship analysis generates potent inhibitors of schistosome p97

We sought to improve the potency of these compounds and conducted a limited structure-activity relationship (SAR) analysis on this chemical series (**Fig. 4*D* and SI Appendix, Fig. S9*C***). There were two primary sites where modifications were made to this parent scaffold. The first modification site is the benzene ring proximal (**R2**) to the propiolamide moiety, and the second is the pyridine ring distal to the same propiolamide moiety (**R1**). Compounds belonging to the 700 series (700s) contain alterations made to the **R2** chemical group, while 800 series (800s) compounds possess **R1** modifications.

We first tested the 700 series of compounds, containing modifications to the **R2** benzene ring. We found that adding electron withdrawing groups, such as a chloride, or an electron donating group, such as a methoxy, to the proximal phenyl group (**R2**) reduced potency on the enzyme (**Fig. 4*D* and SI Appendix, Fig. S9*C***). Swapping the **R2** benzene ring with another non-aromatic cyclic group, like a cyclopropane or cyclohexane, also reduced potency. Altering this benzene ring into a pyridine increased potency, giving us our most potent enzyme inhibitor, 739 (IC_50_ = 150 nM). Then, we tested the 800 series of compounds, determining how modifications to the **R1** pyridine ring affected the potency of these compounds on the enzyme. Adding a hydrophobic methyl group and moving it around the **R1** aromatic pyridine ring had little impact on potency regardless of its position. However, each of these later **R1**-modified compounds displayed sub-micromolar potency on the enzyme and produced the desired pharmacologic effects by western blot after treating adult parasites with the inhibitors (**Fig. 4*G* and SI Appendix, Fig. S9*B* and *C***).

Promisingly, the SAR trends mimicked the potency of these compounds on worms (**Fig. 4*D*-*F* and SI Appendix, Fig. S9*A* and *C***). We treated adult worms with each of these compounds at 10 μM to determine preliminary activity and tracked their attachment to tissue culture substrate as well as their physical morphologies (**Fig. 4*E* and *F* and SI Appendix, Fig. S9*A***). From the parent scaffold, only compounds 242 and 243 showed significant activity on worms, causing detachment within 48 hours in culture, ultimately leading to their death. While compound 245 did cause detachment, it did not lead to parasite death. Compound 244, which presented minimal activity on the enzyme, did not have any adverse effects on worm health and survival.

Modifications to the **R2** group (700 series) had some variable effects on parasites. Several alterations to the scaffold that decreased potency on the enzyme (734, 736, and 737) also led to decreased activity on the worms (**Fig. 4*E* and SI Appendix, Fig. S9*A* and *C***). And the most potent compounds on the enzyme, 738 and 739, had the most potent effects on worms (**SI Appendix, Fig. S9*C***), ultimately leading to death of the parasites following detachment within 24 hours. However, only compound 739 showed on-target activity by western blot, producing the predicted accumulation of K48 polyubiquitin moieties following inhibitor treatment (**Fig. 4*G* and SI Appendix, Fig. S9*B***). Some compounds of this **R2** modification chemical series (732, 733, and 735) were still active on worms despite abolishing activity on the enzyme. These off-target effects suggest that there might be another molecular target of these compounds, as is true for CB-5083^91,96^, or pharmacokinetic properties of the compounds (i.e. solubility) might obfuscate the activity seen on parasites.

Each of the compounds with modifications to the **R1** group (800 series) displayed sub-micromolar potency on the enzyme, produced an on-target biological response in the accumulation of K48 polyubiquitin chains, and killed parasites (**Fig. 4*D*-*G* and SI Appendix, Fig. S9**). With these later 800 series compounds, we found that there was a strong correlation between potency on the enzyme *in vitro* with an impact on the worms, supporting an on-target killing mechanism (**Fig. 4*E* and *F* and SI Appendix, Fig. S9*A* and *C***).

We measured the IC_50_ of each of these covalent inhibitor analogs on the human p97 at the same time as the schistosome enzyme. Our data suggests that, as the potency of these compounds increased for the parasite p97, the selectivity for the parasite enzyme over its human counterpart decreased (**Fig. 4*D* and SI Appendix, Fig. S9*C***). Unfortunately for these covalent inhibitors that appear to have an on-target killing mechanism (800 series and compound 739), there is little to no selectivity for the parasite enzyme (**SI Appendix, Fig. S9*C***). After seeing these changes to selectivity, we tested these compounds on human cells to determine cytotoxicity (**SI Appendix, Fig. S11*A-C***). We treated HepG2 cells with the parent compound (243), the most potent compound on the enzyme (739), and each of the 800 series compounds to determine cytotoxicity, using CB-5083 as a control. Each of these compounds were cytotoxic in roughly equal concentrations to the potency (EC_50_) of the compounds on worms (**SI Appendix, Fig. S11*E***). However, they were noticeably less cytotoxic to cells than CB-5083.

### Cryo-EM structure of *S. mansoni* p97 bound to covalent inhibitors reveals conformational change

To better understand where these compounds bind p97 in the hopes of improving selectivity, we sought to solve the structure of this scaffold bound to *S. mansoni* p97 using single particle cryo-EM analysis. Two cryo-EM datasets were collected for benzoxazole propiolamide inhibitor analogs 739- and 804-bound to schistosome p97 (**Fig. 5*A* and SI Appendix, Fig. S12*A* and *B***, **Fig. S6*C* and *D***, **and Table S6**). We identified density belonging to these inhibitors at an interface between two monomers of p97 (**Fig. 5*A* and *B* and SI Appendix, Fig. S6*C* and *D* and Fig. S12*A* and *B***). As suggested by our mass spectrometry experiments, Cys519 appears to be covalently linked to these molecules (**Fig. 5*A***). Presumably, this occurs through a Michael reaction conducted by the thiol of Cys519, as the entire mass of the inhibitor can be detected by intact mass spectrometry (**Fig. 4*C* and SI Appendix, Fig. S10*A***). This suggests that there is no leaving group in this chemical reaction and instead forms a carbon-carbon double bond in addition to the covalent linkage to Cys519 (**SI Appendix, Fig. S12*E***). Based on our best approximation, 739 and 804 appear to engage in slightly different binding contacts (**Fig. 5*A* and SI Appendix, Fig. S12*C* and *D***), even though they only differ by a single methyl group on the distal (**R1)** pyridine ring. For instance, compound 804 makes more putative contacts with residues in its conjugated p97 monomer (**chain D**). Additionally, compound 739 lays flat within its binding pocket, while compound 804 is rotated around the carbon-carbon double bond so that the thiol of Cys519 forms an adduct on the opposite side of the amide (**Fig. 5*A* and SI Appendix, Fig. S12*C-E***). In the case of 739, it is on the same side. This means that 804 forms an E-configuration olefin, while 739 forms an elongated Z-configuration olefin following nucleophilic attack by Cys519 (**SI Appendix, Fig. S12*E***), enabling these compounds to engage different residues within their binding pocket. However, given the resolution of the maps and density for these inhibitors, we cannot rule out that these compounds may be bound in a different configuration.

**Figure 5.**
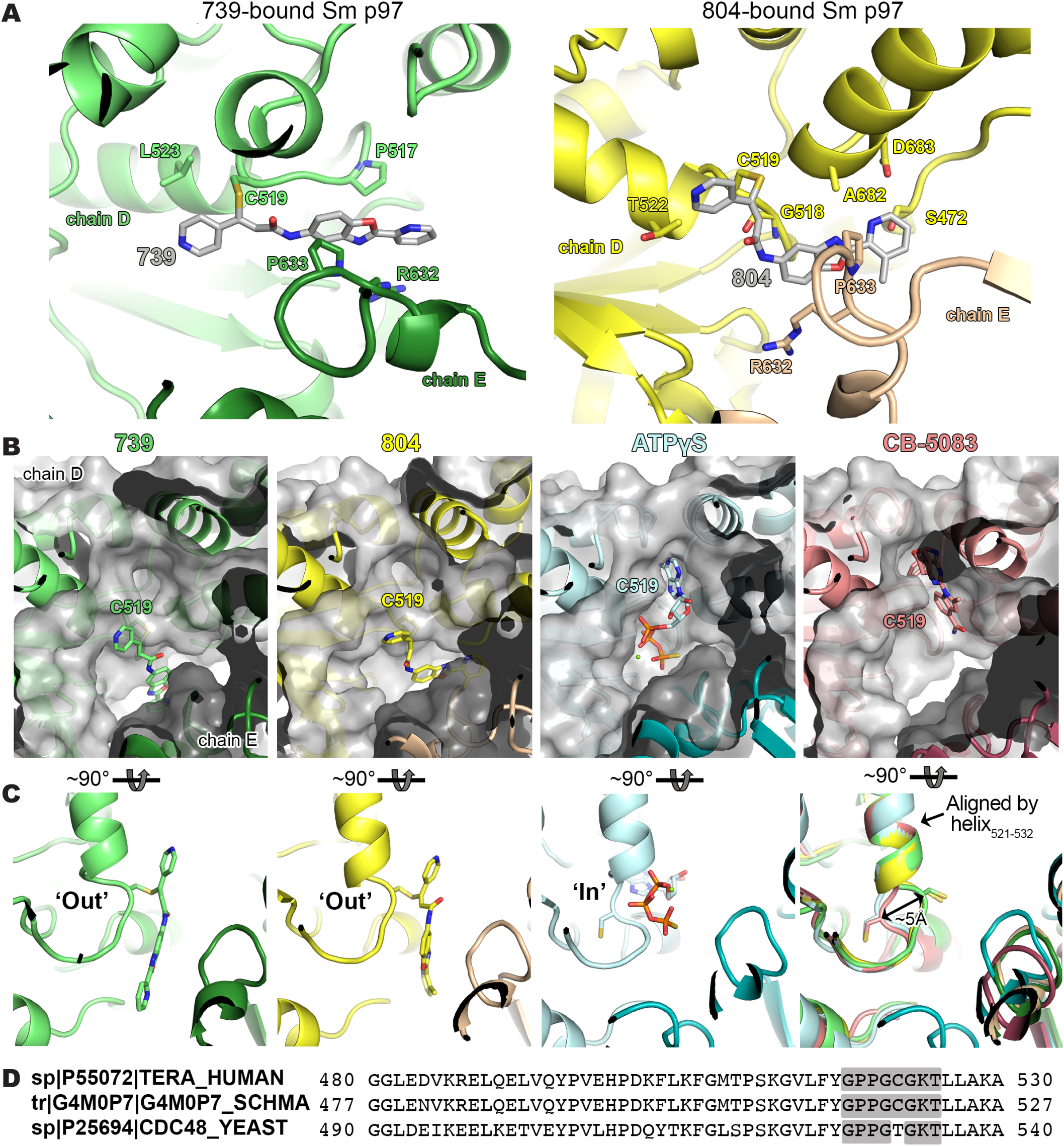
Cryo-EM Structure of *S. mansoni* p97 bound to covalent scaffold. **(A)** Binding pocket of covalent inhibitors 739 (green) and 804 (yellow) within the D2 domain of *S. mansoni* p97 and the adjacent monomer. Potential contact residues of compounds 739 and 804 are shown in sticks. **(B)** Binding pocket of covalent inhibitors 739 (green) and 804 (yellow), shown as grey surfaces, relative to known ligands, ATPγS (blue) and CB-5083 (pink). **(C)** Positioning of P-loop in the ‘In’ or ‘Out’ position within the D2 domain of the schistosome p97, aligned by the indicated alpha helix (residues 521-532). **(D)** Sequence alignment of the conserved human and schistosome P-loop within the D2 walker domain of p97 compared to a yeast homolog.

These molecules occupy an adjacent, and overlapping, binding site to CB-5083 and ATPγS (**Fig. 5*B***). As a result of covalent conjugation, the flexible loop where Cys519 resides is displaced to an outward (‘out’) position (**Fig. 5*C***), which points towards the neighboring subunit. Consistent with the apo and CB-5083-bound states, the conformation of the rest of the enzyme apart from this loop remains unchanged (**SI Appendix, Fig. S8*A* and *B***). Cys519 is positioned within a flexible “P-loop” of the Walker A motif (GxxxGKT) in the D2 domain of p97, which appears to be broadly conserved across multicellular eukaryotes, but not certain unicellular organisms such as yeast (*S. cerevisiae*)^98^(**Fig. 5*D***). Walker motifs are sequences involved in binding to nucleotides and coordinating their hydrolysis^99^. Indeed, the amino backbone of Cys519 is involved in binding ATPγS in both the human^97^ and schistosome protein (**SI Appendix, Fig. S7*B* and *D***). Cys519 is flanked by a conserved region of flexible amino acids [GPPG**<u>C</u>**G] that may enable the loop to move freely depending on its physiological context. Interestingly, Cys519 naturally occupies an inward (‘in’) position in its apo-form, when bound to nucleotide substrate (ATPγS), or active site inhibitors (CB-5083) (**Fig. 5*C***). However, when it binds to these benzoxazole propiolamide inhibitors, the Cys519 moves into an outward (‘out’) position. While the sulfhydryl sidechain of Cys519 is not utilized in the catalysis of nucleotides, the modification of this residue, and the subsequent conformational change of the loop where it resides (**Fig. 5*C***), locks the backbone of Cys519 in a position where it can no longer make the contacts required for catalysis. This conformational change would render the enzyme incapable of binding both the inhibitor and ATP.

This is a conformational change that, to the best of our knowledge, has not been seen for structures of p97 deposited in the PDB^68,69,87,94,97,100,101^. But equivalent conformational changes are well-documented for protein kinases that bind Type II inhibitors^102–105^. Kinase activation loops contain a conserved Asp-Phe-Gly (“DFG”) motif. In an active conformation, the Asp within this motif points inwardly to the ATP-binding pocket and assists in coordination of Mg^2+^ ions. In an autoinhibited state, this same Asp flips ~180° to an outward position in a shift that is ~5 Å from the ATP binding site. The shift seen from Cys519 in p97 is also roughly a 5 Å movement that forms the new allosteric pocket adjacent to the ATP binding pocket (**Fig. 5*C***). The ‘in’/’out’ switch we see in this context for p97 mimics a DFG flip in protein kinases. This represents a significant finding, where new classes of inhibitors can be developed for this allosteric binding pocket of p97 for the numerous disease states (cancer, Alzheimer’s, *etc.*) it is involved in across species.

Visualizing the binding of this compound further explained the trends we saw with SAR (**Fig. 4*D* and SI Appendix, Fig. S9*C***). The pocket where these compounds bind is narrow, so there is little room for bulky substituents (**Fig. 5*A*-*C***). These observations suggest the addition of functional groups to the proximal aromatic group (**R2**) decreased potency as this likely interfered with its fit into the pocket, clashing with sidechain density. However, the distal aromatic group (**R1**) is facing solvent, so the addition of other functional groups, even bulky ones, did not significantly reduce the potency of the scaffold. This binding site has been explored with docking experiments^84,106^ and is confirmed to be similar for our scaffold using cryo-EM. Similar to how the discovery of the ‘DFG’ flip led to the discovery of new classes of inhibitors in kinases, our covalent compounds reveal a new opportunity to develop inhibitors that target this unique allosteric pocket in p97.

## Discussion

### Schistosome p97 as a therapeutic target

p97 is a pivotal constituent of the UPS that extracts and disassembles its substrates from various cellular compartments. The discovery of p97 inhibitors has provided not only basic research tools, but an avenue for effective therapies in the clinic for cancer and neurodegenerative diseases. We conclude that there are also further opportunities for p97 as a drug target for the treatment of infectious parasitic diseases such as schistosomiasis. Despite there being significant identity between the human and schistosome enzyme (~82%), we have identified selective compounds for the schistosome enzyme over its human counterpart that kill parasites. Outside of the covalent scaffold explored in this paper, we also uncovered numerous other inhibitors of p97 through our high-throughput screening efforts. Many of these other scaffolds bear good chemistry (Lipinski’s rule of 5, Veber’s rule, etc.)^63–67^ and display on-target activity in adult worms. In addition, there are numerous binding locations where selective chemicals can be developed even if the active site is mostly conserved. Indeed, we have observed decreased potency for the allosteric inhibitors, NMS-873 and UPCDC-30245, on the schistosome enzyme *in vitro* compared to the human enzyme. This empirical evidence implies that compounds might be engineered to create compounds where is inverse is true, being potent for the schistosome enzyme rather than the human. Not only that, but we have confirmed there are additional allosteric binding sites where molecules can bind, such as the one for covalent inhibitors, 739 and 804, at the interface between two monomers of p97 described in this paper. Interestingly, elucidating the binding pocket of these benzoxazole propiolamide inhibitors has uncovered an inhibitory mechanism where the flexible nucleotide binding motif within the D2 domain of p97 can be locked in an ‘out’ position, distinct from its typical ‘in’ conformation. This provides new insights into the potential mechanistic activity of p97 in its native state and its ability to sense substrates using this flexible motif. Presumably, this conformational change mimics a physiological capacity. We hypothesize that this loop is involved in substrate sensing and ‘breathes’ in a naturally flexible state, but the predominant species is found in the ‘in’ conformation as seen with the apo enzyme or nucleotide-bound state (**Fig. 5*C***). p97 is likely constitutively bound to ATP or ADP as the natural concentrations of these substrates are high (0.5 - 5 mM) in its cellular context^112–116^. Indeed, most purified recombinant p97 has been found to be pre-bound to nucleotide that must be removed by heat or apyrase treatment^30,101,117–119^. However, our enzyme was purified in its apo state, suggesting that the allostery of nucleic acid binding may be different between the human and schistosome enzyme. Flipping into an ‘out’ conformation could be important for releasing ATP and/or ADP, or for binding to another substrate or cofactor. Alternatively, the flexibility of this motif could be important for inter-subunit communication between monomers or domains as is known to occur within p97^118,120–122^. Regardless, binding to this scaffold has revealed a unique conformational change in p97 that could uncover more biology of the hexameric ATPase and lead to further development of therapeutics that target it. While our own covalent scaffold was cytotoxic on human cells comparable to activity on adult worms (**SI Appendix, Fig. S9*C* and Fig. S11*E***), it does not mean that more selective inhibitors cannot be developed in the future that target this pocket. And it would be intriguing to investigate whether non-covalent scaffolds could also access this allosteric binding pocket. Furthermore, p97 is evolutionarily conserved across phyla, and the more removed from metazoans, the more unique differences are present in the structure of the enzyme. This leads to greater opportunities to discover selective therapeutics for the treatment of disease.

### Other potential drug targets

For the purposes of this paper, we have focused primarily on p97 and MAT2A to illustrate the benefits of the outlined prioritization criteria in taking an unbiased approach to pursuing potential schistosome drug targets. However, 65 potential targets scored well in our criteria (cumulative score ≥10), and 51 of them possessed favorable structural data (PSS ≥2) that suggests selective inhibitors could be identified or synthesized (**SI Appendix, Table S3**). Among these were drug targets in various enzymatic categories, such as USP8, a deubiquitylating enzyme, LATS2, a kinase involved in cell proliferation, ADAM17, a peptidase involved in cell signaling, and others. Among those possessing favorable structural data, there are several targets where a nucleophilic residue has replaced a more labile residue present the human ortholog. In the schistosome orthologs of PLD2 and MAT2A, cysteines were located within binding pockets that are not present for their human counterparts (**Fig. 2*F* and SI Appendix, Fig. S3*B***). These residues could not only confer selectivity, but provide a means for targeted covalent strategies to modulate activity.

Many of the potential drug targets identified from RNAi experiments, however, were found to fall below a threshold of 10 when compiling the scores from each individual category. The primary reason for this was the candidates lacked ideal characteristics from two or more of the listed categories. For example, subunits of the proteasome were listed below this threshold, thus deprioritized from future studies. However, it is well known that proteasome inhibitors are effective against many different types of disease, including parasitic infections^125^. Schistosomes are present among those that can be killed by proteasome inhibitors like bortezomib^24^. However, it is essential in human cells and difficult to recombinantly express the entire proteasome, thus it must be purified from entire worms. While the feat has been done^126^, it remains to be seen if it can be done at a scale that enables high-throughput screening of hundreds of thousands of compounds against the enzyme complex. Alternatively, a significant amount of time can be contributed to optimizing an appropriate recombinant system for purification. Therefore, the proteasome is likely a viable target, but not ideal for a ready-made drug discovery campaign.

### Final thoughts

Drug discovery is a rapidly changing landscape, where nontraditional methods of chemical intervention are revealed each year. As such, it would be inappropriate to completely dismiss a single gene encoding an essential protein from becoming a therapeutic target. It is our goal, however, to provide sound data on which potential drug targets provide the path of least resistance for a drug discovery campaign to develop alternative treatments to schistosomiasis. This paper provides a valuable resource to the community to enable the rational discovery of alternative therapeutics to the current standard of care, praziquantel.

## Methods

### Worms and culture

Adult *S. mansoni* (NMRI strain) (6-7 weeks post-infection) worms were harvested from infected female Swiss Webster mice by hepatic portal vein perfusion using 37°C DMEM (Mediatech, Manassas, VA), 8% Horse Serum, and heparin. Parasites were rinsed in DMEM + 8% Horse Serum and cultured at 37°C, 5% CO2 in Basch’s Medium^127^ and Antibiotic Antimycotic (Gibco/Life Technologies, Carlsbad, CA 92008). Experiments with and care of vertebrate animals were performed in accordance with protocols approved by the Institutional Animal Care and Use Committee (IACUC) of UT Southwestern Medical Center (approval APN: 2017-102092).

### Bioinformatic identification of potential drug targets

To identify druggable genes within schistosomes, we compiled known human drug targets from databases such as the Therapeutic Target Database (TTD)^33^, Drug-Gene Interaction Database (DGIdb)^128–130^, ChEMBL^32^, and DrugBank^31^. Drug targets were also pulled from literature reviews^131^, prioritizing kinases as potential drug targets^132,133^. Identifiers (Ensembl and Uniprot)^134,135^ were used in combination with BLASTp^36–38^ to establish schistosome genes with similarity to these drug targets (Schistosome proteome: PRJEA36577) (Human Proteome: Homo_sapiens.GRCh38). Once schistosome accessions were retrieved, we removed any IDs with an e-value > 0.0001. Then, we filtered out duplicate IDs or those that have previously been cloned by the Collins Lab^24^. Next, we removed any IDs that did not possess a predicted catalytic activity according to GO terms^34,35^. Lastly, remaining IDs were prioritized and included for screening if they had > 10 transcripts per million (TPM). Raw and processed RNA-Seq data for adult male parasites have been deposited in NCBI (**GSE290988**).

### Large-scale RNAi screen

For our screen, we designed primers to amplify ~700bp (500-1000bp) fragments using BatchPrimer3 (http://batchprimer3.bioinformatics.ucdavis.edu/index.html). If genes were shorter than 700bp, primers were designed to cover as much of the transcript as possible. To enable RNA synthesis via reverse transcription, we added a T7 promoter (GAATTTAATACGACTCACTATA) sequence to the 5’ end of each oligo. Following the T7 site, and flanking the gene-specific sequence, we inserted a NotI (GC^GGCCGC) and AscI (GG^CGCGCC) restriction enzyme site, respectively, to facilitate DNA sequencing of amplified cDNAs to validate RNAi phenotypes observed with the encoded sequence. Oligos were synthesized and prepared in 96-well format. Target genes were amplified using PCR from adult schistosome mixed sex cDNA template. 5 μL of PCR product was used for *in vitro* transcription (IVT) to generate a total volume of 100 μL dsRNA. IVT reactions proceeded at 37°C overnight, then underwent successive 3-minute annealing steps at 95°C, 75°C, and 55°C, finally cooling to room temperature until use or storage at −20°C. Successful synthesis was verified using agarose gel electrophoresis to determine presence and size of PCR products and resulting dsRNA. RNAi experiments were conducted on approximately 5 adult worm parasite pairs (or 5 adult males) in 12-well plates. Worms were cultured in 3 mL Basch 169 media and treated with 20 μL dsRNA on D0 and D2, then every 7 days afterward (D9, D16, and D23). Experiments finished on D30, where videos were captured using light microscopy (Axio Zoom V16) to document visible phenotypes that manifested during the treatment. Media was changed every 1-2 days, where worm attachment, morphological changes, and any other aberrant observations were recorded in addition.

The identity of hits from the RNAi screen were confirmed by digesting PCR products using NotI (NEB) for 30min at 37°C. DNA bands were purified from agarose gels using Zymoclean Gel DNA Recovery Kit, then sequenced using a T7 primer. For hit validation, primers of non-overlapping gene fragments were designed using BatchPrimer3. In the case of gene sequences too short to design non-overlapping primers, the original primers were used in the absence of initial modifications (T7, NotI, and AscI) added to facilitate large-scale IVT. Amplified PCR products using these genes were inserted into pJC53.2 using TA cloning. Plasmids from positive clones were purified from *E. coli*, then sent for sequencing to verify identity of the inserted constructs. Upon confirmation, these plasmids were used to generate dsRNA to repeat RNAi using the same treatment schedule as before. Hits were only considered validated if they displayed similar, fully penetrant phenotypes in three independent experiments using biologically unique batches of dsRNA and worms.

### *In silico* prioritization rationale

We prioritized potential drug targets in schistosomes based on the rationale that 1). Complete debilitation of parasites upon interfering with a target fares best for host clearance (RNAi Severity), 2). Non-essential homologs in mammals (human and mouse) provide the widest therapeutic window (Mammalian Essentiality), 3). Encoded proteins should be expressed and purified in recombinant systems in sufficient quantity to be measured reliably in high-throughput format to pursue a target-based drug discovery method (Assayability), 4). Potential targets should possess features that can bind drug-like molecules (Druggability), and 5). Sequence and 3D alignment data should supply unique amino acid differences in critical binding regions (Parasite Selectivity). Candidate essential genes were scored on a numerical scale in whole integers from 0 (unfavorable qualities) to 3 (highly favorable qualities) in each category based on how well they fit the criteria described above. Their cumulative score determined how suitable any given candidate drug target was for a ready-made drug discovery campaign. **RNAi severity**

It is reasonable that targets whose RNAi phenotypes occur rapidly and with the most deleterious effects represent the most attractive targets for the development of therapeutics. Therefore, we have prioritized targets based on the time a phenotype manifested relative to the start of the experiment, the severity, and number of phenotypes that manifested (**SI Appendix, Table S1**). Highest priority (a score of 3) was given to essential genes whose phenotypes began to manifest within the first two weeks (D1-D15) of the 30-day RNAi experiment and whose knockdown produced more than one morphological defect (tissue/gut edema, tegument/head degeneration, hypercontraction, death, etc.). Next, a score of 2 was attributed to essential genes who had more than one phenotype as listed above but occurred later in the experiment schedule (D16-25). A score of 1 was given to essential genes who had a modest phenotype (i.e. detachment only) that manifested at an intermediate point in the experiment (D16-25). A score of 0 was given to essential genes producing a modest phenotype late in the RNAi experiment (D25-D30). Attachment and morphologies were compared to negative vector controls (treated with pJC53.2 IVT product encoding the bacterial gene *CcdB*).

### Mammalian essentiality

The current standard for treatment of schistosomiasis, PZQ, poses minimal toxicity to humans, while effectively clearing hosts of parasite burden^136^. Thus, it is vital that an alternative drug provide a large therapeutic window. We reason that targets essential for schistosomes, but dispensable for mammalian (human and mouse) survival are most likely to provide this therapeutic window. Using our initial bioinformatic searches for schistosome genes with high similarity to drug targets, we have defined human and mouse orthologs using data available in Wormbase Parasite^137^ and BLAST^36–38^ (**SI Appendix, Table S3**). Then, we examined these orthologs in the Online Gene Essentiality Database^138^, which catalogs essentiality information from both mouse knockout studies and RNAi/CRISPR screening panels of human cell lines. We have given each target a score for each human and mouse essentiality, giving priority to targets whose orthologs are non-essential following mouse knockout and disruption in human cell lines. The highest possible score (3) was given to targets who are non-essential in mice and human cell lines. A score of 2 was awarded to any candidate whose homolog was essential in some human cell lines, but not others, or essential in mice, yet did not lead to embryonic lethality. If no information was available for mice and human cell lines, then a potential drug target was given a score of 1. The lowest score (0) was given to candidates whose homologs were essential in humans and mice.

### Assayability

Target-based screening is an attractive avenue for the identification of small molecules for drug discovery, especially for those targets whose functions have been thoroughly vetted within their cellular context. It provides a wealth of scaffolds that can engage the target, and is also economically feasible, as large quantities of recombinant protein can be readily purified from other organisms like *E. coli* or insect cells. We have given priority to targets bearing homology to proteins that are documented to be well-expressed recombinantly, solubly, and in their native fold. Furthermore, these ideal targets must possess a predicted biochemical activity that can be measured using a commercially available, high-throughput amenable assay and readout (absorbance, colorimetric, fluorescence, luminescence, etc.). Extensive research has been done in literature to determine if a target protein’s ortholog has been purified and if its biochemical activity has been measured using commonly available reagents and platforms. References have been provided for assays and purification schemes of human orthologs that have been identified (**SI Appendix, Table S3**). We have prioritized targets with orthologs that have historically been purified from recombinant systems in sufficient amounts to perform a large compound library screen, and whose activity can be measured using an easily adaptable, commercially available assay. The most ideal potential drug targets were awarded a score of 3 if there were established purification schemes for a given target ortholog, and a method existed to measure its biochemical activity using a robust, commercially available assay. A score of 2 was given if target purification and assay could be achieved with minimal optimization. This means that the target can be purified, and its activity assayed. However, there are additional steps that may require some optimization, but can generally be achieved with products that can be purchased commercially or protocols that exist and need to be adapted slightly. A lower score (1) was given if purification and assay could be achieved but required extensive optimization, meaning protocols need to be developed or extensively adapted, and reagents to achieve purification and assay are not available commercially, and may need to be made in-house. The lowest score (0) was given if there was no clear path to recombinant purification of the target, and no currently available way to measure its biochemical activity with amenability to high-throughput format. There are many enzymes that can be purified for which a general assay can be applied to determine if a given molecule is interacting with a protein, such as Isothermal Titration Calorimetry (ITC), Differential Scanning Fluorimetry (DSF), and Surface Plasmon Resonance (SPR). These were rarely considered because we wanted to utilize assays that measured the distinct biochemical activity of a given target protein. Likewise, virtual screening is a powerful method to identify potential ligands for enzymes where there is no simple biochemical assay to utilize for traditional high-throughput screening, yet does not factor into our ranking for potential drug targets.

### Druggability

For our purposes of identifying potential drug targets that could yield alternatives of PZQ, it is not sufficient that a gene is essential. Indeed, the gene must also encode a protein whose activity can be modulated by binding a small, drug-like molecule. Here, we define drug-like molecules as small molecules having characteristics that are similar to existing drugs according to well-known criteria such as Lipinksi’s rule of 5 and Veber’s rules (i.e. LogP < 5, molecular weight < 500 Da, < 10 hydrogen bond acceptors, < 5 hydrogen bond donors, ≤ 10 rotating bonds, < 140 Å^2^ polar surface area)^63–67^. We have already narrowed our initial search to proteins with high amino acid identity to well-documented drug targets in databases such as ChEMBL, DrugBank, and TTD^31–33^. This greatly enhances the chance of identifying a druggable schistosome protein. However, some of these targets represent “theoretical” targets that are heavily implicated in diseases but have never been the subject of drug-discovery campaigns. Others are ‘bona fide’ drug targets that have undergone drug discovery efforts, with some compounds making it into clinical trials. Therefore, we have assessed the predicted biochemical properties and function of each potential schistosome target to determine if they have features that can bind to drug-like molecules^63–67^. We have also searched these databases for existing compounds that have been developed or identified to bind human orthologs of these potential schistosome targets. Simultaneously, we have searched through chemical vendors such as Selleck Chem and MedChemExpress to find inhibitors of target protein orthologs. Targets have received priority if these compounds are available commercially, possess drug-like properties, and have shown on-target activity *in cellulo*. Where possible, we have provided references for inhibitors of target protein orthologs (**SI Appendix, Table S3**). If a potential schistosome drug target has similar properties to its human ortholog, and there are one or more one drug-like compounds catalogued in databases such as DrugBank^31^ or on commercial vendor websites such as MedChemExpress, then it was given a score of 3. A score of 2 was given if a candidate target was similar to its druggable ortholog, and there were described inhibitors, but these compounds were not drug-like in nature^63–67^. A target was awarded a score of 1 if a candidate target was similar to its druggable homolog, but there were no inhibitors described. Lastly, a potential target was given a score of 0 if it was only weakly related to a druggable target in humans.

### Parasite selectivity

As many of our potential targets are derived from the [Human] “Druggable Genome”, it is important to confer specificity where available to reduce potential toxicity. Utilizing resources such as AlphaFold^70^ and Clustal^71,72^, we have identified drug targets that supply unique differences between the parasite and mammalian ortholog. This enables us to compare sequence alignments and 3D structural information available in the Protein Data Bank (PDB)^68,69^ to determine inherent selectivity between these enzymes. We have used these databases to compare protein sequence alignments and generate homology models to determine predicted similarity of active sites and potential allosteric pockets (**SI Appendix, Fig. S3**). Higher priority was given to potential targets bearing critical amino acid differences in these positions or possessing allosteric binding sites where selective drugs can be synthesized. Because of the intensive nature of structural studies, we have limited this category to only potential targets that yielded the highest composite score in the other listed categories (**SI Appendix, Table S3**). If a gene scored ≥10 in the categories of RNAi Severity, Mammalian Essentiality, Assayability, and Druggability, they qualified for additional investigation for parasite selectivity. This cut-off afforded structural analysis for 65 essential genes.

Structural analysis was first performed by identifying binding pockets and key binding residues from data available in literature or on the Protein Data Bank (PDB)^68,69^ for human orthologs of schistosome target proteins. For sequence alignment, corresponding human and schistosome protein sequences were retrieved using UniProt^135^ and WormBase Parasite^137^, respectively. Sequences were aligned using Clustal^71,72^. Then, we looked for differences in key residues involved in substrate or drug binding interactions or the formation of binding pockets. For 3D homology modelling, we searched the PDB^68,69^ for solved structures (x-ray crystallography, cryo-EM, etc.) of the human orthologs of potential parasite target proteins. Structures were used if they were of sufficient quality or involved binding of a drug included in inhibitor testing. Here, we define sufficient quality according to wwPDB validation reporting for each PDB dataset, where good quality structures have metrics (Rfree, Clashscore, Ramachanran outliers, Sidechain outliers, RSRZ outliers) that rank as ‘better’ relative to X-ray structures of similar resolution and bearing ‘better’ quality ligand structure fit. Additionally, we retrieved theoretical models of schistosome proteins available on AlphaFold. We have provided identifiers for both AlphaFold and PDB models used in these studies (**SI Appendix, Table S3**). Homology modelling was conducted using ChimeraX^139^ and Pymol^140^, identifying residue overlap using the matchmaker and/or alignment function. In some instances, we also utilized LigPlot^141,142^ to visualize binding pockets of human orthologs of schistosome target proteins to display important binding interactions.

We awarded the highest score (3) to potential targets whose active sites and allosteric binding pockets are well-defined, and there are unique residues in either of these respective features that could be exploited for the development of selective therapeutics. The next highest score (2) was given to candidates whose active sites were not well known or had <70% identity based on sequence alignment using Clustal^71,72^. A score of 1 was given to a target whose active site residues are known and conserved but only based on sequence alignment. The lowest score (0) was given to a target whose active site residues are known and conserved based on 3D structure and sequence alignment.

### Biological outcomes

A final consideration to the amenability of a candidate target to drug discovery is whether on-target activity can be established. It is vital to reliably measure target-specific biological outcomes as a result of its modulation by a small molecule. Some biological processes are highly conserved in species and can be readily applied, such as protein degradation by the proteasome, which can be monitored with a western blot using antibodies that recognize various ubiquitin linkages (DUBs, p97, proteasome, etc.). Additional examples include the maturation of lysosomes using an LC3B antibody (ULK2, ATG4B), or histone methylation (KMT2, KDM1A, etc.). Alternatively, other targets produce an essential metabolite whose abundance can be measured through robust techniques like LC-MS/MS (MAT2A, PGD, PLD2, etc.). Kinases, while heavily studied, may not be ideal in this manner because understanding the downstream outcomes of inhibition may require the conservation of a phosphorylation site or process. Because of this, antibodies detecting phosphorylated substrates might not be reliable for detection of on-target effects of inhibitors without additional studies to validate these tools. To be given final consideration, a potential drug target must have ideal characteristics in a majority of the above criteria, and a viable means to measure on-target pharmacologic action of a small molecule on its target. To establish whether a biological outcome could be measured following inhibition, we performed literature searches on known inhibitors of human orthologs of schistosome target proteins. If there was an established, robust method to measure target engagement and pharmacologic action, then we prioritized these genes for additional studies.

### Identification of inhibitors for drug targets to test on adult schistosome parasites

Because each essential gene in our RNAi screen originated from a specific human ortholog included in our initial bioinformatic queries, we revisited this dataset to determine which orthologous human drug target prompted discovery of our drug target candidates in schistosomes. After this, we manually searched various databases (GeneCards, DrugBank, Google, Therapeutic Target Database, etc.)^31,33,143^ and chemical vendors (Medchemexpress, Selleck Chem, etc.) for inhibitors that target these human proteins. In each instance, we consulted published literature to determine which inhibitors were worth testing. We pursued inhibitors for targets that displayed favorable characteristics in each of our *in silico* prioritization criteria, and whose actions could possibly be assessed for on-target activity. Preference was also given to inhibitors that were more drug-like in nature^63–67^, had been validated in other systems and had sub-micromolar IC_50_ or EC_50_ values determined experimentally. Also, we pursued drugs that were in clinical trials, FDA-approved, or whose targets had published structures in complex with these inhibitors.

### Evaluation of commercially available inhibitors

Selected compounds were evaluated *in vitro* for preliminary potency at a single concentration (10 μM in 0.1% DMSO). Roughly 5 adult worm pairs were placed in 3mL Basch medium in a 12-well culture vessel, and incubated at 37°C, 5% CO2 for the length of the experiment. Compound was administered at D0, D1, and D2, media and drug refreshed every 24 hr. Worms were then allowed to remain in culture until D5, where a final assessment of worm attachment to tissue culture substate was made by light microscopy. Observations on movement and morphology were also recorded. A negative control (0.1% DMSO) and positive control (PZQ; 10 μM in 0.1% DMSO) were included in each experiment, and each experiment was repeated in triplicate for validation. For compounds that caused complete detachment in replicate experiments, the treatment was expanded to three concentrations (10 μM, 5 μM, and 1 μM) following the same treatment schedule and controls as detailed above. If compounds still displayed potency at 1 μM, then a full dose-response titration (50 μM - 10 nM) of compound was performed to assess schistosomicidal potency. Each titration experiment was performed in triplicate according to the treatment regimen described above, using DMSO and PZQ as appropriate controls. Morphologies and motility were assessed by light microscopy, and dose-response curves were calculated using the primary readout of parasite attachment to tissue culture substrate in GraphPad Prism.

### Western blot analysis

For drug treatment studies, 10 male adult worms (single or paired with females) were supplemented with either 0.1% DMSO or inhibitor (0.1% DMSO final). For all experiments assessing on-target effects of potential parasite p97 inhibitors, CB-5083 was used as a positive control (0.1% DMSO). After 48 hr, male parasites were separated from females using 0.25% tricaine in Basch Media 169, flash frozen in liquid nitrogen, and stored at −80°C until further processing. Male worm samples were homogenized with a pestle in 50 µL lysis buffer containing 2X sample buffer [0.471 M Tris (pH 6.7 with phosphoric acid), 20% glycerol, 5% SDS), protease inhibitor cocktail (Roche, cOmplete Mini, EDTA-free Tablets) and 10 mM DTT. The lysates were then sonicated on high for 5 min (30 sec on, 30 sec off) using a Bioruptor UCD-200. Lysates were centrifuged for 5 min at 10,000 g to remove debris. Total protein was measured using the Detergent Compatible Bradford Assay (Pierce). 50 µg of protein samples denatured in SDS Sample buffer (95°C for 5 min) were separated on a Bio-Rad 4-20% TGX Stain-Free gel along with Precision Plus Protein Dual Color Standards (Bio-Rad) as a marker. Proteins were then transferred to a nitrocellulose membrane (Bio-Rad). The membrane was blocked in a 1:5 solution of Li-Cor Odyssey Blocking buffer in PBST for 1 hr before being immunoblotted overnight at 4°C with 1:500 K48-linkage Specific Polyubiquitin Antibody (Cell Signaling Technology, 4289S) and 0.01 µg/mL mouse anti-actin antibody (1:10,000) (Developmental Studies Hybridoma Bank, JLA20) diluted in a 1:5 solution of Li-Cor Odyssey Blocking buffer in PBST. The membrane was washed 3x in TBST and then incubated in 1:5 Li-Cor Odyssey Blocking buffer containing the secondary antibodies (1:10,000 Li-Cor, 925-68071, goat anti-rabbit IRDye 680 RD, and 1:20,000 Li-Cor, 925-32280, goat anti-mouse IgM IRDye 800CW) for 1 hr at RT. The blot was washed in TBST 3x before being imaged on a LiCor Odyssey Infrared Imager.

### Purification of recombinant p97

For recombinant expression, full-length wildtype *Schistosoma mansoni* p97 (Smp_018240) or *Homo sapiens* p97 (P55072) were synthesized and cloned into pET28a(+) with a C-terminal 6x His tag by GenScript (Piscaaway, NJ). *E. coli* BL21 (DE3) containing the desired plasmid were grown in LB medium containing 50 μg/L kanamycin while shaking at 37 °C to an OD600 of 0.6-0.8. The temperature was reduced to 18°C and 0.8 mM isopropyl-beta-D-thiogalactopyranoside (IPTG) was added. The bacterial culture was harvested 16hr later by centrifugation. The resulting pellet was suspended in 60 mL lysis buffer/2L bacterial culture [20 mM Tris (pH 7.4), 300 mM NaCl, 5 mM MgCl2, 20 mM imidazole, 1% Triton X-100, 1 mg/mL lysozyme, 10% glycerol, 3 mM β-mercaptoethanol, 0.2 mM PMSF, and protease inhibitor tablet (Roche)]. The cells were incubated for 30min at 4°C, rocking, then subjected to subsequent lysis by dounce homogenization (10x) and sonication (3x 30s pulses, 65% amplitude, 3min rest on ice in between). The lysate was centrifuged at 40,000 x rpm for 35 min at 4°C. The resulting supernatant was incubated with Ni-NTA beads equilibrated to lysis buffer for 1 hr at 4°C before being loaded onto a gravity column. The bead bed was washed with wash buffer [20 mM Tris (pH 7.4), 500 mM NaCl, 5 mM MgCl2, 40 mM imidazole, 0.05% Triton X-100, 10% glycerol, 3 mM β-mercaptoethanol, 0.2 mM PMSF, 0.001 mg/mL Aprotinin, and 0.002 mg/mL Leupeptin], then eluted in the same buffer with 250 mM imidazole in wash buffer. Fractions containing p97 were concentrated using Pierce Protein Concentrators (100k MWCO) and loaded onto a gel-filtration column (Superdex 200PG) and eluted using SEC buffer [20 mM Tris (pH 7.4), 180 mM NaCl, 5 mM MgCl2, and 10% glycerol] at 0.5 mL/min flow rate on a BioRad NGC Chromatography System. Fractions corresponding to an apparent molecular weight of 500 – 600 kDa were collected and analyzed by 4– 20% SDS/PAGE to evaluate purity. Concentration was determined using a BSA standard. Fractions containing purified protein were aliquoted, snap frozen in liquid nitrogen, and stored at −80 °C until use.

### Cryo-EM Sample Preparation

Prior to grid preparation, 20 µl of a freshly thawed aliquot of 1 mg/mL purified p97 [20 mM Tris (pH 7.4), 180 mM NaCl, 5 mM MgCl2, and 1 mM tris(2-carboxyethyl)phosphine (TCEP)] was incubated with 100 µM inhibitor (0.5% DMSO final) for 30 min at room temperature. For experiments with ATPγS, the protein was pre-incubated with 1 mM ATPγS (dissolved in H_2_O) for 30 minutes at room temperature before making grids. 3 µL were applied to holey carbon grids (Quantifoil R1.2/1.3, 300 mesh copper) and plunge frozen using the Vitrobot Mark IV System (Thermo Fisher). Grids were glow-discharged using a PELCO easiGlow glow discharge apparatus (Ted Pella) for 80 s at 30 mA before use sample application. Grids were screened at UTSW Cryo Electron Microscopy Facility (CEMF) on a 200kV microscope and the best grid with optimal particle distribution was used for data collection.

### Cryo-EM Data Acquisition and Processing

Apo *S. mansoni* p97 dataset was collected at the CEMF on a 300kV Titan Krios microscope equipped with BioQuantum energy filter and K3 detector (Gatan) at CEMF in non-CDS mode at 105 kX magnification with SerialEM^144^ at ~15 e^−^/pix/sec, with defocus range −0.9 to −2.2 μm and total dose of 50 e^−^/Å^2^ in 24 hr, which yielded 6,939 movies at pixel size of 0.83 Å. ATPγS- and CB-5083-bound data sets were collected on the same microscope in CDS mode at 105 kX magnification with SerialEM at ~9 e^−^/pix/sec, with defocus range −0.9 to −2.2 μm, and −1.0 to −2.4 μm, respectively, and total dose of ~60 e^−^/Å^2^ in 24 hr. 5,730 and 4,579 movies were yielded, respectively. 739-bound sample was collected on a Titan Krios microscope equipped with a Selectris energy filter and Falcon 4i detector (Thermo Fisher Scientific) at CEMF at 165 kX magnification with SerialEM at ~9 e^−^/pix/sec, with defocus range −0.9 to −2.2 μm and total dose of 60 e^−^/Å^2^ in 24 hr, which yielded 9,352 movies at pixel size of 0.738 Å. 804-bound sample was collected at the Pacific Northwest Cryo-EM Center (PNCC) for 48 hr data collection on a Titan Krios microscope equipped with a Selectris energy filter and Falcon 4i detector at 165 kX magnification with EPU at ~9 e^−^/pix/sec, with defocus range −0.9 to −2.2 μm and total dose of 50 e^−^/Å^2^, yielded a total of 17,748 movies at pixel size of 0.7296 Å.

All data processing was performed using the software cryoSPARC v4.2 (Apo, ATPγS- and CB5083-bound) or v4.5 (739- and 804-bound) using default parameters^145^. For K3 datasets, a ½ F-crop factor was applied during patch motion correction, followed by patch CTF estimation. For Falcon 4i datasets, eer upsampling factor of 1 was used during motion correction and CTF estimation in cryoSPARC Live.

The first dataset collected was the CB-5083-bound dataset, a subset of 135 micrographs was selected and blob picker was used to pick 18,949 particles, which went through two rounds of 2D classification to yield 8,843 particles that were used as input for Topaz training. Topaz was used to repick 16,463 particles from the same subset, which were used to generate three *ab initio* reconstructions with C6 symmetry. Topaz was then used to pick particles from the whole dataset of 4,174 micrographs, which yielded an initial 968,168 particles. After one round of 2D classification, 529,148 particles and the three *ab initio* models were used as input for heterogenous refinement with C6 symmetry. 75% of the particles were partitioned in a class that resembles p97. These particles were further submitted in *ab initio* reconstruction to obtain three new models and cleaned up with two rounds of heterogenous refinement with C6 symmetry. In the last heterogenous refinement, 88% of the particles, which includes 174,830 particles, resembled p97 and were submitted for final refinement using homogenous refinement with C6 symmetry applied, with global CTF refinement and local motion correction to obtain a 2.85 Å map. The map was sharpened by deepEMhancer^146^, which was used for modeling.

For the Apo dataset, the CB-5083-bound map was used to generate templates which was used for template picking on the whole dataset to yield 1,949,076 initial particles. These were cleaned up for two rounds of 2D classification to yield 878,454 particles. After two rounds of heterogenous refinement with C6 symmetry (using three of the same CB-5083-bound map as input model), 85% of the particles, which includes 478,635 particles, resembled p97 in the last iteration and were submitted for final refinement using homogenous refinement with C6 symmetry and global CTF refinement to obtain a 2.72 Å map. The map was also sharpened by deepEMhancer to obtain final map used for modeling.

Similar procedure was applied to the ATPγS-bound dataset. Template picking yielded 1,681,589 initial picks, which were cleaned up by two rounds of 2D classification. 1,074,975 particles were submitted to one round of heterogenous refinement with C6 symmetry. 62% of the particles, which includes 665,490 particles, were submitted to final refinement using homogenous refinement with C6 symmetry, global and local CTF refinements, to obtain a 2.2 Å map. DeepEMhancer sharpened map was used for modeling.

For compound 739-bound dataset, the same CB-5083-bound map was used to generate templates for Template picking in cryoSPARC Live, which yielded 2,812,039 particles. After two rounds of 2D classification, 373,213 particles were selected and used in two rounds of heterogenous refinement using two of the same CB-5083-bound map as initial references and C6 symmetry. 78% of the particles in the last heterogenous refinement job, which includes 101,826 particles, were submitted to final refinement using non-uniform refinement with C6 symmetry applied, with global and CTF refinements, to yield a 3.06 Å map. Unsharpened map was used for modeling.

Similar procedure was applied for compound 804-bound dataset. Template picking in cryoSPARC Live yielded 2,272,397 particles, which were cleaned up in one round of 2D classification to 966,087 particles. After two rounds of heterogeneous refinement, 95% of the particles in the last iteration, including 823,143 particles, were submitted to final refinement with homogenous refinement with C6 symmetry applied, and global and local CTF refinements, to yield a 2.76 Å map. DeepEMhancer sharpened map was used for modeling.

### Cryo-EM model building, refinement and analysis

ModelAngelo^147^ was used to generate an initial 3D-model for CB-5083-bound p97 in one chain. Errors were fixed manually in Coot^148^. The other chains were built based on the initial ModelAngelo model by manually fitting into the map in Chimera, followed by merging all chains in Coot. Modeling of the CB-5083 is kept consistent with previous published models (PDB: 7RLI, 6MCK) as the ligand density is ambiguous for the primary amide extension on the inhibitor. Model refinement was carried out by multiple cycles of real-space refinement in Phenix^149^ and manual model building using Coot. The same initial model from ModelAngelo was used to build the Apo model, where the single chain was docked into the map in ChimeraX and undergone manual modeling in Coot. The rest of the chains were built using molrep in CCP-EM^150^. Model refinement was carried out as described above. The ATPγS-bound model was built using the fully built Apo model with 6 chains as an initial model, which underwent model refinement using ISOLDE^151^ in ChimeraX. ATPγS and Mg^2+^ were first modelled manually in coot and included in refinement in Phenix in later steps. For compound 739- and 804-bound models, the fully built Apo model was used and first refined against their respective maps in Phenix. Restraints for the compounds and its conjugation to a cysteine side chain were generated in JLigand^152^ and used in Coot for manual building of the compounds into chain D. Other copies of the ligands were built in Coot using NCS ligand tool and refined individually to optimize map fitting and geometry. All model validation was performed and analyzed in PyMOL 2.2.3 or 3.1.3 where the final figures were generated^140^. Ligplot^141^ was used to generate interaction figures.

### Evaluation of p97’s ATPase activity and compound potency

For manual ATP consumption assays, 1 µM p97 was incubated with 1% DMSO or inhibitor (1% DMSO final) in assay buffer [50 mM Tris (pH 7.5), 20 mM MgCl2, 1 mM EDTA, 0.5 mM tris(2-carboxyethyl)phosphine (TCEP)] for 5 minutes prior to the addition of 20 µM ATP in a 384-well plate for a total reaction volume of 10 µL, and a final concentration of 500nM enzyme and 10 µM ATP. The reaction was allowed to proceed for 50 minutes at room temperature before adding equivalent volume of Kinase-Glo Plus reagent (Promega), and a further incubation of 10 minutes in the dark. Luminescence was read using a BioTek Synergy 2 plate reader (model no. 3375752).

To determine IC_50_ values, compounds were assayed at a range of concentrations (100 μM, 50 μM, 10 μM, 5 μM, 1 μM, 500 nM, 100 nM, 50 nM, 10 nM, 5 nM, 1 nM, and 0 nM) in triplicate, maintaining a concentration of 1% DMSO. The percent of inhibition was calculated using CB-5083 as a control for 100% inhibition. IC_50_ values were calculated using GraphPad Prism.

### High-throughput screening (HTS) assay

Kinase-Glo (Promega, part K1214) was purchased from Promega. Microtiter plates (384 wells) were purchased from Perkin Elmer. All chemicals included in the UTSW library were sourced from commercial sources [ChemDiv (150,000 compounds), ChemBridge (125,500), ComGenex (22,000), Prestwick Chemical, (1,100) and TimTec (500)] or UT Southwestern chemists (https://www.utsouthwestern.edu/research/core-facilities/high-throughput-screening/libraries/). For initial screening, compounds were dispensed by Echo 555 robot for a final concentration of 10 µM (0.2% DMSO). Luminescence was read by EnVision multimode plate reader (Perkin-Elmer). Assay conditions were consistent between manual and high-throughput assays [10 µM ATP, 50 mM Tris (pH 7.5), 20 mM MgCl2, 1 mM EDTA, 0.5 mM tris(2-carboxyethyl)phosphine (TCEP)]. The plates were then incubated for 50 min at room temperature before addition of Kinase-Glo reagent and further incubation in the dark for 10 minutes before reading luminescence. CB-5083 was used as a control for 100% inhibition, 10 µM ATP for maximal signal, and DMSO for 0% inhibition. Raw values and values normalized to CB-5083 control were obtained to determine activity of compounds on the enzyme. Hit compounds were validated using a dose-response curve (19.8 µM, 9.92 µM, 4.96 µM, 3.72 µM, 2.48 µM, 1,24 µM, 0.619 µM, 0.342 µM, 0.256 µM, 0.171 µM, 0.0854 µM, 0.0427 µM, and 0 µM) in triplicate. Compounds were also subjected to a counter screen using a dose-response curve against the full-length human p97 enzyme to determine preliminary selectivity.

### Mass spectrometry analysis of benzoxazole propiolamide covalent inhibitors

1mg/mL aliquots of recombinant *S. mansoni* p97 [20 mM Tris (pH 7.4), 180 mM NaCl, 5 mM MgCl2, and 1 mM tris(2-carboxyethyl)phosphine (TCEP)] were thawed to room temperature. Purified enzyme was then incubated with 100 µM inhibitor (242 and 243) (1% DMSO) or negative control (1% DMSO only) for 20 min prior to submission for intact mass spectrometry analysis. In-gel LC-MS/MS experiments were performed similarly on recombinant *S. mansoni* and *H. sapiens* p97. However, the reaction with 100 µM inhibitor (242 and 243) (1% DMSO) or negative control (1% DMSO only) was halted by the addition of 4X loading (1X final concentration). The samples were denatured in SDS sample buffer (95°C for 5min), then separated on a Bio-Rad 4-20% TGX Stain-Free gel along with Precision Plus Protein Dual Color Standards (Bio-Rad) as a marker. Following separation, bands were detected using GelCode™ Blue Safe Protein Stain (Cat# 24594). Peptides were excised from the gel corresponding to the molecular weight of p97, then submitted to UTSW Proteomics Core for in-gel trypsin digestion prior to extraction, desalting, and downstream LC-MS/MS analysis.

### HepG2 cytotoxicity assay of benzoxazole propiolamide covalent inhibitors

HepG2 cells (ATCC) were plated at 2500 cells/well in 50 µL/well, dispersing cells using a 25g needle to break up clumps prior to plating. Plates were incubated for 24hr at 37C, 5% CO2. Following 24hr, 50 µL media containing 0.6% DMSO was added to a set of wells on one plate. The plate and Cell Titer Glo 2.0 reagent were brought to RT. A T=0 read was taken by adding 40 µL reagent/well, rocking the plate at 100 rpm for 2 min, then reading luminescence ~10 min later. Serial 1:3 dilutions of compound at 333 x concentration were made in DMSO. These stocks were then diluted 167-fold into media to make a 2x concentrated stock containing 0.6% DMSO. 50 µL 2x concentrated compound was added to wells containing cells in 50 µL 24 hr after plating. Concentrations were tested in triplicate at following values: 30, 10, 3.3, 1.1, 0.37, 0.12, and 0.04 µM. Plates were incubated for 72 hr at 37C, 5% CO2. Following equilibration to room temperature, 40 µL undiluted reagent was added to each well, the plate rocked at 100 rpm for 2 min and then read for luminescent signal ~10 min later.

GI50 is the concentration that causes 50% growth inhibition. It corrects for the cell count at time zero; thus, GI50 is the concentration of test drug where 100 × (T - T0)/(C - T0) = 50. IC_50_ is the concentration that results in 50% growth inhibition relative to control and is calculated as the test drug concentration where 100 x (T/C) = 50. Values are plotted in excel and the 50% point determined by interpolation. Values were revisualized in Prism.

## Chemical Synthesis Methods

### General procedure 1 (Amidation by HATU)

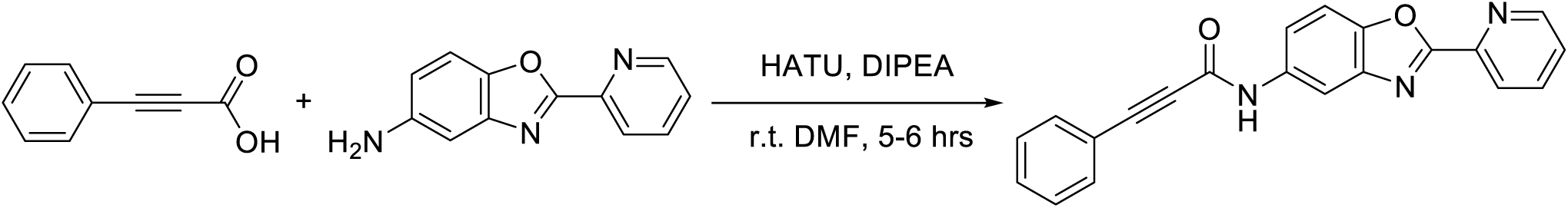

#### Experimental procedure (General Procedure 1, Amidation by HATU)

To a solution of phenylpropiolic acid (0.3 mmol, 1 equiv) in *N,N*-dimethylformamide (2.0 mL) was added amines (0.3 mmol, 1 equiv), HATU (125.5 mg, 0.33 mmol, 1.1 equiv) and *N,N*-diisopropylethylamine (104.5 μL, 0.6 mmol, 2 equiv). The mixture was sealed in a 20-mL scintillation vial and stirred at room temperature for 5 to 6 hours. The reaction crudes were poured onto ice to form off-white precipitates. The precipitates were collected by filtration and washed with water to afford off-white solid as products, yields are 23.7-99.6%.

### General procedure 2 (Amidation by T_3_P)

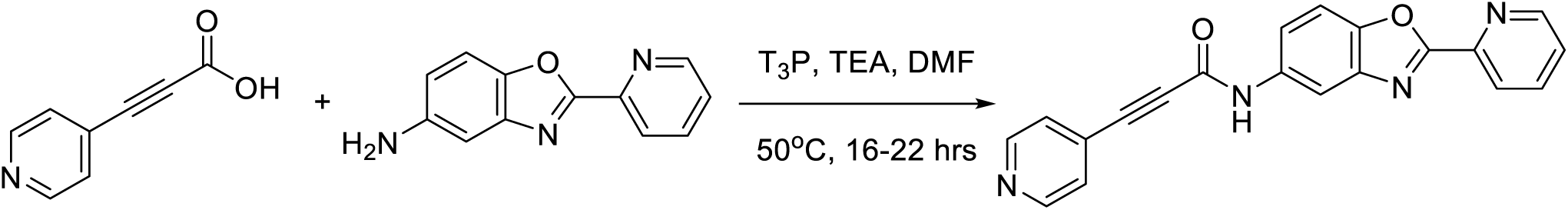

#### Experimental procedure (General Procedure 2, Amidation by T_3_P)

To a solution of 3-(4-pyridyl)propiolic acid (0.1 mmol, 1 equiv) in *N,N*-dimethylformamide (0.8 mL) was added amines (0.1 mmol, 1 equiv), T_3_P (propylphosphonic anhydride solution, ≥50 wt. % in ethyl acetate, 95.5 mg, 0.15 mmol, 1.5 equiv) and triethylamine (44.6 μL, 0.32 mmol, 3.2 equiv). The mixture was sealed in a 20-mL scintillation vial and stirred at room temperature to 50°C for about 20 hours. The reaction crudes were poured onto ice to form grey to brown precipitates. The precipitates were collected by filtration and washed with water and/or dichloromethane and hexane to afford off-white solid as products, yields are 32.1-68.5%.

### General procedure 3 (Synthesis of Pyridinyl Benzoxazole Amines by PPA)

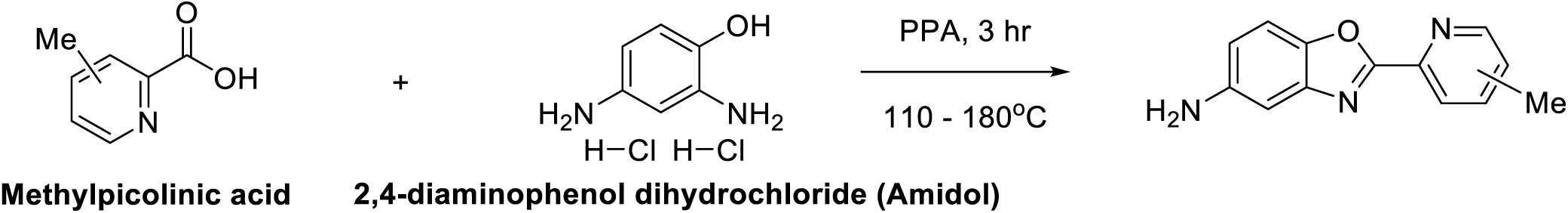

#### Experimental procedure (General Procedure 3, Synthesis of Pyridinyl Benzoxazole Amines by PPA)

To PPA (polyphosphoric acid, 1.0 g, 0.5 mL) at 110°C were added simultaneously 2,4-diaminophenol dihydrochloride (98.5 mg, 0.5 mmol, 1.0 equiv) and methylpicolinic acid (68.6 mg, 0.5 mmol, 1.0 equiv). The resulting mixture was then heated to 180°C for 3 hours. The mixture turned very gluey, hard to stir. It may or may not form bubbles at this moment. After cooling, ice chips were added to the crude and then sodium carbonate powder (about 1.2 g) was added in small portions. Lots of bubbles formed in the beginning. When no more bubbles formed, it turned to greenish-black suspension, and pH is about 11. The greenish-black suspension was extracted by dichloromethane. The combined organic layers were concentrated on vacuum to yield yellow crude solid. It was further purified by column chromatography or trituration. Yield 33.6-78.6%.

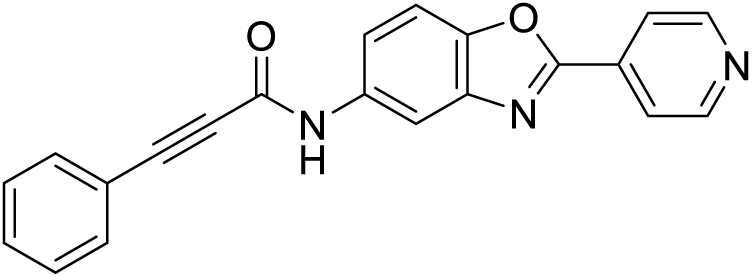

3-phenyl-*N*-(2-(pyridin-4-yl)benzo[*d*]oxazol-5-yl)propiolamide (**242**)

#### Experimental procedure

**242** was synthesized by using phenylpropiolic acid (44.8 mg, 0.3 mmol, 1 equiv) and 2-pyridin-4-yl-benzooxazol-5-ylamine (61.2 mg, 0.3 mmol, 1 equiv) in General Procedure 1. The reaction was stirred at room temperature for 5 hours. The product was washed by water and 1 M NaOH as 70.0 mg brown solid. Yield 71.2%. ^1^H NMR (400 MHz, Chloroform-*d*) δ 8.62 (d, J = 5.5 Hz, 2H), 8.01 (d, J = 2.1 Hz, 1H), 7.94 (d, J = 5.2 Hz, 2H), 7.59 (dd, J = 8.9, 2.1 Hz, 1H), 7.43 (d, J = 8.0 Hz, 3H), 7.28 (d, J = 7.2 Hz, 1H), 7.26 – 7.22 (m, 2H). ^13^C NMR (101 MHz, Chloroform-*d*) δ 161.0, 151.8, 150.1, 147.5, 141.5, 135.6, 134.4, 132.4, 130.1, 128.4, 121.1, 119.9, 119.5, 111.7, 110.8, 85.9, 83.2. ESI-MS *m/z* = 340.1 ([M+H]^+^), C_21_H_13_N_3_O_2_ requires 340.1

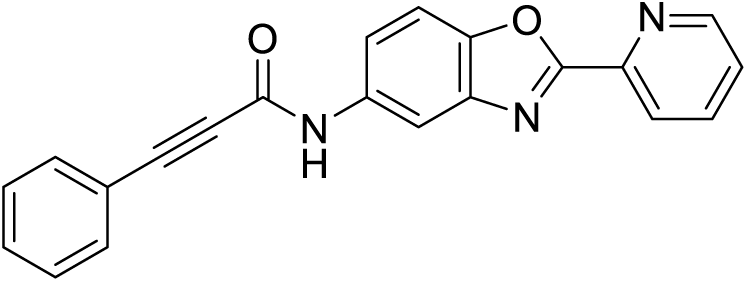

3-phenyl-*N*-(2-(pyridin-2-yl)benzo[*d*]oxazol-5-yl)propiolamide (**243**)

#### Experimental procedure

**243** was synthesized by using phenylpropiolic acid (44.4 mg, 0.3 mmol, 1 equiv) and 2-pyridin-2-yl-benzooxazol-5-ylamine (61.5 mg, 0.3 mmol, 1 equiv) in General Procedure 1. The reaction was stirred at room temperature for 5 hours. The product was washed by water as 98.4 mg off-white solid. Yield 99.6%. ^1^H NMR (400 MHz, Chloroform-*d*) δ 8.59 (d, J = 4.9 Hz, 1H), 8.11 (d, J = 7.9 Hz, 1H), 8.01 (s, 1H), 7.72 (t, J = 7.5 Hz, 1H), 7.54 (d, J = 8.8 Hz, 1H), 7.36 (d, J = 8.8 Hz, 1H), 7.33 – 7.25 (m, 3H), 7.19 (d, J = 7.4 Hz, 1H), 7.12 (t, J = 7.5 Hz, 2H). ^13^C NMR (101 MHz, Chloroform-*d*) δ 162.0, 151.8, 149.9, 147.6, 145.2, 141.5, 137.3, 135.3, 132.3, 130.0, 128.3, 125.8, 123.5, 119.8, 119.2, 111.8, 110.9, 85.9, 83.3. ESI-MS *m/z* = 340.1 ([M+H]^+^), C_21_H_13_N_3_O_2_ requires 340.1

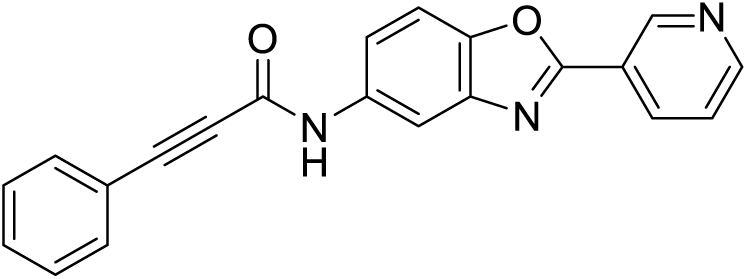

3-phenyl-*N*-(2-(pyridin-3-yl)benzo[*d*]oxazol-5-yl)propiolamide (**244**)

#### Experimental procedure

**244** was synthesized by using phenylpropiolic acid (45.4 mg, 0.3 mmol, 1 equiv) and 2-(3-pyridinyl)-5-benzoxazolamine (63.1 mg, 0.3 mmol, 1 equiv) in General Procedure 1. The reaction was stirred at room temperature for 5 hours. The product was washed by water as 95.4 mg red-brown solid. Yield 94.1%. ^1^H NMR (400 MHz, Chloroform-*d*) δ 9.36 (s, 1H), 8.69 (d, J = 4.9 Hz, 1H), 8.45 (d, J = 8.0 Hz, 1H), 7.98 (d, J = 2.1 Hz, 1H), 7.67 (dd, J = 8.9, 2.1 Hz, 1H), 7.56 – 7.51 (m, 2H), 7.45 (dd, J = 8.0, 4.9 Hz, 2H), 7.38 (dd, J = 8.0, 1.7 Hz, 1H), 7.35 – 7.29 (m, 2H). ^13^C NMR (101 MHz, Chloroform-*d*) δ 161.1, 151.8, 151.5, 148.0, 147.4, 141.5, 135.4, 135.1, 132.4, 130.1, 128.4, 1241, 123.4, 119.9, 118.9, 111.4, 110.6, 85.8, 83.2. ESI-MS *m/z* = 340.1 ([M+H]^+^), C_21_H_13_N_3_O_2_ requires 340.1

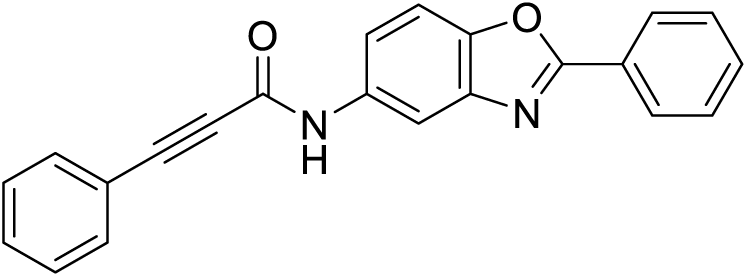

3-phenyl-*N*-(2-phenylbenzo[*d*]oxazol-5-yl)propiolamide (**245**)

#### Experimental procedure

**245** was synthesized by using phenylpropiolic acid (44.9 mg, 0.3 mmol, 1 equiv) and 2-phenyl-benzooxazol-5-ylamine (63.6 mg, 0.3 mmol, 1 equiv) in General Procedure 1. The reaction was stirred at room temperature for 5 hours. The product was washed by water as 103.7 mg off-white solid. Quantitative yield. ^1^H NMR (400 MHz, Chloroform-*d*) δ 8.18 (dd, J = 7.8, 2.0 Hz, 2H), 7.91 (d, J = 2.1 Hz, 1H), 7.67 (dd, J = 8.8, 2.1 Hz, 1H), 7.54 – 7.47 (m, 6H), 7.39 – 7.30 (m, 3H). ^13^C NMR (101 MHz, Chloroform-*d*) δ 164.0, 151.8, 147.4, 141.7, 135.0, 132.4, 131.8, 130.1, 128.9, 128.4, 127.4, 126.4, 120.0, 118.3, 111.1, 110.5, 85.8, 83.3. ESI-MS *m/z* = 339.1 ([M+H]^+^), C_22_H_14_N_2_O_2_ requires 339.1

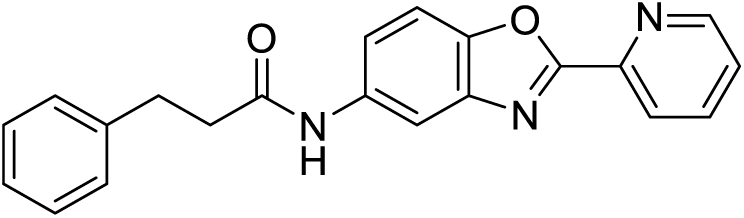

3-phenyl-*N*-(2-(pyridin-2-yl)benzo[*d*]oxazol-5-yl)propanamide (**326**)

#### Experimental procedure

**326** was synthesized by hydrogenation of **243**. A hydrogenation balloon was attached to a sealed vial with alkyne (17.2 mg, 0.05 mmol, 1 equiv) and 10% palladium on carbon (10.6 mg, 0.01 mmol, 0.2 equiv) in a mixed solvent of 1 mL ethyl acetate and 1 mL of methanol. The mixture was stirred at room temperature for 6 hours. The catalyst was filtered off by celite and solvents were removed. The product was washed by hexane to obtain as 16.8 mg off-white solid. Yield 96.5%. ^1^H NMR (400 MHz, Chloroform-*d*) δ 8.78 (d, J = 4.7 Hz, 1H), 8.31 (d, J = 7.9 Hz, 1H), 7.92 – 7.83 (m, 2H), 7.52 (d, J = 8.7 Hz, 1H), 7.46 – 7.41 (m, 2H), 7.31 – 7.18 (m, 5H), 3.06 (t, J = 7.5 Hz, 2H), 2.68 (t, J = 7.7 Hz, 2H). ^13^C NMR (101 MHz, Chloroform-*d*) δ 171.7, 161.9, 150.0, 145.4, 140.7, 137.4, 135.8, 128.4, 128.2, 126.1, 125.5, 123.5, 119.4, 118.9, 111.6, 110.8, 110.5, 38.8, 31.6. ESI-MS *m/z* = 344.2 ([M+H^+^]), C_21_H_17_N_3_O_2_ requires 344.1

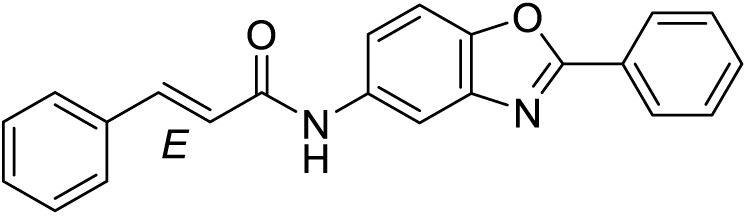

*N*-(2-phenylbenzo[*d*]oxazol-5-yl)cinnamamide (**135**)

#### Experimental procedure

**135** was synthesized by using *trans*-cinnamic acid (8.3 mg, 0.05 mmol, 1 equiv) and 2-phenyl-benzooxazol-5-ylamine (9.9 mg, 0.05 mmol, 1 equiv) in General Procedure 1. The reaction was stirred at room temperature overnight. The product was washed by dichloromethane and hexane as 13.6 mg off-white solid. Yield 84.8%. ^1^H NMR (400 MHz, Chloroform-*d*) δ 8.14 (dd, J = 7.7, 2.1 Hz, 2H), 7.87 (d, J = 2.2 Hz, 1H), 7.78 (dd, J = 8.7, 2.2 Hz, 1H), 7.67 (d, J = 15.6 Hz, 1H), 7.51 – 7.41 (m, 6H), 7.33 – 7.26 (m, 3H), 6.62 (d, J = 15.6 Hz, 1H). ^13^C NMR (151 MHz, Chloroform-*d*) δ 164.8, 163.9, 147.3, 141.8, 141.7, 135.8, 134.8, 131.8, 129.7, 128.9, 128.7, 127.8, 127.5, 126.6, 120.9, 118.4, 110.7, 110.5. ESI-MS *m/z* = 341.1 ([M+H^+^]), C_22_H_16_N_2_O_2_ requires 341.1

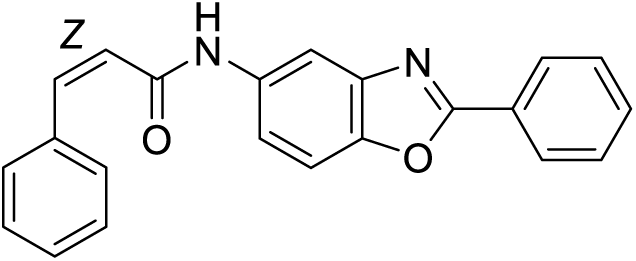

(*Z*)-3-phenyl-*N*-(2-phenylbenzo[*d*]oxazol-5-yl)acrylamide (**324**)

#### Experimental procedure

**324** was synthesized by hydrogenation of **245**. A hydrogenation balloon was attached to a sealed vial with alkyne (15.7 mg, 0.05 mmol, 1 equiv) and Lindlar catalyst (2.1 mg, 0.001 mmol, 0.02 equiv) in 0.25 mL of methanol. The mixture was stirred at room temperature for 20 hours until the complete disappearance of starting material. The catalyst was filtered off by celite and solvents were removed. The product was washed by hexane to obtain 14.6 mg off-white solid. Yield 92.4%. ^1^H NMR (400 MHz, Chloroform-*d*) δ 8.13 (dd, J = 7.8, 2.0 Hz, 2H), 7.69 (d, J = 2.1 Hz, 1H), 7.53 – 7.41 (m, 8H), 7.33 – 7.22 (m, 2H), 6.83 (d, J = 12.5 Hz, 1H), 6.09 (d, J = 12.5 Hz, 1H). ^13^C NMR (101 MHz, Chloroform-*d*) δ 165.6, 163.9, 149.9, 147.4, 141.8, 137.9, 134.8, 131.8, 129.1, 128.9, 128.8, 1284, 127.5, 126.6, 124.1, 118.5, 111.1, 110.4. ESI-MS *m/z* = 341.1 ([M+H^+^]), C_22_H_16_N_2_O_2_ requires 341.1

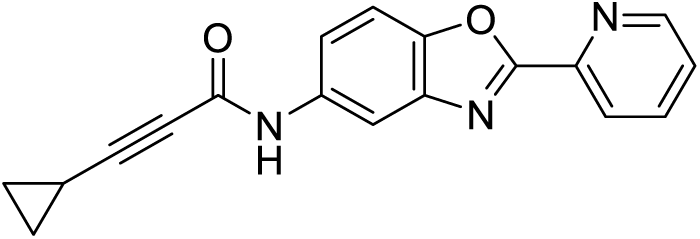

3-cyclopropyl-*N*-(2-(pyridin-2-yl)benzo[*d*]oxazol-5-yl)propiolamide (**732**)

#### Experimental procedure

**732** was synthesized by using 3-cyclopropylprop-2-ynoic acid (7.5 mg, 0.05 mmol, 1 equiv) and 2-pyridin-2-yl-benzooxazol-5-ylamine (10.6 mg, 0.05 mmol, 1 equiv) in General Procedure 1. The reaction was stirred at room temperature overnight. The product was washed by water, followed by trituration in dichloromethane and hexane to afford 15.0 mg light-brown solid. Yield 90.8%. ^1^H NMR (600 MHz, Chloroform-*d*) δ 8.74 (d, J = 4.1 Hz, 1H), 8.28 (d, J = 7.9 Hz, 1H), 7.93 (d, J = 2.2 Hz, 1H), 7.87 (td, J = 7.8, 1.8 Hz, 1H), 7.62 (dd, J = 8.8, 2.2 Hz, 1H), 7.53 (d, J = 8.9 Hz, 1H), 7.44 (ddd, J = 7.6, 4.8, 1.2 Hz, 1H), 1.35 (tt, J = 8.2, 5.1 Hz, 1H), 0.93 – 0.83 (m, 4H). ^13^C NMR (151 MHz, Chloroform-*d*) δ 162.3, 151.9, 150.2, 147.8, 145.6, 141.7, 137.5, 135.4, 126.0, 123.6, 119.3, 111.7, 111.2, 92.4, 71.1, 9.0, −0.6. ESI-MS *m/z* = 304.1 ([M+H^+^]), C_18_H_13_N_3_O_2_ requires 304.1

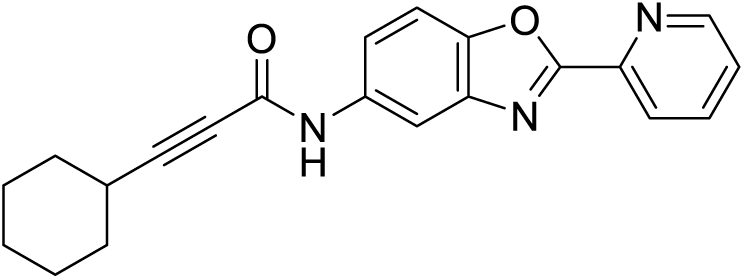

3-cyclohexyl-*N*-(2-(pyridin-2-yl)benzo[*d*]oxazol-5-yl)propiolamide (**733**)

#### Experimental procedure

**733** was synthesized by using 3-cyclohexylpropiolic acid (9.1 mg, 0.05 mmol, 1 equiv) and 2-pyridin-2-yl-benzooxazol-5-ylamine (11.1 mg, 0.05 mmol, 1 equiv) in General Procedure 1. The reaction was stirred at room temperature overnight. The product was washed by water, followed by trituration in dichloromethane and hexane to afford 12.6 mg off-white solid. Yield 69.4%. ^1^H NMR (600 MHz, Chloroform-*d*) δ 8.72 (d, J = 4.7 Hz, 1H), 8.26 (d, J = 7.8 Hz, 1H), 7.94 (t, J = 2.8 Hz, 1H), 7.87 (ddd, J = 7.7, 5.2, 2.5 Hz, 1H), 7.62 (dt, J = 8.9, 2.3 Hz, 1H), 7.52 (dd, J = 9.1, 3.8 Hz, 1H), 7.43 (s, 1H), 2.47 (td, J = 9.5, 4.7 Hz, 1H), 1.81 (d, J = 12.9 Hz, 2H), 1.68 (p, J = 5.3, 4.5 Hz, 2H), 1.51 – 1.40 (m, 3H), 1.27 (s, br, 3H). ^13^C NMR (151 MHz, Chloroform-*d*) δ 162.2, 152.1, 150.2, 147.8, 145.5, 141.7, 137.5, 135.4, 126.0, 123.6, 119.4, 111.8, 111.2, 92.5, 75.8, 31.7, 29.0, 25.6, 24.8. ESI-MS *m/z* = 346.2 ([M+H^+^]), C_21_H_19_N_3_O_2_ requires 346.2

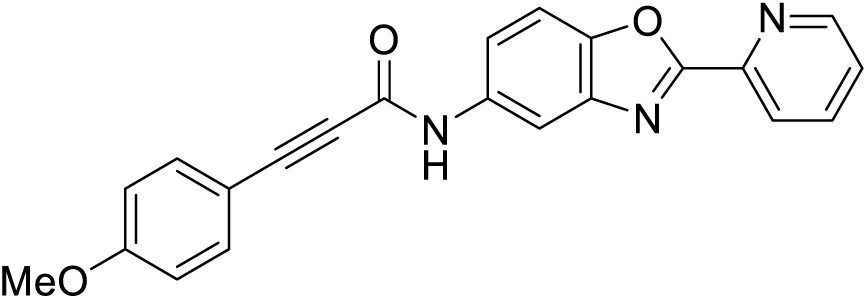

3-(4-methoxyphenyl)-*N*-(2-(pyridin-2-yl)benzo[*d*]oxazol-5-yl)propiolamide (**734**)

#### Experimental procedure

**734** was synthesized by using 3-(4-methoxyphenyl)propiolic acid (9.7 mg, 0.05 mmol, 1 equiv) and 2-pyridin-2-yl-benzooxazol-5-ylamine (10.2 mg, 0.05 mmol, 1 equiv) in General Procedure 1. The reaction was stirred at room temperature overnight. The product was eluted by 2.5% methanol in dichloromethane from silica gel column to afford 11.1 mg light-brown solid. Yield 62.2%. ^1^H NMR (600 MHz, Chloroform-*d*) δ 8.81 (dt, J = 4.7, 1.3 Hz, 1H), 8.35 (d, J = 7.9 Hz, 1H), 8.10 (d, J = 2.0 Hz, 1H), 8.01 (s, 1H), 7.90 (td, J = 7.8, 1.8 Hz, 1H), 7.62 (dd, J = 8.8, 2.0 Hz, 1H), 7.59 (d, J = 8.7 Hz, 1H), 7.49 (d, J = 8.8 Hz, 2H), 7.46 (ddd, J = 7.6, 4.8, 1.2 Hz, 1H), 6.87 (d, J = 8.8 Hz, 2H), 3.82 (s, 3H). ^13^C NMR (151 MHz, Chloroform-*d*) δ 162.4, 161.2, 151.5, 150.3, 148.1, 145.7, 142.2, 137.2, 134.8, 134.4, 125.7, 123.6, 119.1, 114.3, 112.1, 111.6, 111.3, 86.8, 82.6, 55.4. ESI-MS *m/z* = 370.2 ([M+H^+^]), C_22_H_15_N_3_O_3_ requires 370.1

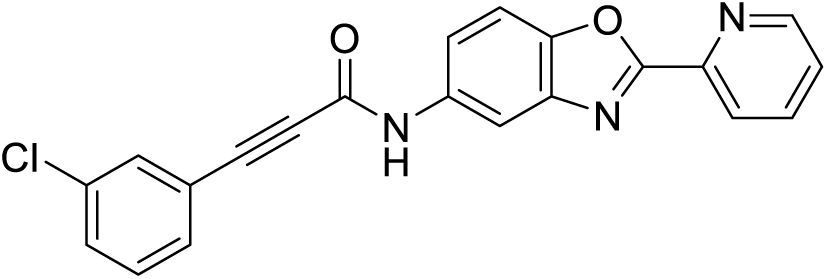

3-(3-chlorophenyl)-*N*-(2-(pyridin-2-yl)benzo[*d*]oxazol-5-yl)propiolamide (**735**)

#### Experimental procedure

**735** was synthesized by using 3-(3-chlorophenyl)propiolic acid (10.4 mg, 0.05 mmol, 1 equiv) and 2-pyridin-2-yl-benzooxazol-5-ylamine (10.0 mg, 0.05 mmol, 1 equiv) in General Procedure 1. The reaction was stirred at room temperature overnight. The product was eluted by 5% methanol in dichloromethane from silica gel column, followed by trituration in dichloromethane and hexane to afford 4.2 mg off-white solid. Yield 23.7%. ^1^H NMR (600 MHz, Chloroform-*d*) δ 8.82 (d, J = 4.8 Hz, 1H), 8.36 (d, J = 7.9 Hz, 1H), 8.09 (d, J = 1.9 Hz, 1H), 7.91 (td, J = 7.8, 1.7 Hz, 1H), 7.63 (d, J = 8.7 Hz, 1H), 7.61 (dd, J = 8.8, 1.9 Hz, 1H), 7.55 (t, J = 1.8 Hz, 1H), 7.48 – 7.45 (m, 3H), 7.33 (t, J = 7.9 Hz, 1H). ^13^C NMR (151 MHz, Chloroform-*d*) δ 162.5, 161.5, 152.3, 150.3, 148.3, 145.8, 142.3, 137.2, 134.4, 132.3, 130.7, 129.8, 125.8, 123.6, 119.1, 112.2, 112.1, 111.4, 84.3, 84.0. ESI-MS *m/z* = 374.1 ([M+H^+^]), C_21_H_12_ClN_3_O_2_ requires 374.1

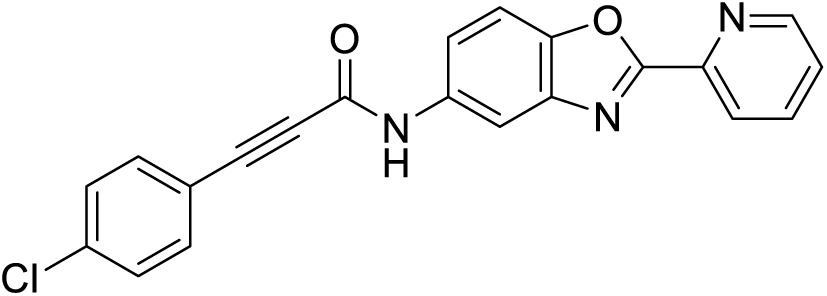

3-(4-chlorophenyl)-*N*-(2-(pyridin-2-yl)benzo[*d*]oxazol-5-yl)propiolamide (**736**)

#### Experimental procedure

**736** was synthesized by using 3-(4-chlorophenyl)propiolic acid (11.2 mg, 0.05 mmol, 1 equiv) and 2-pyridin-2-yl-benzooxazol-5-ylamine (10.6 mg, 0.05 mmol, 1 equiv) in General Procedure 1. The reaction was stirred at room temperature overnight. The product was eluted by 5% methanol in dichloromethane from silica gel column, followed by trituration in dichloromethane and hexane to afford 10.8 mg off-white solid. Yield 57.6%. ^1^H NMR (600 MHz, Chloroform-*d*) δ 8.80 (d, J = 5.1 Hz, 1H), 8.34 (d, J = 7.9 Hz, 1H), 8.08 (d, J = 2.1 Hz, 1H), 7.90 (t, J = 7.7 Hz, 1H), 7.62 (dd, J = 8.9, 1.9 Hz, 1H), 7.59 (d, J = 8.7 Hz, 1H), 7.45 – 7.42 (m, 3H), 7.33 – 7.30 (m, 2H). ^13^C NMR (151 MHz, Chloroform-*d*) δ 162.4, 161.7, 152.1, 150.3, 148.2, 145.7, 142.1, 137.2, 134.5, 133.7, 129.0, 125.8, 123.6, 119.1, 112.2, 111.8, 111.3, 84.8, 84.0. ESI-MS *m/z* = 374.1 ([M+H^+^]), C_21_H_12_ClN_3_O_2_ requires 374.1

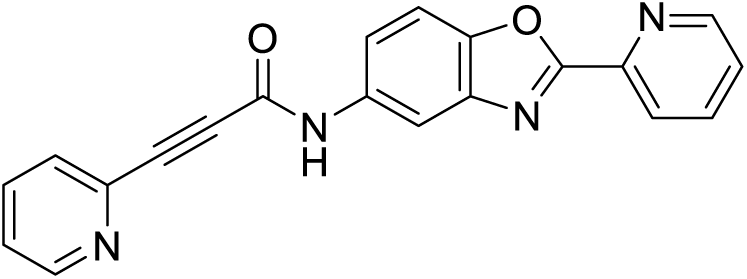

3-(pyridin-2-yl)-*N*-(2-(pyridin-2-yl)benzo[*d*]oxazol-5-yl)propiolamide (**737**)

#### Experimental procedure

**737** was synthesized by using 3-(pyridin-2-yl)propiolic acid (3.8 mg, 0.025 mmol, 1 equiv) and 2-pyridin-2-yl-benzooxazol-5-ylamine (4.5 mg, 0.025 mmol, 1 equiv) in General Procedure 2. The reaction was stirred at room temperature for 20 hours. The product was eluted by 5% methanol in dichloromethane from silica gel column to afford 3.5 mg light yellow solid. Yield 48.3%. ^1^H NMR (600 MHz, Chloroform-*d*) δ 8.82 (dt, J = 4.7, 1.3 Hz, 1H), 8.66 (dt, J = 4.9, 1.3 Hz, 1H), 8.37 (d, J = 7.9 Hz, 1H), 8.24 (d, J = 5.1 Hz, 1H), 8.13 (d, J = 2.1 Hz, 1H), 7.91 (td, J = 7.7, 1.8 Hz, 1H), 7.77 (td, J = 7.7, 1.8 Hz, 1H), 7.65 (dt, J = 7.8, 1.1 Hz, 1H), 7.63 (d, J = 8.7 Hz, 1H), 7.59 (dd, J = 8.8, 2.1 Hz, 1H), 7.47 (ddd, J = 7.6, 4.7, 1.2 Hz, 1H), 7.39 (ddd, J = 7.7, 4.9, 1.2 Hz, 1H). ^13^C NMR (151 MHz, Chloroform-*d*) δ 162.5, 150.4, 150.3, 150.3, 148.3, 145.8, 142.2, 140.6, 137.2, 136.7, 134.5, 128.7, 125.7, 124.7, 123.6, 118.9, 112.1, 111.4, 83.6, 82.1. ESI-MS *m/z* = 340.1 ([M+H^+^]), C_20_H_12_N_4_O_2_ requires 340.1

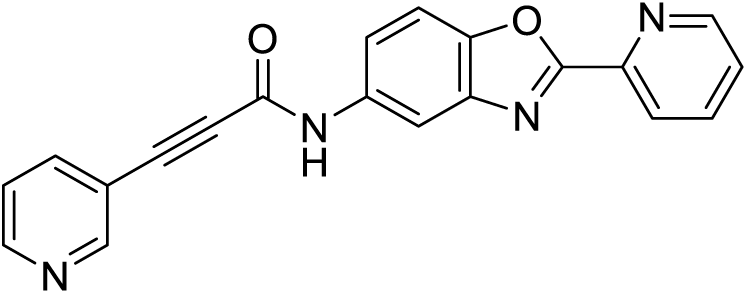

*N*-(2-(pyridin-2-yl)benzo[*d*]oxazol-5-yl)-3-(pyridin-3-yl)propiolamide (**738**)

#### Experimental procedure

**738** was synthesized by using 3-(pyridin-3-yl)propiolic acid (15.4 mg, 0.1 mmol, 1 equiv) and 2-pyridin-2-yl-benzooxazol-5-ylamine (24.0 mg, 0.1 mmol, 1 equiv) in General Procedure 2. The reaction was stirred at room temperature for 26 hours. The product was washed by water, dichloromethane and hexane to afford 24.4 mg off-white solid. Yield 68.5%. ^1^H NMR (600 MHz, Methanol-*d_4_*) δ 8.81 (d, J = 2.0 Hz, 1H), 8.77 (dt, J = 4.8, 1.3 Hz, 1H), 8.64 (dd, J = 5.0, 1.6 Hz, 1H), 8.39 (d, J = 7.9 Hz, 1H), 8.27 (d, J = 2.0 Hz, 1H), 8.08 (dt, J = 7.9, 1.9 Hz, 1H), 8.06 (dd, J = 7.8, 1.7 Hz, 1H), 7.71 (d, J = 8.8 Hz, 1H), 7.66 (dd, J = 8.8, 2.1 Hz, 1H), 7.62 (ddd, J = 7.6, 4.8, 1.2 Hz, 1H), 7.52 (dd, J = 7.9, 5.0 Hz, 1H). ^13^C NMR (151 MHz, Methanol-*d_4_*) δ 163.5, 153.3, 152.5, 151.1, 150.9, 149.1, 146.3, 142.9, 141.4, 139.1, 136.8, 127.4, 125.0, 124.8, 120.3, 119.2, 112.9, 112.2, 87.2, 82.6. ESI-MS *m/z* = 340.1 ([M+H^+^]), C_20_H_12_N_4_O_2_ requires 340.1

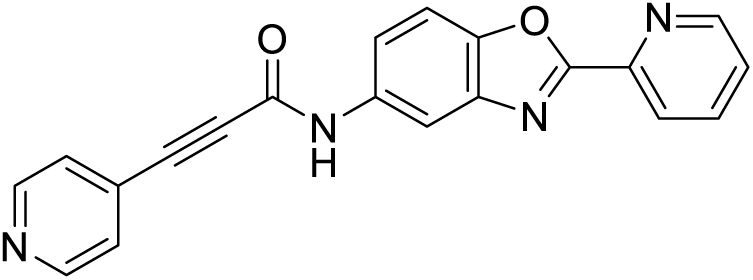

*N*-(2-(pyridin-2-yl)benzo[*d*]oxazol-5-yl)-3-(pyridin-4-yl)propiolamide (**739**)

#### Experimental procedure

**739** was synthesized by using 3-(4-pyridyl)propiolic acid (17.7 mg, 0.1 mmol, 1 equiv) and 2-pyridin-2-yl-benzooxazol-5-ylamine (23.5 mg, 0.1 mmol, 1 equiv) in General Procedure 2. The reaction was stirred at room temperature for 26 hours. The product was eluted by 5% methanol in dichloromethane from silica gel column to afford 25.5 mg light yellow solid. Yield 67.3%. ^1^H NMR (400 MHz, Chloroform-*d*) δ 8.81 (d, J = 4.9 Hz, 1H), 8.65 (d, J = 5.2 Hz, 2H), 8.35 (d, J = 7.9 Hz, 1H), 8.07 (d, J = 2.1 Hz, 1H), 7.91 (td, J = 7.8, 1.7 Hz, 1H), 7.71 – 7.64 (m, 1H), 7.62 (d, J = 8.8 Hz, 1H), 7.47 (ddd, J = 7.5, 4.8, 1.1 Hz, 1H), 7.44 (d, J = 6.0 Hz, 2H). ^13^C NMR (151 MHz, Chloroform-*d*) δ 162.2, 150.5, 150.1, 149.5, 147.9, 145.3, 141.6, 137.4, 135.1, 129.0, 126.1, 125.9, 123.5, 119.2, 111.7, 111.2, 87.0, 81.5. ESI-MS *m/z* = 340.1 ([M+H^+^]), C_20_H_12_N_4_O_2_ requires 340.1

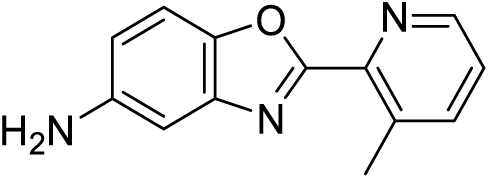

2-(3-methylpyridin-2-yl)benzo[*d*]oxazol-5-amine (**804-i**)

#### Experimental procedure

**804-i** was synthesized by using 3-methylpicolinic acid (68.8 mg, 0.5 mmol, 1 equiv) in General Procedure 3. The product was eluted by 5% methanol in dichloromethane from silica gel column to afford 51.4 mg brown solid. Yield 45.5%. ^1^H NMR (400 MHz, Chloroform-*d*) δ 8.66 (dd, J = 5.1, 1.5 Hz, 1H), 7.69 (ddd, J = 7.8, 1.7, 0.8 Hz, 1H), 7.45 (d, J = 8.6 Hz, 1H), 7.33 (dd, J = 7.8, 4.6 Hz, 1H), 7.11 (d, J = 2.3 Hz, 1H), 6.77 (dd, J = 8.6, 2.3 Hz, 1H), 2.85 (s, 3H). ^13^C NMR (151 MHz, Chloroform-*d*) δ 161.7, 147.3, 144.3, 144.3, 143.7, 142.8, 139.8, 135.0, 124.6, 114.8, 111.1, 105.3, 21.0. ESI-MS *m/z* = 226.1 ([M+H^+^]), C_13_H_11_N_3_O requires 226.1

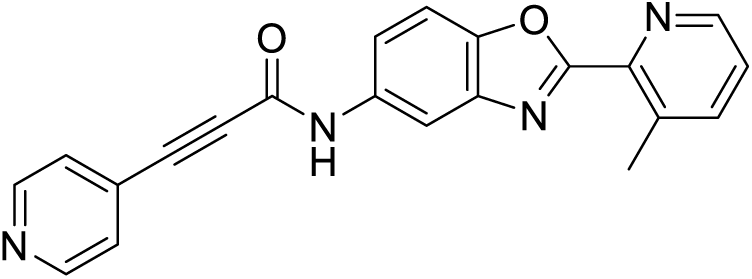

*N*-(2-(3-methylpyridin-2-yl)benzo[*d*]oxazol-5-yl)-3-(pyridin-4-yl)propiolamide (**804**)

#### Experimental procedure

**804** was synthesized by using 3-(4-pyridyl)propiolic acid (8.1 mg, 0.05 mmol, 1 equiv) and 2-(3-methylpyridin-2-yl)benzo[*d*]oxazol-5-amine (**804-i**, 12.2 mg, 0.05 mmol, 1 equiv) in General Procedure 2. The reaction was stirred at 50°C for 26 hours. The product was eluted by 5% methanol in dichloromethane from silica gel column to afford 11.0 mg light yellow solid. Yield 57.3%. ^1^H NMR (600 MHz, Methanol-*d_4_*) δ 8.63 – 8.61 (m, 2H), 8.58 (dd, J = 4.7, 1.6 Hz, 1H), 8.25 (d, J = 2.0 Hz, 1H), 7.84 (dt, J = 7.8, 1.2 Hz, 1H), 7.66 (d, J = 8.8 Hz, 1H), 7.62 (dd, J = 8.8, 2.1 Hz, 1H), 7.60 – 7.57 (m, 2H), 7.47 (dd, J = 7.8, 4.7 Hz, 1H), 2.83 (s, 3H). ^13^C NMR (151 MHz, Methanol-*d_4_*) δ 163.1, 151.9, 150.3, 148.3, 147.9, 144.5, 142.8, 141.5, 136.8, 136.2, 130.5, 127.4, 126.5, 120.1, 113.1, 111.8, 87.9, 82.4, 21.1. ESI-MS *m/z* = 355.2 ([M+H^+^]), C_21_H_14_N_4_O_2_ requires 355.1

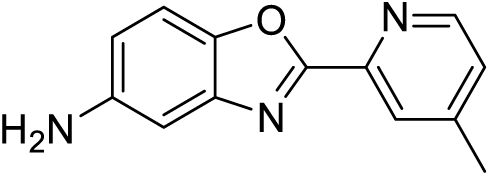

2-(4-methylpyridin-2-yl)benzo[*d*]oxazol-5-amine (**805-i**)

#### Experimental procedure

**805-i** was synthesized by using 4-methylpicolinic acid (140.6 mg, 1.0 mmol, 1 equiv) in General Procedure 3. The product was eluted by 5% methanol in dichloromethane from silica gel column to afford 181.6 mg yellow solid. Yield 78.6%. ^1^H NMR (400 MHz, Chloroform-*d*) δ 8.61 (d, J = 5.0 Hz, 1H), 8.12 (s, 1H), 7.39 (d, J = 8.7 Hz, 1H), 7.20 (d, J = 5.0 Hz, 1H), 7.06 (d, J = 2.3 Hz, 1H), 6.74 (dd, J = 8.7, 2.3 Hz, 1H), 2.42 (s, 3H). ^13^C NMR (151 MHz, Chloroform-*d*) δ 161.5, 149.4, 148.0, 145.4, 144.4, 144.0, 142.3, 125.8, 123.6, 114.4, 110.7, 104.5, 20.6. ESI-MS *m/z* = 226.1 ([M+H^+^]), C_13_H_11_N_3_O requires 226.1

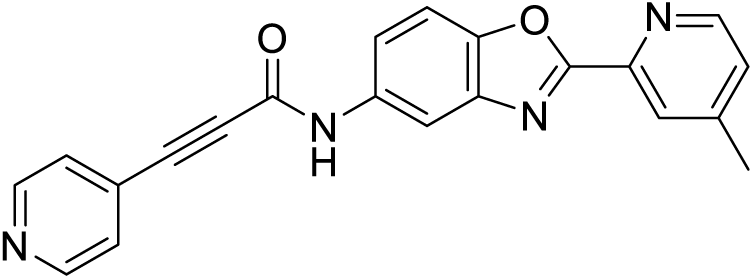

*N*-(2-(4-methylpyridin-2-yl)benzo[*d*]oxazol-5-yl)-3-(pyridin-4-yl)propiolamide (**805**)

#### Experimental procedure

**805** was synthesized by using 3-(4-pyridyl)propiolic acid (7.5 mg, 0.05 mmol, 1 equiv) and 2-(4-methylpyridin-2-yl)benzo[*d*]oxazol-5-amine (**805-i**, 10.9 mg, 0.05 mmol, 1 equiv) in General Procedure 2. The reaction was stirred at 50°C for 22 hours. The product was eluted by 5% methanol in dichloromethane from silica gel column to afford 5.5 mg off-white solid. Yield 32.1%. ^1^H NMR (600 MHz, Methanol-*d_4_*) δ 8.65 – 8.63 (m, 2H), 8.59 (d, J = 5.0 Hz, 1H), 8.25 (d, J = 2.1 Hz, 1H), 8.22 (d, J = 1.6 Hz, 1H), 7.69 (d, J = 8.8 Hz, 1H), 7.64 (dd, J = 8.8, 2.1 Hz, 1H), 7.62 – 7.59 (m, 2H), 7.43 (dd, J = 5.2, 1.9 Hz, 1H), 2.52 (s, 3H). ^13^C NMR (151 MHz, Methanol-*d_4_*) δ 163.5, 152.0, 150.9, 150.6, 150.4, 149.1, 146.0, 142.8, 136.6, 130.5, 128.2, 127.5, 125.5, 120.2, 112.8, 112.1, 87.9, 82.5, 21.2. ESI-MS *m/z* = 355.2 ([M+H^+^]), C_21_H_14_N_4_O_2_ requires 355.1

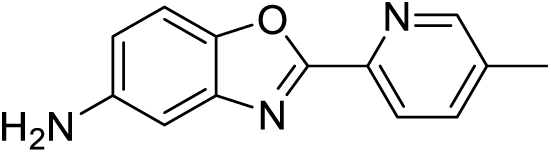

2-(5-methylpyridin-2-yl)benzo[*d*]oxazol-5-amine (**806-i**)

#### Experimental procedure

**806-i** was synthesized by using 5-methylpicolinic acid (137.5 mg, 1.0 mmol, 1 equiv) in General Procedure 3. The product was eluted by 5% methanol in dichloromethane from silica gel column to afford 124.4 mg yellow solid. Yield 55.1%. ^1^H NMR (400 MHz, Chloroform-*d*) δ 8.51 (s, 1H), 8.10 (d, J = 8.1 Hz, 1H), 7.63 (d, J = 7.7 Hz, 1H), 7.36 (dd, J = 8.7, 1.5 Hz, 1H), 7.04 (s, 1H), 6.76 (dt, J = 8.6, 2.2 Hz, 1H), 2.36 (s, 3H). ^13^C NMR (151 MHz, Chloroform-*d*) δ 161.8, 150.4, 144.7, 144.0, 143.0, 142.3, 137.5, 135.8, 122.7, 114.7, 111.0, 104.9, 18.4. ESI-MS *m/z* = 226.1 ([M+H^+^]), C_13_H_11_N_3_O requires 226.1

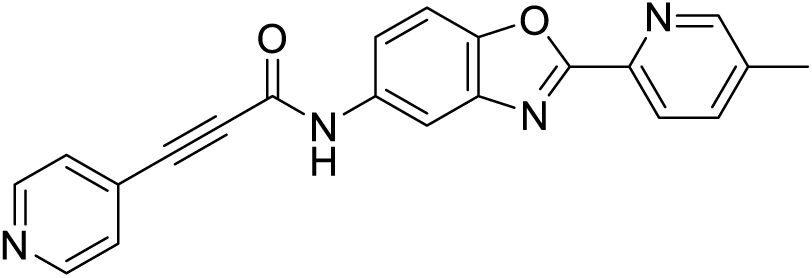

*N*-(2-(5-methylpyridin-2-yl)benzo[*d*]oxazol-5-yl)-3-(pyridin-4-yl)propiolamide (**806**)

#### Experimental procedure

**806** was synthesized by using 3-(4-pyridyl)propiolic acid (8.0 mg, 0.05 mmol, 1 equiv) and 2-(5-methylpyridin-2-yl)benzo[*d*]oxazol-5-amine (**806-i**, 10.4 mg, 0.05 mmol, 1 equiv) in General Procedure 2. The reaction was stirred at 50°C for 22 hours. The product was eluted by 5% methanol in dichloromethane from silica gel column to afford 6.8 mg light yellow solid. Yield 41.6%. ^1^H NMR (600 MHz, Methanol-*d_4_*) δ 8.64 – 8.61 (m, 2H), 8.58 (d, J = 2.3 Hz, 1H), 8.24 (d, J = 8.0 Hz, 1H), 8.21 (d, J = 2.0 Hz, 1H), 7.83 (dd, J = 8.0, 3.0 Hz, 1H), 7.66 (d, J = 8.8 Hz, 1H), 7.62 (dd, J = 8.8, 2.1 Hz, 1H), 7.60 – 7.58 (m, 2H), 2.46 (s, 3H). ^13^C NMR (151 MHz, Methanol-*d_4_*) δ 163.5, 151.9, 151.3, 150.4, 148.9, 143.5, 142.7, 139.2, 138.1, 136.5, 130.5, 127.4, 124.3, 120.0, 112.7, 112.0, 87.9, 82.4, 18.7. ESI-MS *m/z* = 355.2 ([M+H^+^]), C_21_H_14_N_4_O_2_ requires 355.1

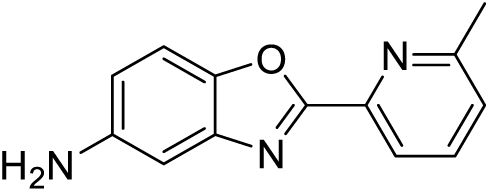

2-(6-methylpyridin-2-yl)benzo[*d*]oxazol-5-amine (**807-i**)

#### Experimental procedure

**807-i** was synthesized by using 6-methylpicolinic acid (138.8 mg, 1.0 mmol, 1 equiv) in General Procedure 3. The product was eluted by 5% methanol in dichloromethane from silica gel column to afford 76.5 mg orange solid. Yield 33.6%. ^1^H NMR (400 MHz, Chloroform-*d*) δ 8.11 (d, J = 7.8 Hz, 1H), 7.74 (t, J = 7.8 Hz, 1H), 7.42 (d, J = 8.6 Hz, 1H), 7.28 (d, J = 7.7 Hz, 1H), 7.08 (d, J = 2.3 Hz, 1H), 6.75 (dd, J = 8.6, 2.3 Hz, 1H), 2.71 (s, 3H). ^13^C NMR (151 MHz, Chloroform-*d*) δ 162.0, 159.3, 145.6, 145.0, 144.1, 142.9, 137.1, 125.1, 120.5, 114.7, 111.3, 105.2, 24.7. ESI-MS *m/z* = 226.1 ([M+H^+^]), C_13_H_11_N_3_O requires 226.1

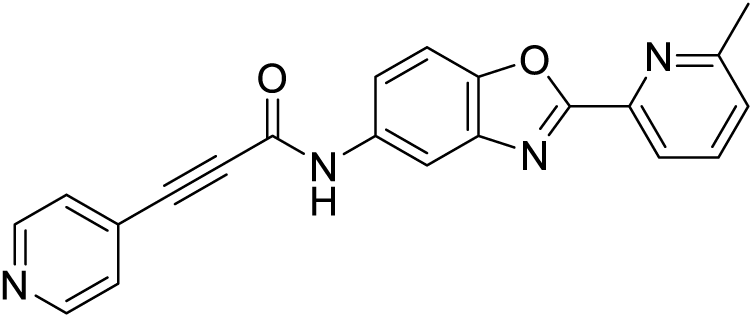

*N*-(2-(6-methylpyridin-2-yl)benzo[*d*]oxazol-5-yl)-3-(pyridin-4-yl)propiolamide (**807**)

#### Experimental procedure

**807** was synthesized by using 3-(4-pyridyl)propiolic acid (7.1 mg, 0.05 mmol, 1 equiv) and 2-(6-methylpyridin-2-yl)benzo[*d*]oxazol-5-amine (**807-i**, 12.2 mg, 0.05 mmol, 1 equiv) in General Procedure 2. The reaction was stirred at 50°C for 26 hours. The product was eluted by 5% methanol in dichloromethane from silica gel column to afford 7.3 mg light yellow solid. Yield 42.7%. ^1^H NMR (600 MHz, Methanol-*d_4_*) δ 8.66 – 8.61 (m, 2H), 8.23 (d, J = 1.9 Hz, 1H), 8.16 (d, J = 7.8 Hz, 1H), 7.90 (t, J = 7.8 Hz, 1H), 7.69 (d, J = 8.8 Hz, 1H), 7.64 (dd, J = 8.9, 2.0 Hz, 1H), 7.62 – 7.58 (m, 2H), 7.45 (d, J = 7.8 Hz, 1H), 2.68 (s, 3H). ^13^C NMR (151 MHz, Methanol-*d_4_*) δ 163.5, 160.6, 151.8, 150.2, 148.9, 145.5, 142.6, 138.7, 136.3, 130.4, 127.3, 127.0, 121.8, 120.0, 112.7, 111.9, 87.8, 82.3, 24.3. ESI-MS *m/z* = 355.2 ([M+H^+^]), C_21_H_14_N_4_O_2_ requires 355.1

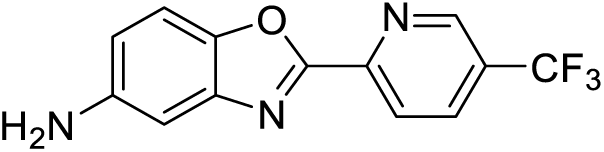

2-(5-(trifluoromethyl)pyridin-2-yl)benzo[*d*]oxazol-5-amine (**808-i**)

#### Experimental procedure

**808-i** was synthesized by using 5-(trifluoromethyl)-2-pyridinecarboxylic acid (98.9 mg, 0.5 mmol, 1 equiv) in General Procedure 3. The product was purified by trituration in dichloromethane and hexane to afford 68.2 mg yellow solid. Yield 47.2%. ^1^H NMR (400 MHz, Chloroform-*d*) δ 9.04 (d, J = 1.2 Hz, 1H), 8.44 (d, J = 8.3 Hz, 1H), 8.12 (dd, J = 8.3, 2.3 Hz, 1H), 7.46 (d, J = 8.7 Hz, 1H), 7.10 (d, J = 2.3 Hz, 1H), 6.82 (dd, J = 8.6, 2.3 Hz, 1H). ^13^C NMR (151 MHz, Chloroform-*d*) δ 160.5, 149.1, 147.1 (q, J = 4.1 Hz), 145.2, 144.6, 142.8, 134.4 (dd, J = 6.8, 3.3 Hz), 127.7 (d, J = 33.3 Hz), 125.7 (q, J = 493.8 Hz), 122.8, 115.8, 111.5, 105.2. ESI-MS *m/z* = 280.1 ([M+H^+^]), C_13_H_8_F_3_N_3_O requires 280.1

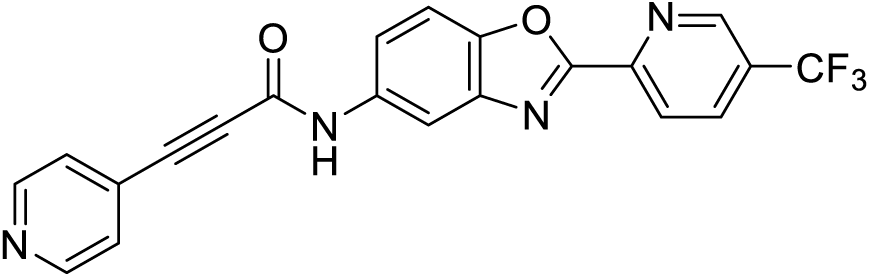

3-(pyridin-4-yl)-*N*-(2-(5-(trifluoromethyl)pyridin-2-yl)benzo[*d*]oxazol-5-yl)propiolamide (**808**)

#### Experimental procedure

**808** was synthesized by using 3-(4-pyridyl)propiolic acid (9.6 mg, 0.05 mmol, 1 equiv) and 2-(5-(trifluoromethyl)pyridin-2-yl)benzo[*d*]oxazol-5-amine (**808-i**, 15.1 mg, 0.05 mmol, 1 equiv) in General Procedure 2. The reaction was stirred at 50°C for 22 hours. The product was eluted by 5% methanol in dichloromethane from silica gel column to afford 9.9 mg light yellow solid. Yield 44.8%. ^1^H NMR (600 MHz, Methanol-*d_4_*) δ 9.03 (d, J = 2.3 Hz, 1H), 8.65 – 8.61 (m, 2H), 8.51 (d, J = 8.2 Hz, 1H), 8.34 – 8.23 (m, 2H), 7.71 – 7.65 (m, 2H), 7.59 (q, J = 2.4 Hz, 2H). ^13^C NMR (151 MHz, Methanol-*d_4_*) δ 160.3, 150.9, 149.3, 148.4, 148.1, 146.8 (d, J = 3.6 Hz), 141.6, 135.7, 135.1 (d, J = 3.6 Hz), 129.4, 128.0 (dd, J = 65.5, 32.5 Hz), 126.4, 123.4, 123.2 (q, J = 272.5 Hz), 119.8, 116.7, 111.2, 86.8, 81.4. ESI-MS *m/z* = 409.2 ([M+H^+^]), C_21_H_11_F_3_N_4_O_2_ requires 409.1

## Supporting information

Supplemental Materials Chemical Synthesis

## Acknowledgements

A portion of this research was supported by NIH grant R24GM154185 and performed at the Pacific Northwest Center for Cryo-EM (PNCC) with assistance from Marcelo De Farias. For support with cryo-EM studies, we thank the Structural Biology Lab (SBL) and the Cryo-Electron Microscopy Facility (CEMF) at UT Southwestern Medical Center which is partially supported by grant RP220582 from the Cancer Prevention & Research Institute of Texas (CPRIT). For support with proteomic studies, we thank the Proteomics Core at UT Southwestern Medical Center. For support with small molecule screening efforts, we thank the High-throughput Screening Core at UT Southwestern Medical Center. For support with toxicity studies, we thank the Preclinical Pharmacology Core at UT Southwestern Medical Center. The Schistosome-infected mice and *B. glabrata* snails were provided by the National Institute of Allergy and Infectious Diseases (NIAID) Schistosomiasis Resource Center of the Biomedical Research Institute (Rockville, MD, USA) through National Institutes of Health (NIH)-NIAID Contract HHSN272201700014I for distribution through BEI Resources. Biorender was used for some schematic material (**Fig. 1*D* and 2*D***). This work was supported by the National Institutes of Health R01AI167967 (J.J.C.) and R01AI150776(J.J.C.), as well as the Welch Foundation I-1948-20240404 (J.J.C.). JJC is an investigator of the Howard Hughes Medical Institute.

---

**Supplemental Figure 1.**
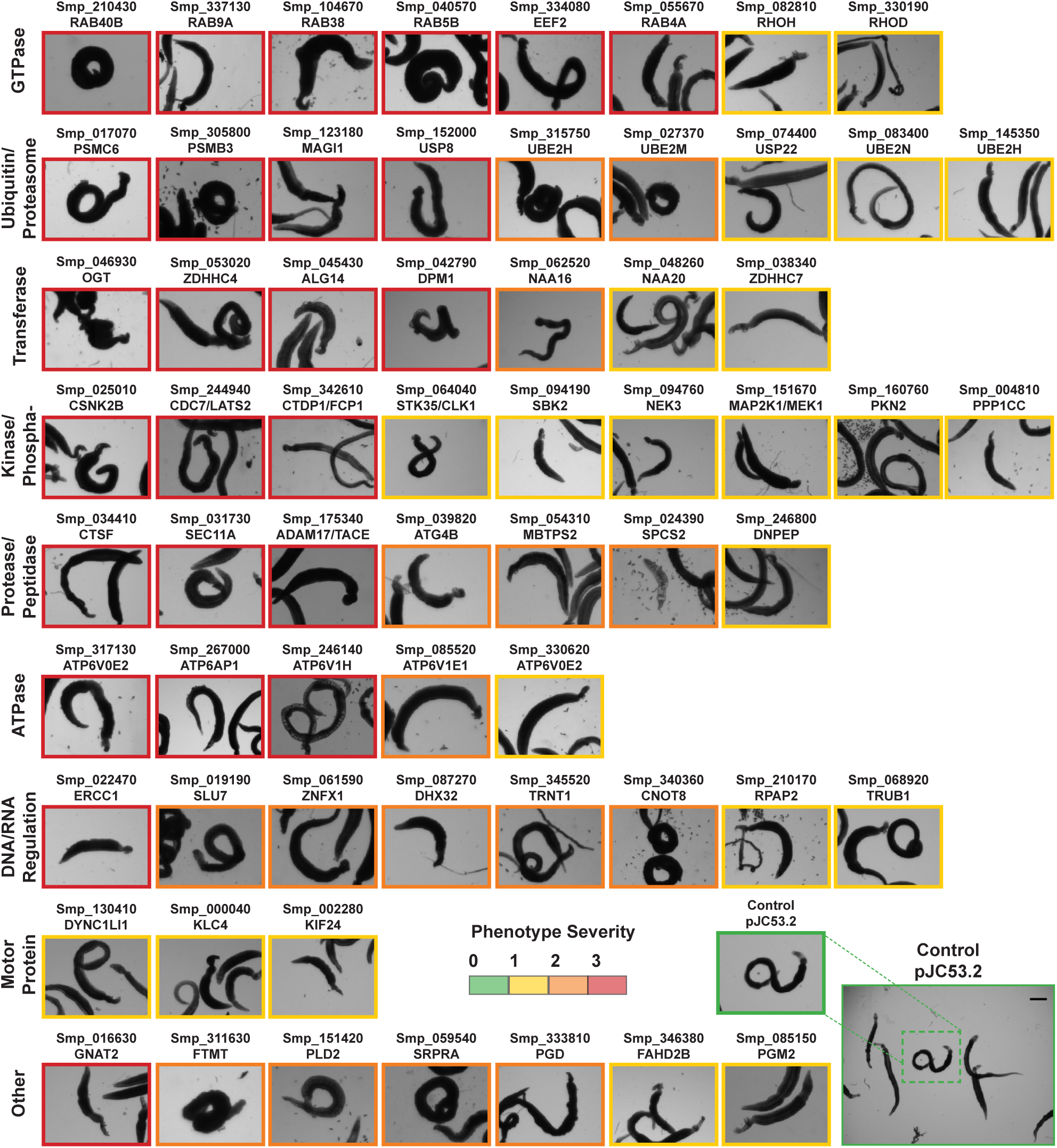
Treatment of adult parasites with dsRNA targeting potential druggable genes. Light microscopy images of adult parasites treated with control dsRNA (pJC53.2) and dsRNAs targeting potential druggable genes within *S. mansoni* that bear homology to human drug targets. Targets were arranged according to order found in Figure 1c of enzymatic activity classification and phenotype severity. Scale bar, 1,000 μm.

**Supplemental Figure 2.**
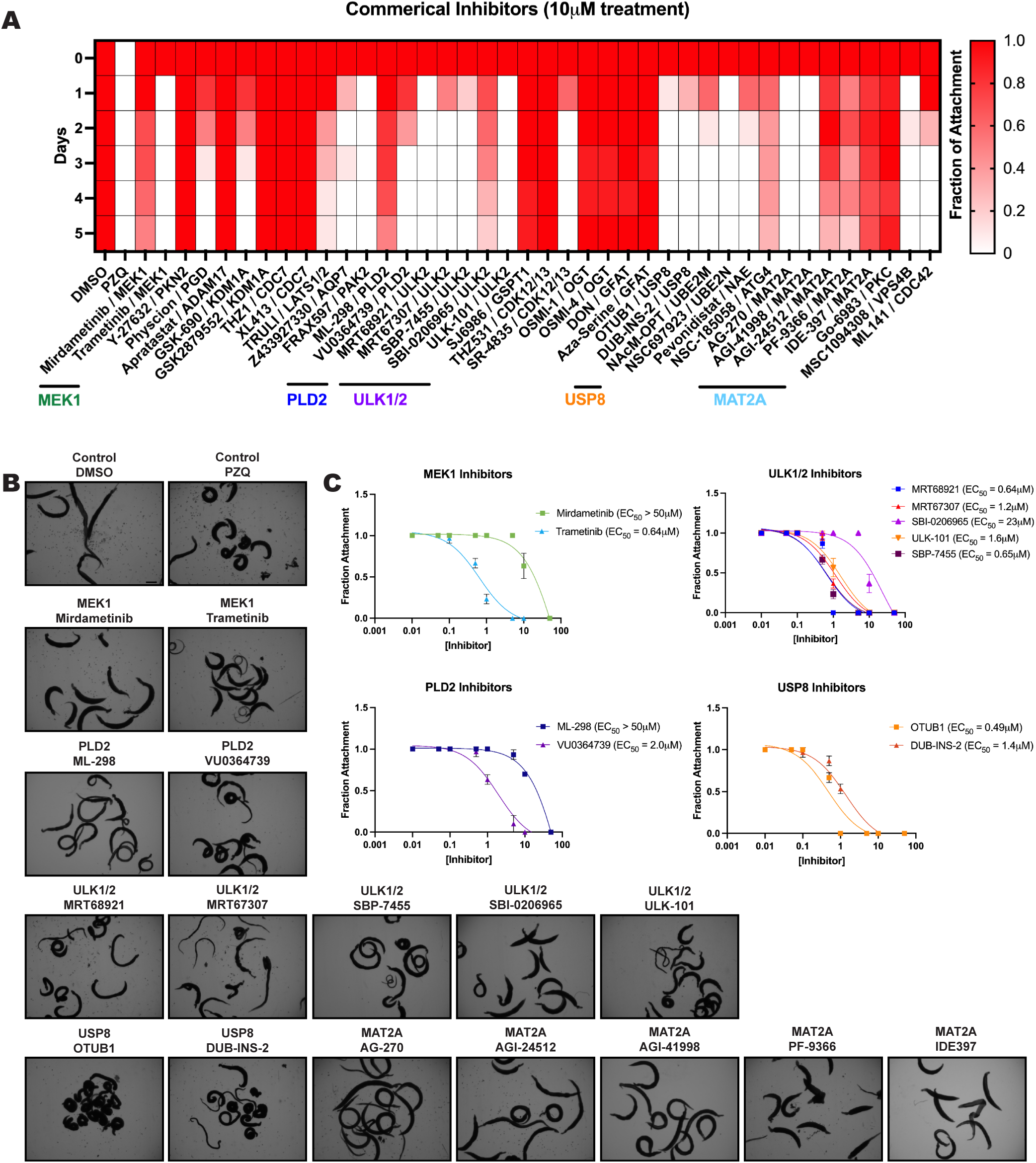
Treatment of worms with human drug target inhibitors. **(A)** Heat map showing time course following treatment of adult worms at 10 μM with commercially available inhibitors targeting human orthologs of essential genes identified in RNAi experiments (DMSO; negative control, PZQ; positive control). The fraction of a population of 10 adult worms attached to the culture plate is quantified (dark red; 1 - complete attachment of entire population, white; 0 - no attachment of any worms in population). Worms were treated with inhibitor for 72 hours, replacing drug and media every 24 hr, then monitored until the end of the experiment on day 5. **(B)** Light microscopy images of worms treated with DMSO control or reported inhibitors of human orthologs of essential schistosome genes; MEK1, PLD2, ULK1/2, USP8, and sMAT2A. **(C)** Dose-response curves of worms treated with inhibitors targeting human PLD2, MEK1, ULK1/2, and USP8. Compounds were tested from a range of 50 μM to 10 nM to determine EC_50_. Values were determined by Prism. Scale bar **(B)**, 1,000 μm.

**Supplemental Figure 3.**
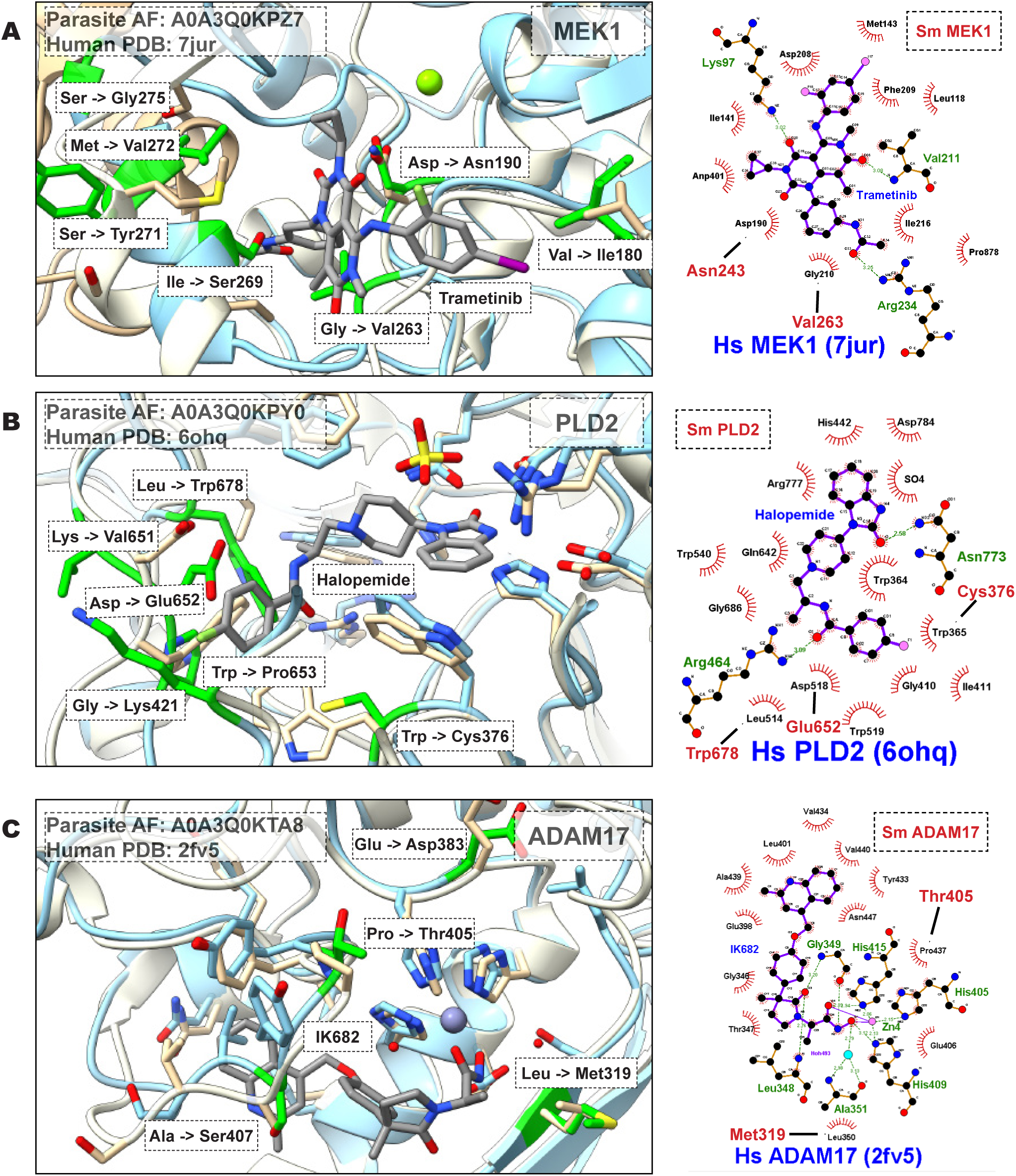
3D homology models of potential schistosome drug targets and their human orthologs. 3D alignments of predicted AlphaFold structures of schistosome (blue) proteins overlayed with structures of their closest human (white) homologs. Structures for human proteins bound to their respective inhibitors were retrieved from the Protein Data Bank for **(A)** MEK1 (PDB: 7JUR), **(B)** PLD2 (PDB: 6OHQ), and **(C)** ADAM17 (PDB: 2FV5). Unique schistosome residues are highlighted with green. Ligplots were generated using LigPlot+ v2.2 using the same PDB structures for human orthologs as listed above. Residues forming hydrophobic interactions are colored black, while other interactions are depicted in green. Unique schistosome residues are outlined in red.

**Supplemental Figure 4.**
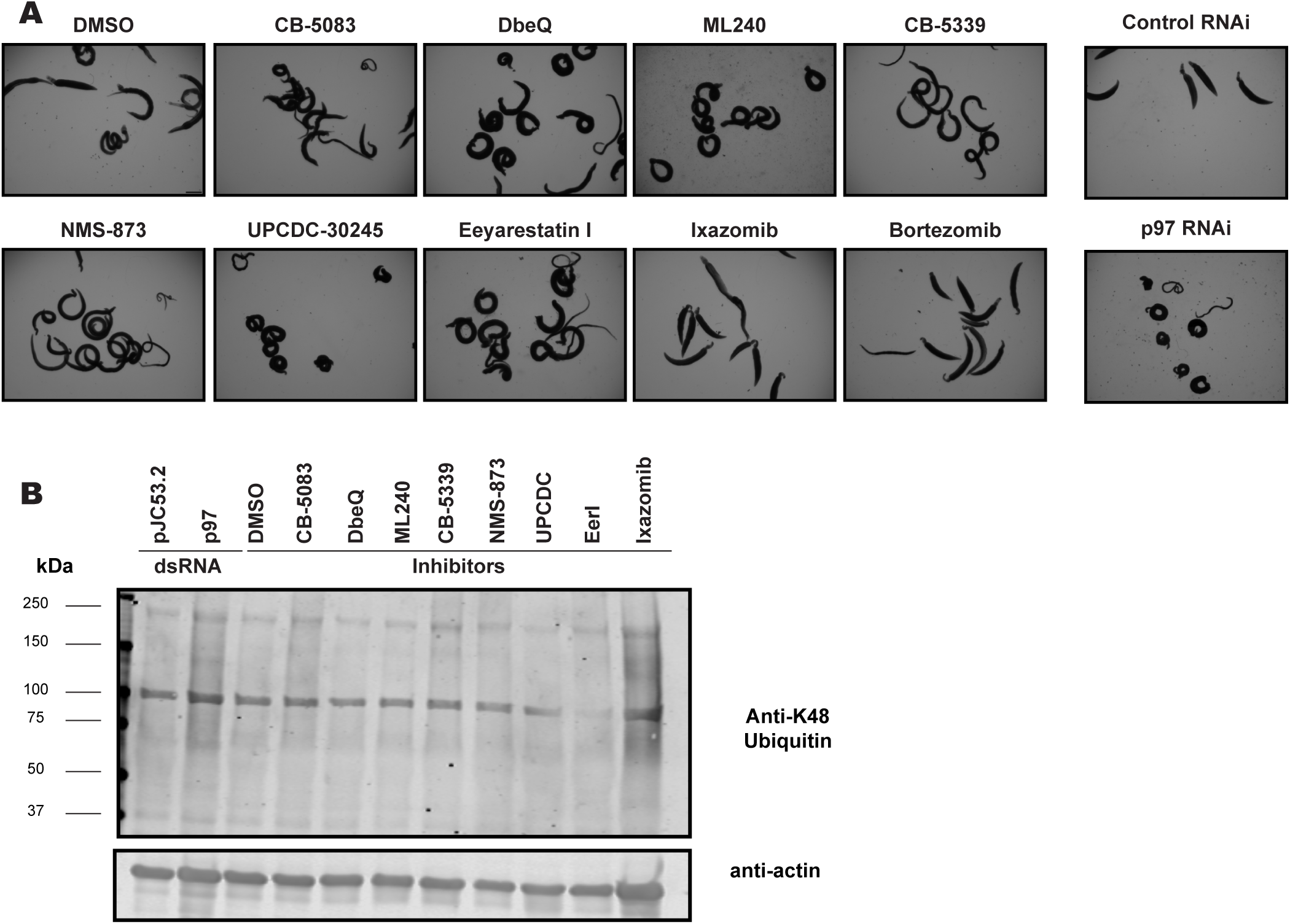
Treatment of adult parasites with known human p97 inhibitors. **(A)** Light microscopy images of adult parasites treated with either dsRNA targeting p97 (pJC53.2 control) or human p97 inhibitors (DMSO control) at 10 μM. **(B)** Western blot depicting polyubiquitinated protein profile (K48 antibody) in worm lysate following treatment of adult worms with p97 dsRNA or inhibitors (actin loading control). Scale bar **(A)**, 1,000 μm.

**Supplemental Figure 5.**
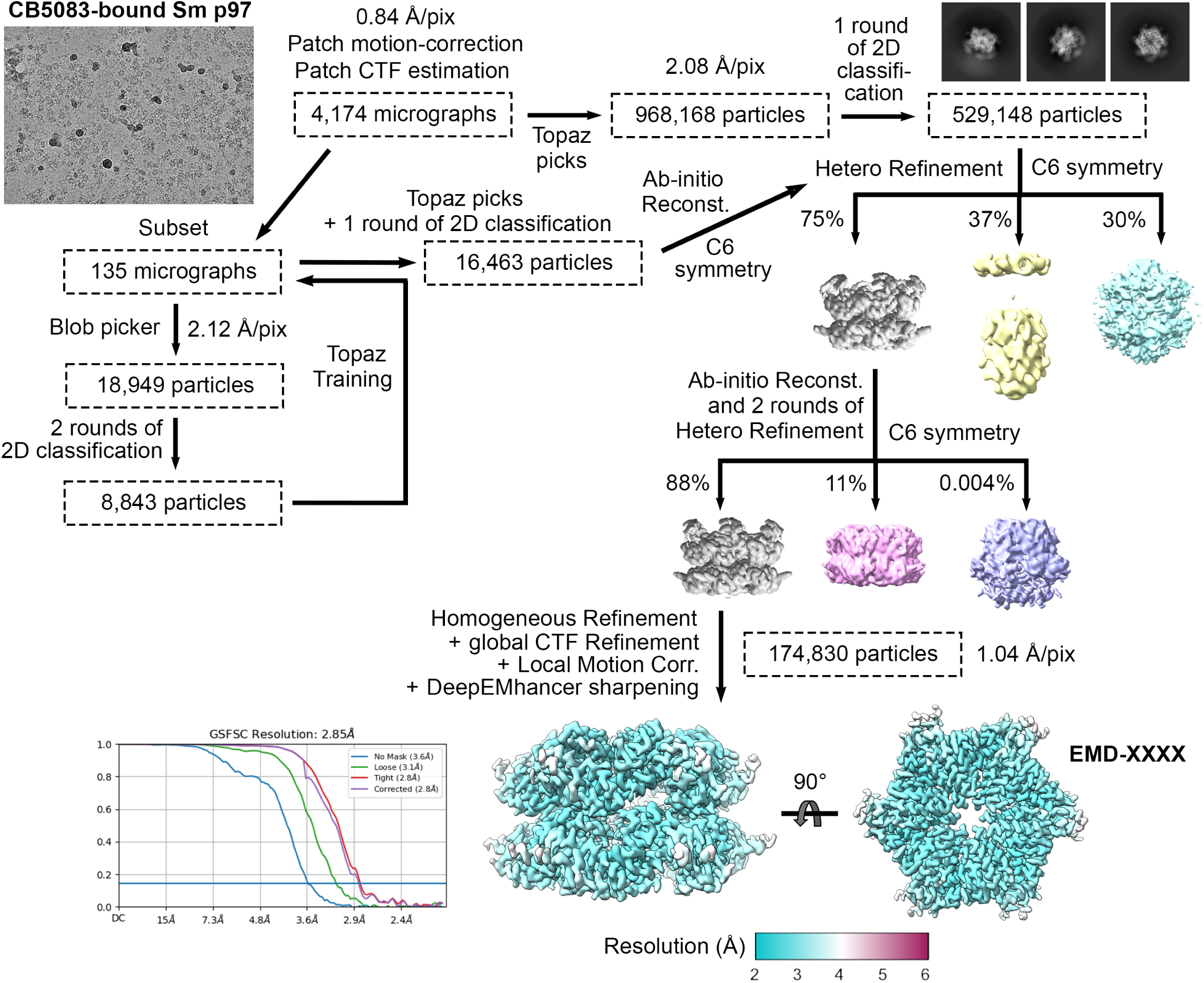
Data processing of *S. mansoni* p97 bound to CB-5083. Data processing scheme for cryo-EM dataset involving *S. mansoni* p97 bound to active site inhibitor CB-5083.

**Supplemental Figure 6.**
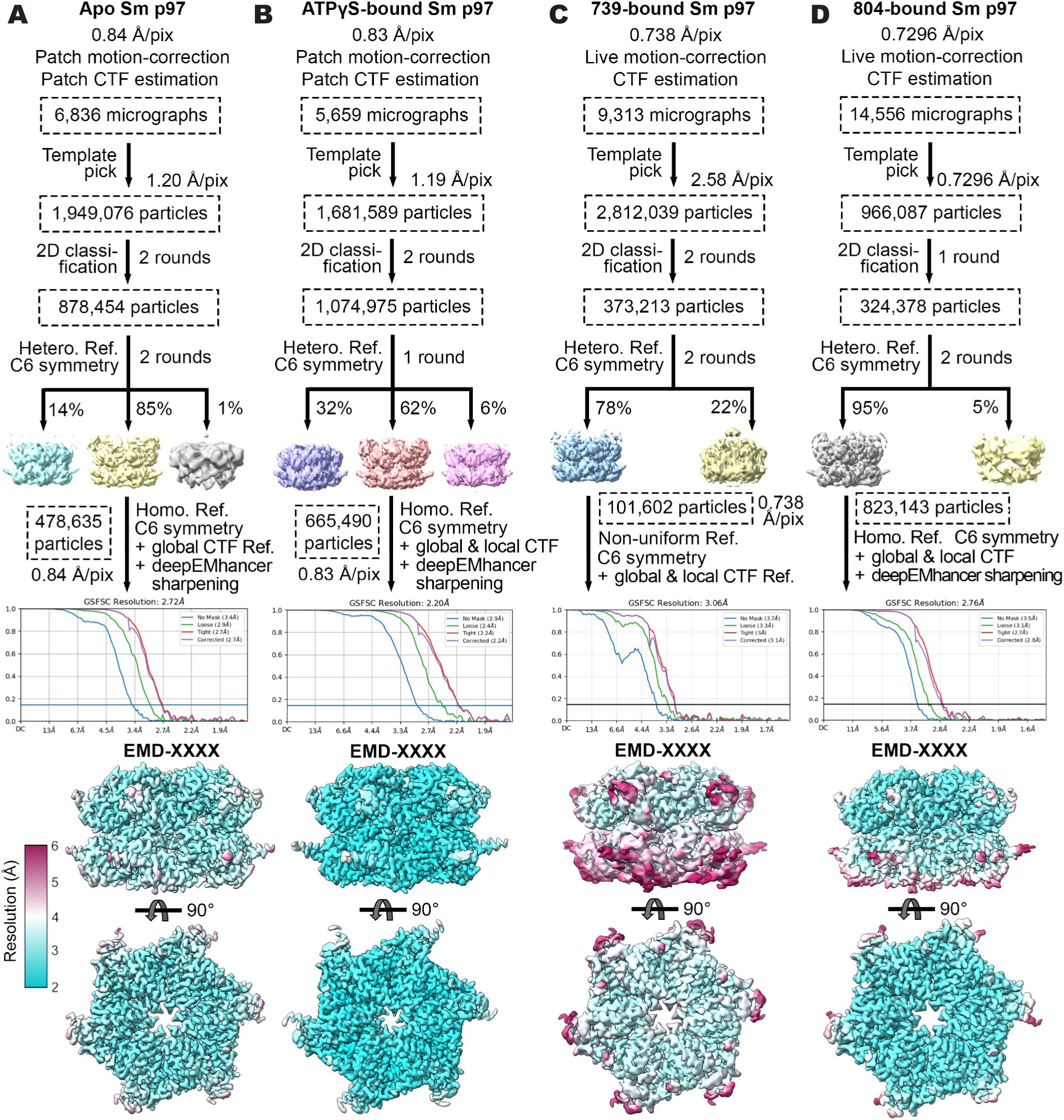
Data processing of *S. mansoni* p97 apo enzyme, and bound to ATPγS and covalent inhibitor analogs 739 and 804. **(A-D)** Data processing scheme for cryo-EM datasets involving *S. mansoni* p97 **(A)** apo enzyme and bound to **(B)** ATPγS and covalent inhibitors, **(C)** 739 and **(D)** 804.

**Supplemental Figure 7.**
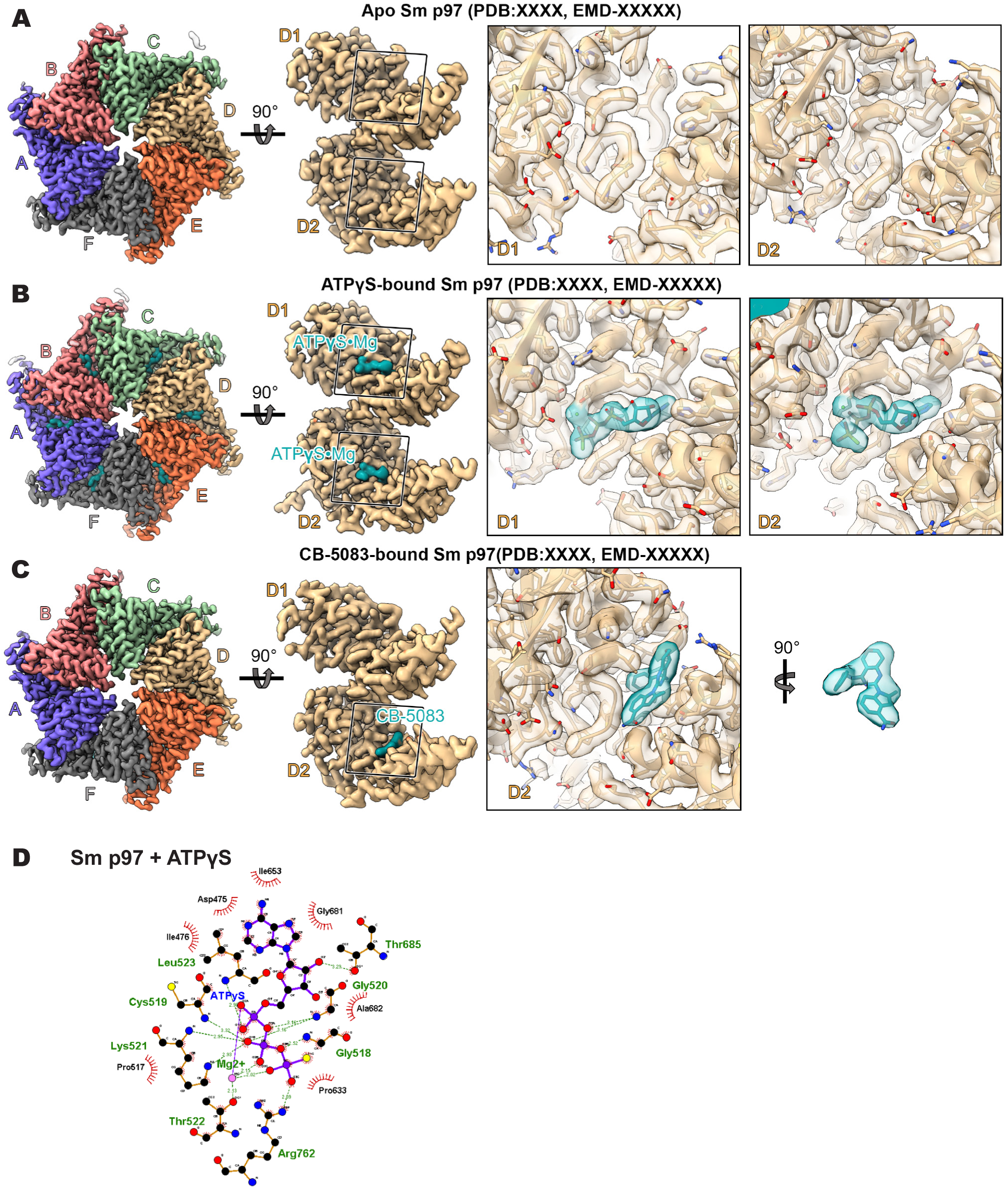
Cryo-EM map and structure of *S. mansoni* p97 apo enzyme and bound to known ligands, ATPγS and CB-5083. **(A-C)** Cryo-EM map of hexamer of schistosome p97 **(A)** apo enzyme or bound to **(B)** ATPγS or **(C)** CB-5083, colored by the final structure. Zoom of ATP binding pocket in the D1 and D2 domain of apo *S. mansoni* p97 (chain D). Boxes on the right show the map quality of the two nucleotide binding pockets. Density for CB-5083 is shown in two views. **(D)** Ligplot of residues involved in *S. mansoni* p97 binding to ATPγS in the D2 domain. Residues forming hydrophobic interactions are colored black, while other interactions are depicted in green.

**Supplemental Figure 8.**
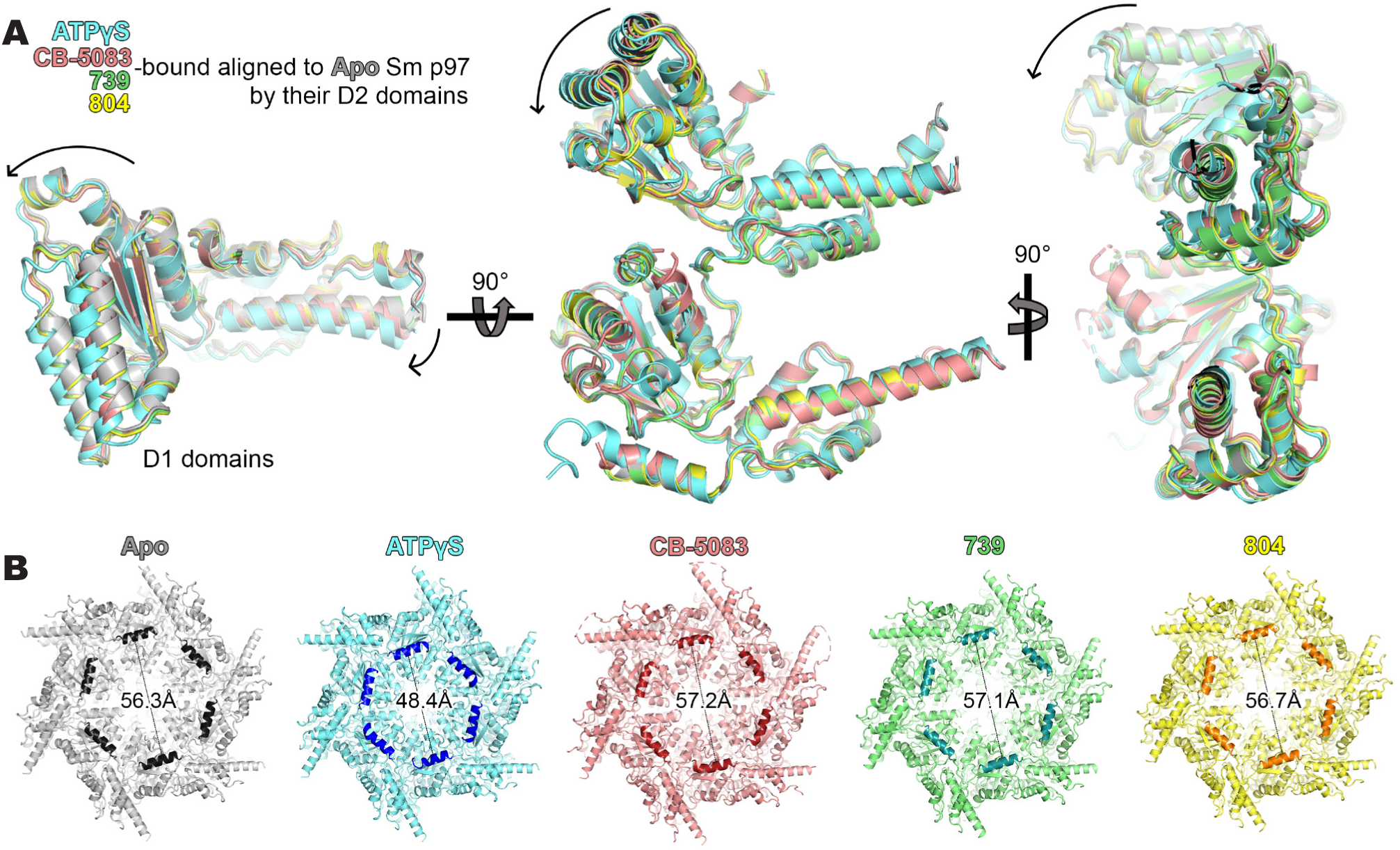
Conformational changes induced in *S. mansoni* p97 following ligand binding. Depiction of schistosome p97 conformational changes following binding to nucleotide substrate (ATPγS) and small molecule inhibitors (CB-5083, 739, 804) **(A)** Overlay of *S. mansoni* D1 domains following alignment to apo-enzyme D2 domain. **(B)** Bottom-up view of the *S. mansoni* p97 hexamer. Measurement of the diameter of the central pore of the schistosome p97 between residue K750 of two opposite chains in helix 750-757 (bold) for apo enzyme (grey) compared to ligand-bound states (blue; ATPγS, pink; CB-5083, green; 739, yellow; 804).

**Supplemental Figure 9.**
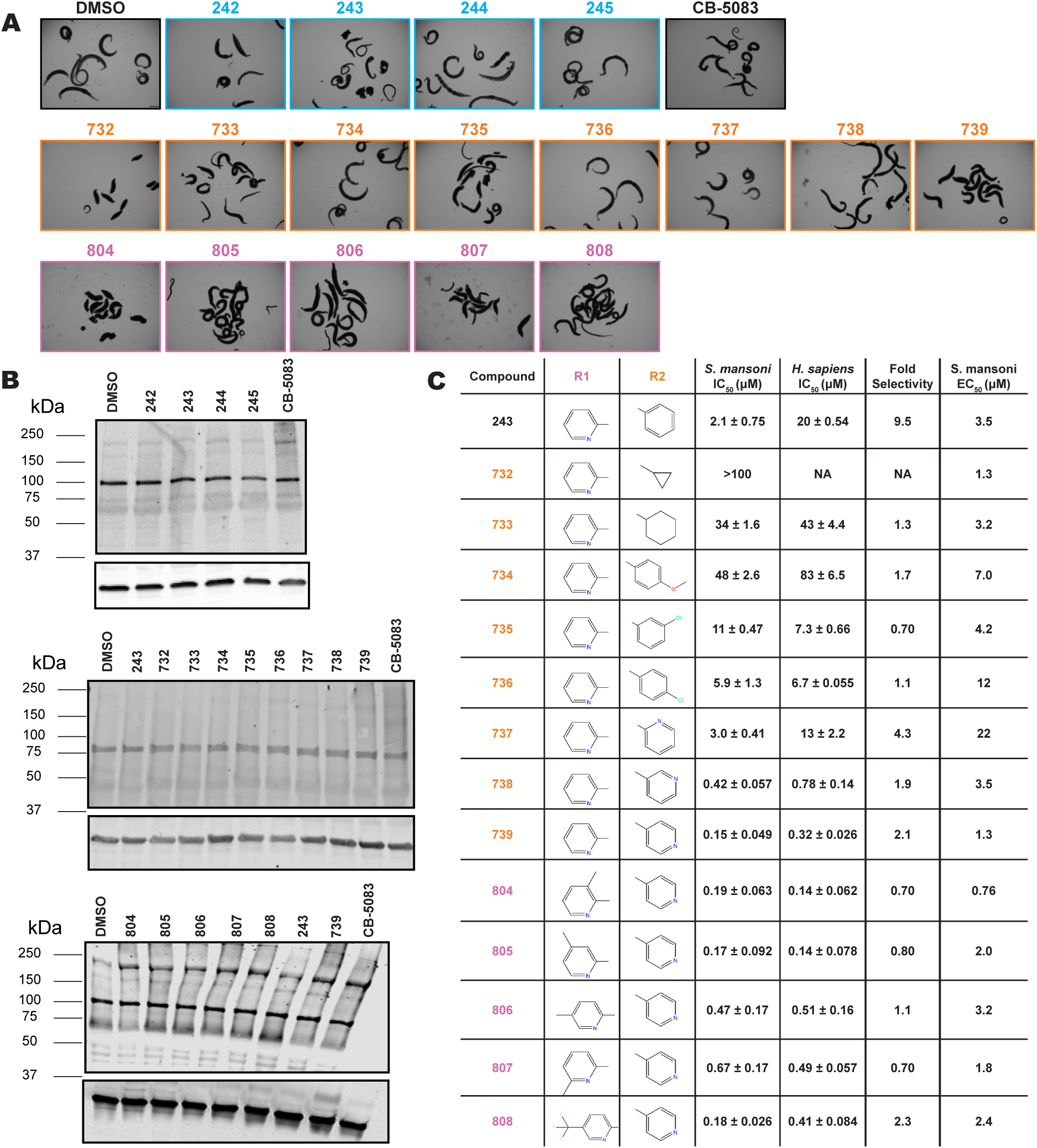
Treatment of adult parasites with analogs of covalent p97 scaffold identified in high-throughput screen and known human p97 inhibitors. **(A)** Light microscopy images of adult parasites treated with controls (negative; DMSO, positive; CB-5083) and analogs of the covalent p97 inhibitor scaffold identified in high-throughput screen. **(B)** Western blot depicting polyubiquitinated protein profile (K48 antibody) in worm lysate following treatment by DMSO control or p97 covalent inhibitor analogs (actin loading control). **(C)** Full structural activity relationship modifications of lead compound series outlining **R1** (800 series) and **R2** (700 series) modifications made to the depicted scaffold. Comparative IC_50_ values display the potency of each compound on the recombinant parasite (*Sm* p97) and human (*Hs* p97) enzyme. Compounds were tested from 100 μM – 1 nM. Values were calculated using Prism. EC_50_ values for benzoxazole propiolamide scaffold analogs on adult parasites determined by fraction of worms attached to tissue culture plate following 72 hours of drug treatment, refreshing media and drug every 24 hr, then allowing worms to remain in culture until D5 (50 μM - 10 nM) (negative; DMSO, positive; CB-5083). Values were calculated using Prism. Scale bar (A), 1,000 μm.

**Supplemental Figure 10.**
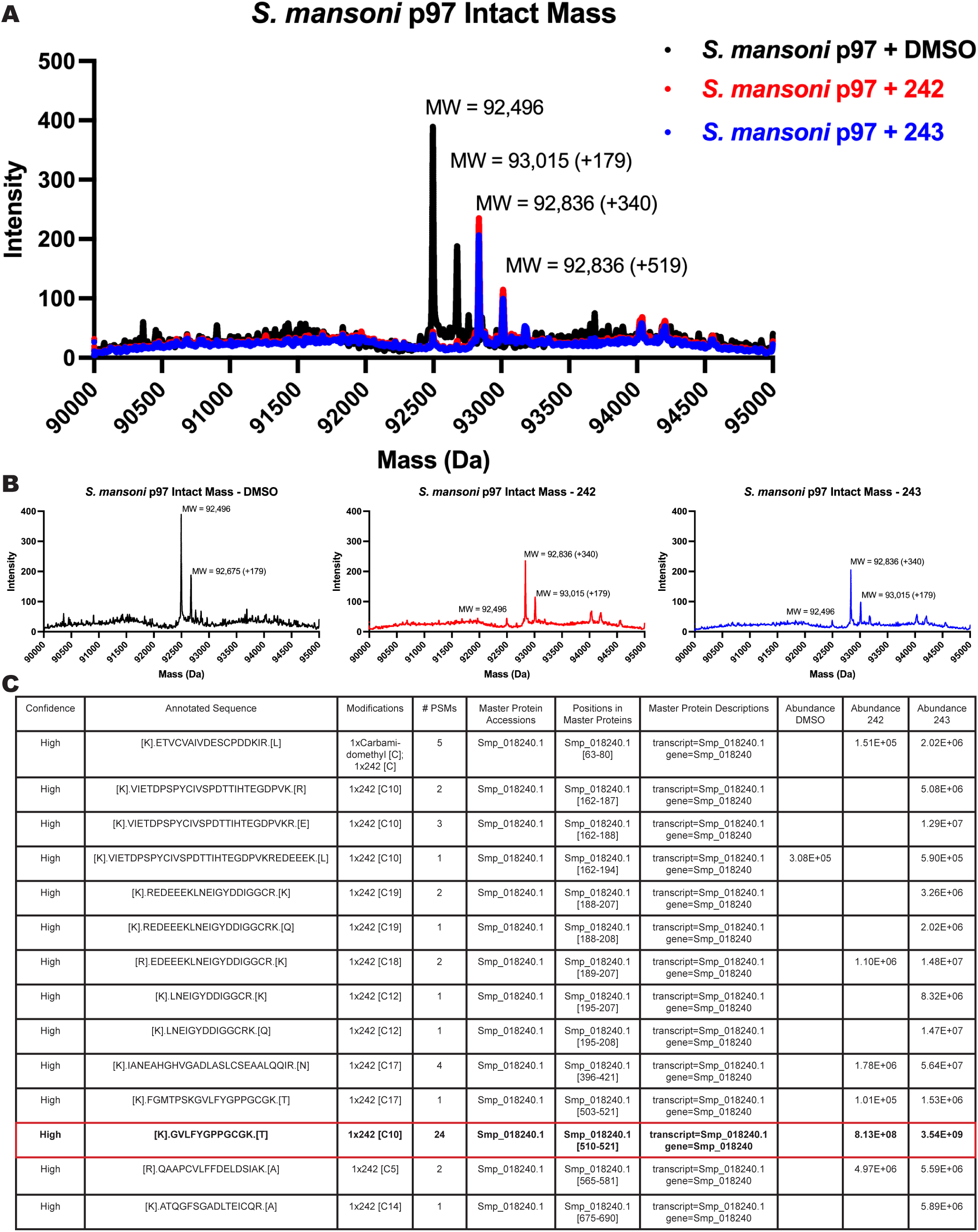
Mass spectrometry of covalent compounds in complex with schistosome p97. **(A)** Intact mass spectrum of recombinant *S. mansoni* p97 in solution with DMSO control (black) or covalent scaffold compounds (242 - red and 243 - blue). Major peaks consist of a single p97 monomer (~92.5 kDa) and an additional isoform (+179 Da). Mass shifts seen following incubation with either covalent inhibitor (339.35 Da) or DMSO. **(B)** Isolated mass spectrum of recombinant *S. mansoni* p97 in solution with DMSO control (black) or covalent scaffold compounds (242 - red and 243 - blue). **(C)** Table outlining detected peptides following incubation of schistosome p97 with covalent compound 242 or 243. Resulting reaction mixture was run on an SDS-PAGE, then the corresponding band was isolated and submitted for trypsin digest and peptide identification by LC-MS.

**Supplemental Figure 11.**
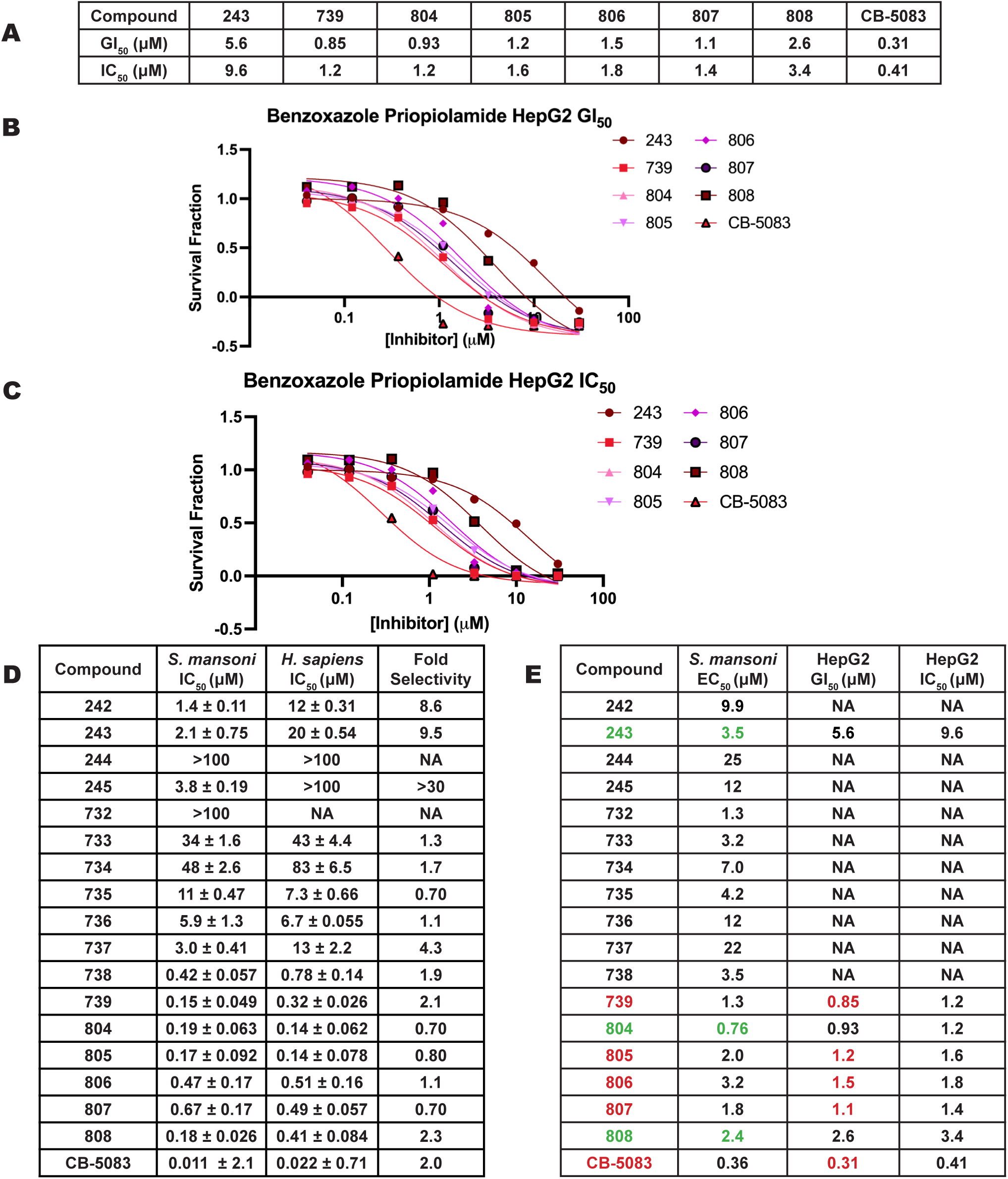
Covalent p97 inhibitor series HepG2 toxicity studies. **(A)** Table of GI_50_ (50% growth inhibition normalized to time = 0) and IC_50_ (50% growth inhibition relative to control) values for HepG2 cells treated by covalent p97 inhibitor scaffold compared to CB-5083 control. Values were plotted in excel and the 50% point was determined by interpolation. **(B)** Graphs visualized in Prism depicting GI_50_ and **(C)** IC_50_ curves for HepG2 cytotoxicity experiment following treatment by covalent analogs. Table comparing potencies on **(D)** p97 enzyme (schistosome and human) and **(E)** worms and human cells. Values were calculated in Prism.

**Supplemental Figure 12.**
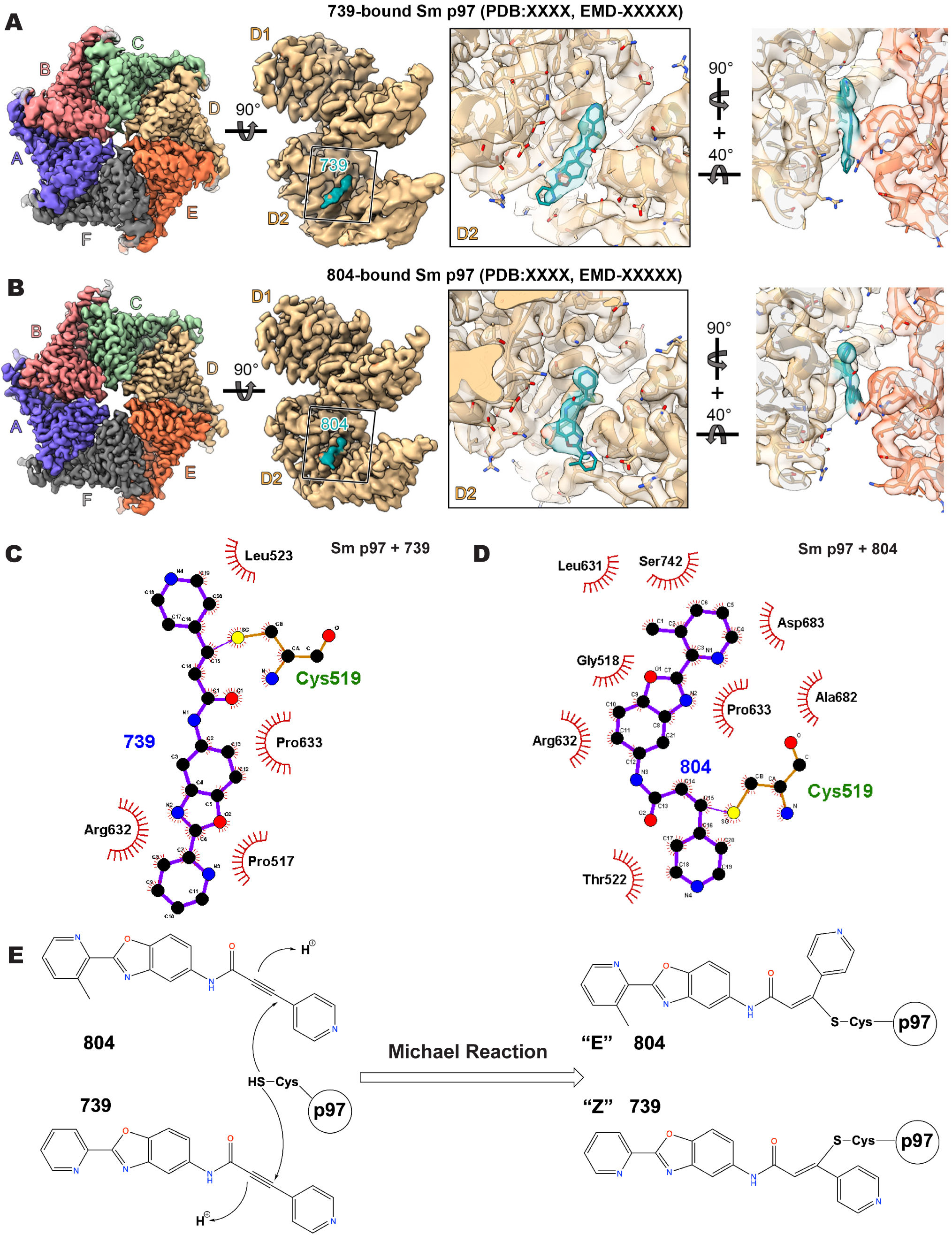
Cryo-EM Structure of *S. mansoni* p97 - Compound 739 or 804 complex. (**A-B**) Cryo-EM map of the schistosome p97 bound to compound **(A)** 739 and **(B)** 804, colored by the final structures. Two zoom-in views of the D2 domain of *S. mansoni* p97 is shown on the right. (**C-D**) Ligplot of residues involved in binding to **(C)** 739 and **(D)** 804. **(E)** Schematic depicting Michael addition reaction conducted by the thiol side chain of Cys519 in the schistosome p97 resulting in “E-“ and “Z-“ olefin configuration liganded states for compounds 804 and 739, respectively.

