## Supplemental Table 6 for "A genome-scale drug discovery pipeline uncovers new therapeutic targets and a unique p97 allosteric binding site in *Schistosoma mansoni*"

**Supplemental Table 6. Cryo-EM data collection and model statistics.**

|  | **Apo**  **Sm p97** | **Sm p97-ATPγS** | **Sm p97-CB5083** | **Sm p97-739** | **Sm p97-804** |
| --- | --- | --- | --- | --- | --- |
| **Data collection and processing** | | | | | |
| Magnification | 105 kX | 105 kX | 105 kX | 165 kX | 165 kX |
| Voltage (kV) | 300 | 300 | 300 | 300 | 300 |
| Electron exposure (e^–^/Å^2^) | 50 | 60 | 62 | 60 | 50 |
| Defocus range (μm) | -0.9 to -2.2 | -0.9 to -2.2 | -1.0 to -2.4 | -0.9 to -2.2 | -0.9 to -2.2 |
| Pixel size (Å) | 0.84 | 0.83 | 0.83 | 0.738 | 0.7296 |
| Symmetry imposed | C6 | C6 | C6 | C6 | C6 |
| Initial particle images (no.) | 1,949,076 | 1,681,589 | 968,168 | 2,812,039 | 2,272,397 |
| Final particle images (no.) | 478,635 | 665,490 | 174,830 | 101,602 | 823,143 |
| Map resolution (Å) | 2.72 | 2.2 | 2.85 | 3.07 | 2.76 |
| FSC threshold | 0.143 | 0.143 | 0.143 | 0.143 | 0.143 |
| **Refinement** | | | | | |
| Initial model used | CB5083-structure | Apo structure | Model  Angelo | Apo structure | Apo structure |
| Model composition |  |  |  |  |  |
| Non-H atoms | 24929 | 25626 | 25437 | 24664 | 24626 |
| Protein residues | 3183 | 3234 | 3230 | 3133 | 3127 |
| Ligand | 0 | MG: 12  AGS: 12 | JDP: 6 | I73: 6 | I80: 6 |
| R.m.s. deviations |  |  |  |  |  |
| Bond lengths (Å) | 0.004 | 0.003 | 0.002 | 0.006 | 0.003 |
| Bond angles (°) | 0.494 | 0.465 | 0.434 | 0.537 | 0.517 |
| CC (volume/mask) | 0.82/0.82 | 0.83/0.82 | 0.78/0.78 | 0.82/0.83 | 0.72/0.71 |
| CC for ligands | - | 0.84 | 0.77 | 0.79 | 0.46 |
| **Validation** | | | | | |
| MolProbity score | 1.27 | 1.12 | 1.34 | 1.47 | 1.39 |
| Clashscore | 3.26 | 3.33 | 5.09 | 4.88 | 5.98 |
| Poor rotamers (%) | 0.04 | 0.00 | 0.00 | 0.00 | 0.00 |
| Ramachandran plot |  |  |  |  |  |
| Favored (%) | 97.15 | 98.14 | 97.70 | 96.62 | 97.72 |
| Allowed (%) | 2.85 | 1.86 | 2.30 | 3.38 | 2.28 |
| Disallowed (%) | 0.00 | 0.00 | 0.00 | 0.00 | 0.00 |
| Protein residues included in the model | A: 192-428, 434-546, 555-581, 593-708, 722-758  B: 193-427, 434-546, 555-581, 593-709, 722-758  C: 192-427, 434-546, 555-581, 593-709, 722-758  D: 193-428, 433-546, 555-581, 593-709, 722-758  E: 193-427, 434-546, 555-581, 594-709, 722-758  F: 191-425, 434-546, 555-581, 593-709, 722-758 | A/B/C/D/E/F: 196-427, 434-547, 554-582, 593-709, 722-768 | A: 193-427, 434-550, 554-582, 593-711, 720-758  B: 193-427, 434-550, 554-582, 593-710, 720-758  C: 193-427, 435-550, 554-582, 592-709, 720-758  D: 193-427, 434-550, 554-582, 592-709, 720-758  E: 193-427, 434-550, 554-582, 592-708, 720-758  F: 193-427, 434-550, 554-582, 593-713, 720-758 | A: 194-426, 435-546, 555-580, 595-706, 722-758  B: 194-425, 435-550, 554-580, 595-707, 722-758  C: 194-427, 434-546, 554-581, 594-707, 722-758  D: 193-426, 436-546, 554-581, 594-707, 722-758  E: 194-426, 436-546, 554-579, 595-707, 722-758  F: 193-425, 435-546, 554-580, 595-707, 722-758 | A: 192-424, 436-546, 555-581, 595-706, 722-758  B: 193-424, 436-546, 554-581, 595-706, 722-758  C: 192-424, 436-546, 554-582, 595-706, 722-758  D: 193-424, 436-546, 554-581, 595-706, 722-758  E: 193-424, 435-546, 554-581, 594-706, 722-758  F: 191-424, 435-546, 554-581, 595-706, 722-758 |
| PDB/EMDB code | XXXX/EMD-XXXXX | XXXX/EMD-XXXXX | XXXX/EMD-XXXXX | XXXX/EMD-XXXXX | XXXX/EMD-XXXXX |
