## Supplemental Table 7 for "A genome-scale drug discovery pipeline uncovers new therapeutic targets and a unique p97 allosteric binding site in *Schistosoma mansoni*"

| Category | Parameter | Description |
| --- | --- | --- |
| Assay | Type of assay | *In vitro* ATP consumption (Kinase-Glo) |
|  | Target | *S. mansoni* vcp/p97 (Smp_018240) |
|  | Primary measurement | Luminescent signal compared to DMSO or known inhibitor (CB-5083) control to indicate inhibition of p97’s ATPase activity |
|  | Key reagents | Full-length (1-804aa) 6xHis *S. mansoni* p97, Promega Kinase-Glo, PerkinElmer 384-well plates |
|  | Assay protocol | Described in Methods section |
|  | Additional comments | None |
| Library | Library size | ~350,000 compounds; 1 compound/well |
|  | Library composition | Synthetic small molecules and partially purified natural product fractions |
|  | Source | ChemDiv (150,000 compounds), ChemBridge (125,500), ComGenex (22,000), Preswick Chemical (1,100), TimTec (500), UT Southwestern Chemistry in-house collection (2,500), Dr. John Macmillan (UC Santa Cruz) natural product collection (7,100 partially purified fractions) |
|  | Additional Comments |  |
| Screen | Format | 384-well plates |
|  | Concentration(s) tested | 10 μM compound concentration; 0.2% DMSO |
|  | Plate controls | Negative control (0.2% DMSO); Positive control (10 μM CB-5083); Maximal signal (10 μM ATP) |
|  | Assay validation / Quality Control | Average plate Z’value = 0.673 |
|  | Correction factors | None |
|  | Normalization | Normalized to internal inhibitor control (CB-5083) |
|  | Additional comments | Screened at UTSW Medical Center HTS Facility |
| Post-HTS Analysis | Hit criteria | >25% enzyme activity inhibition |
|  | Hit rate | 0.2% (655 of 350,000) in initial screen. Each hit was re-assayed in quadruplicate using an 8-point dose-response curve (0.00614-15 μM) |
|  | Additional assay(s) | Counter-screened each hit vs. purified recombinant full-length (1-806aa) *H. sapiens* p97 to determine preliminary selectivity using an 8-point dose-response curve (0.00614-15 μM) in quadruplicate |
|  | Confirmation of hit purity and structure |  |
|  | Additional comments | Analysis performed at UTSW Medical Center by HTS Facility. Dr. Joseph Ready (UTSW Chemistry) assisted with removal of PAINS and promiscuous compounds. |

**Supplementary Table 7. p97 high-throughput screen pipeline**
