## Supplemental Materials Chemical Synthesis for "A genome-scale drug discovery pipeline uncovers new therapeutic targets and a unique p97 allosteric binding site in *Schistosoma mansoni*"

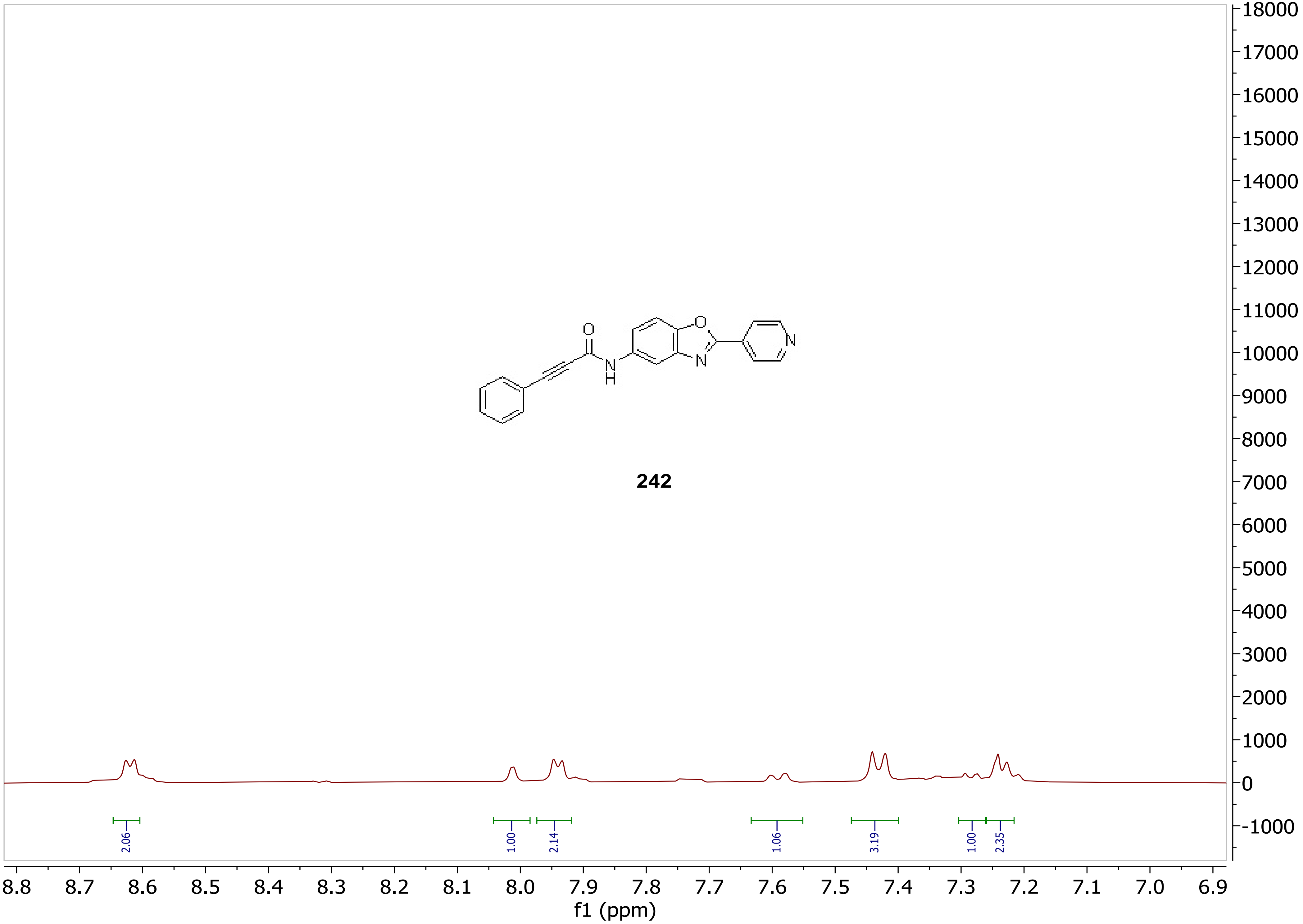

242
O
N
H
N
2.06
1.00
2.14
1.06
3.19
1.00
2.35
8.8 8.7 8.6 8.5 8.4 8.3 8.2 8.1 8.0 7.9 7.8 7.7 7.6 7.5 7.4 7.3 7.2 7.1 7.0 6.9
f1 (ppm)
18000 17000 16000 15000 14000 13000 12000 11000 10000 9000 8000 7000 6000 5000 4000 3000 2000 1000 0 -1000

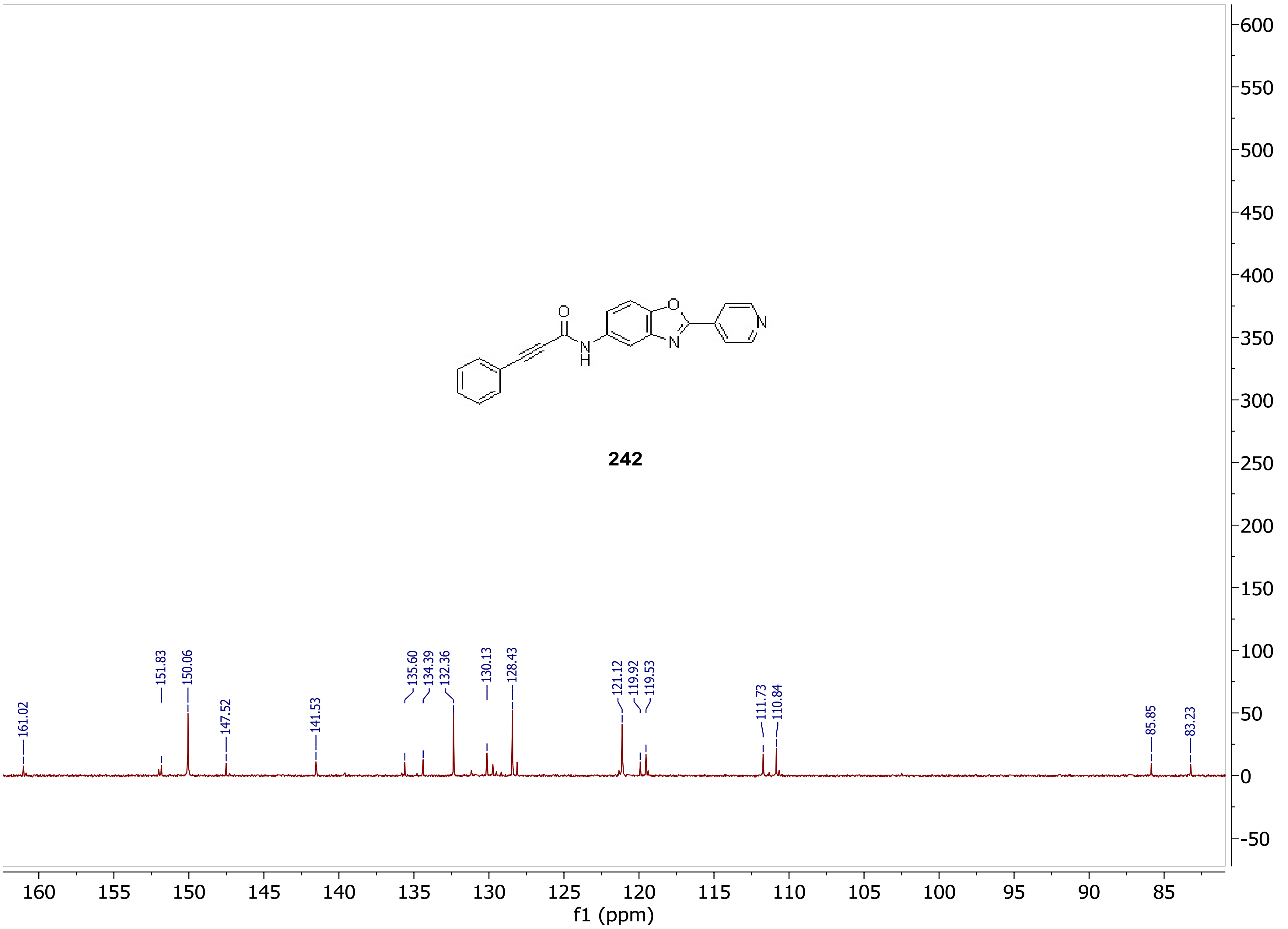

242
161.02
151.83
150.06
147.52
141.53
135.60
134.39
132.36
130.13
128.43
121.12
119.92
119.53
111.73
110.84
85.85
83.23
f1 (ppm)

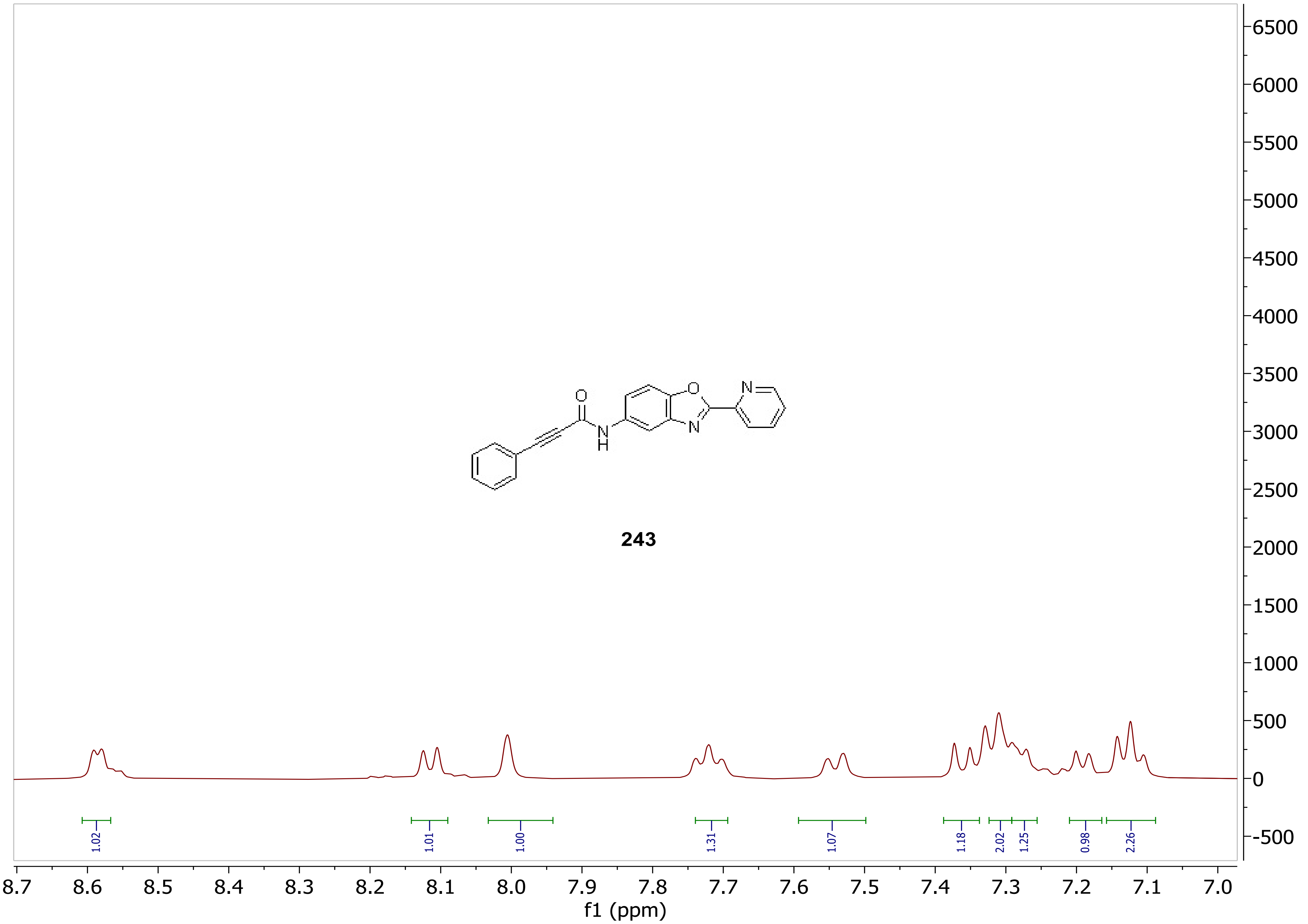
243
1.02
1.01
1.00
1.31
1.07
1.18
2.02
1.25
0.98
2.26
8.7
8.6
8.5
8.4
8.3
8.2
8.1
8.0
7.9
7.8
7.7
7.6
7.5
7.4
7.3
7.2
7.1
7.0
f1 (ppm)
6500
6000
5500
5000
4500
4000
3500
3000
2500
2000
1500
1000
500
0
-500

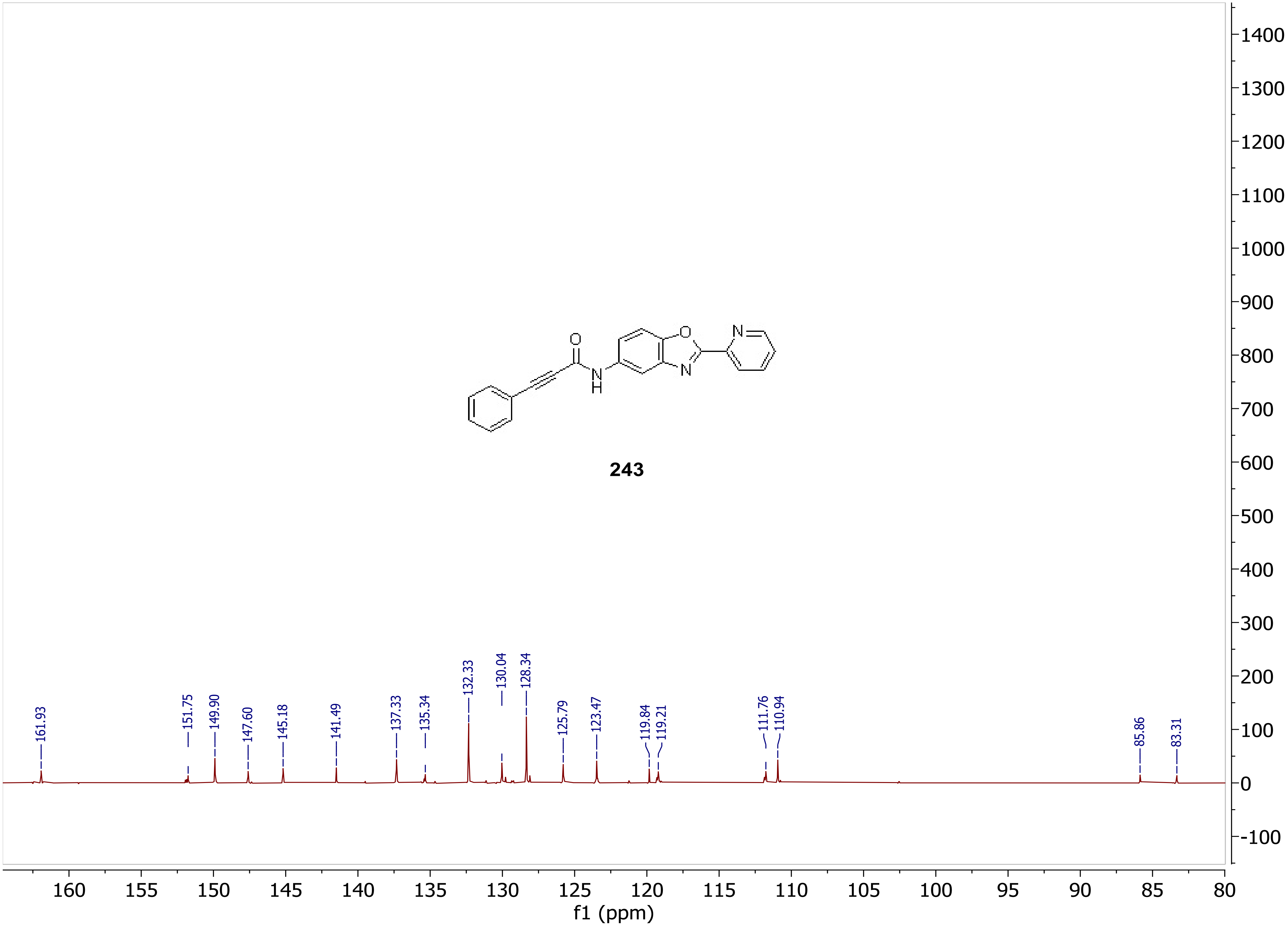
O
N
H
O
N
N
243
161.93
151.75
149.90
147.60
145.18
141.49
137.33
135.34
132.33
130.04
128.34
125.79
123.47
119.84
119.21
111.76
110.94
85.86
83.31
1400
1300
1200
1100
1000
900
800
700
600
500
400
300
200
100
0
-100
160
155
150
145
140
135
130
125
120
115
110
105
100
95
90
85
80
f1 (ppm)

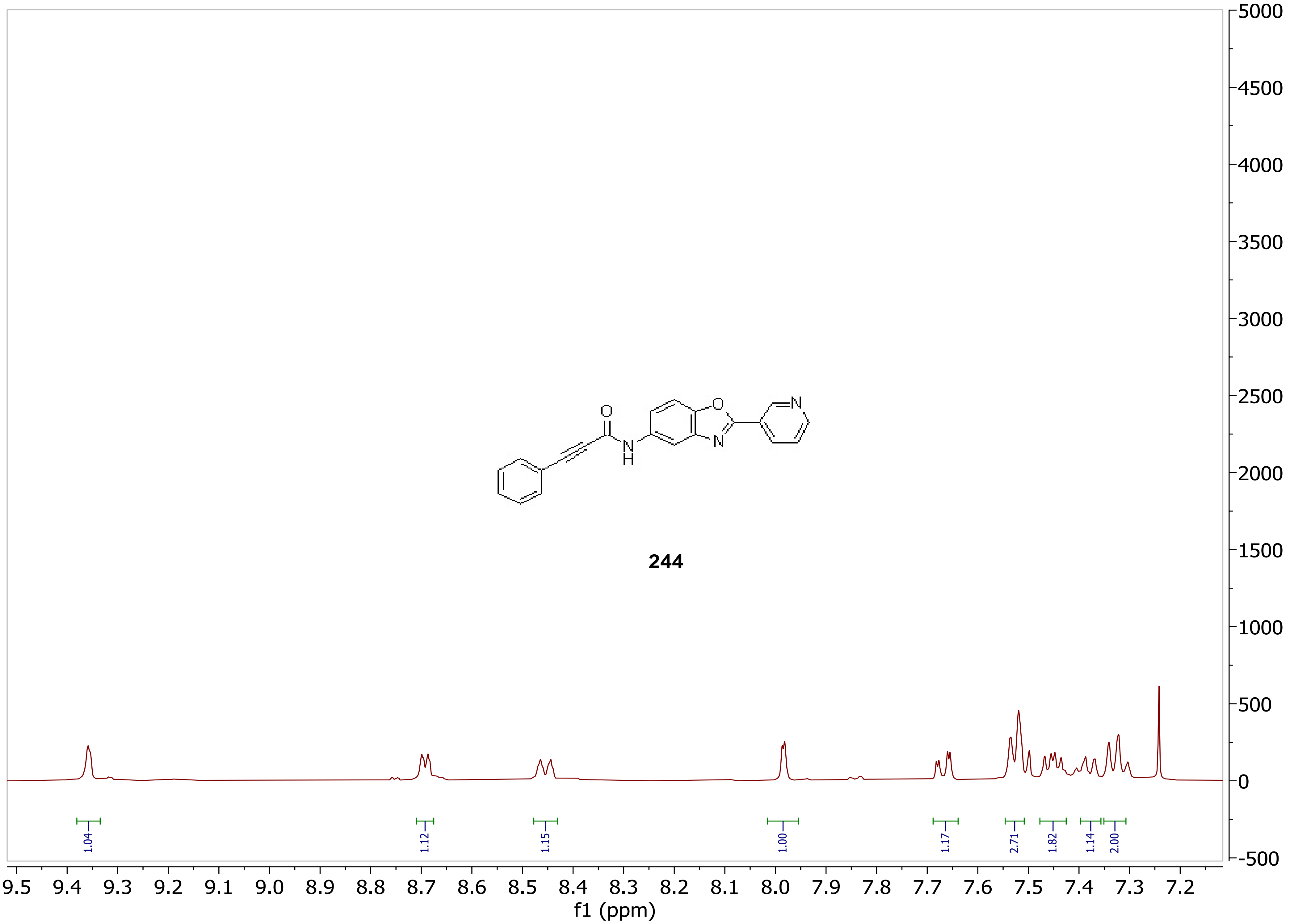

244
O
N
H
N
N
1.04
1.12
1.15
1.00
1.17
2.71
1.82
1.14
2.00
9.5 9.4 9.3 9.2 9.1 9.0 8.9 8.8 8.7 8.6 8.5 8.4 8.3 8.2 8.1 8.0 7.9 7.8 7.7 7.6 7.5 7.4 7.3 7.2
f1 (ppm)
5000 4500 4000 3500 3000 2500 2000 1500 1000 500 0 -500

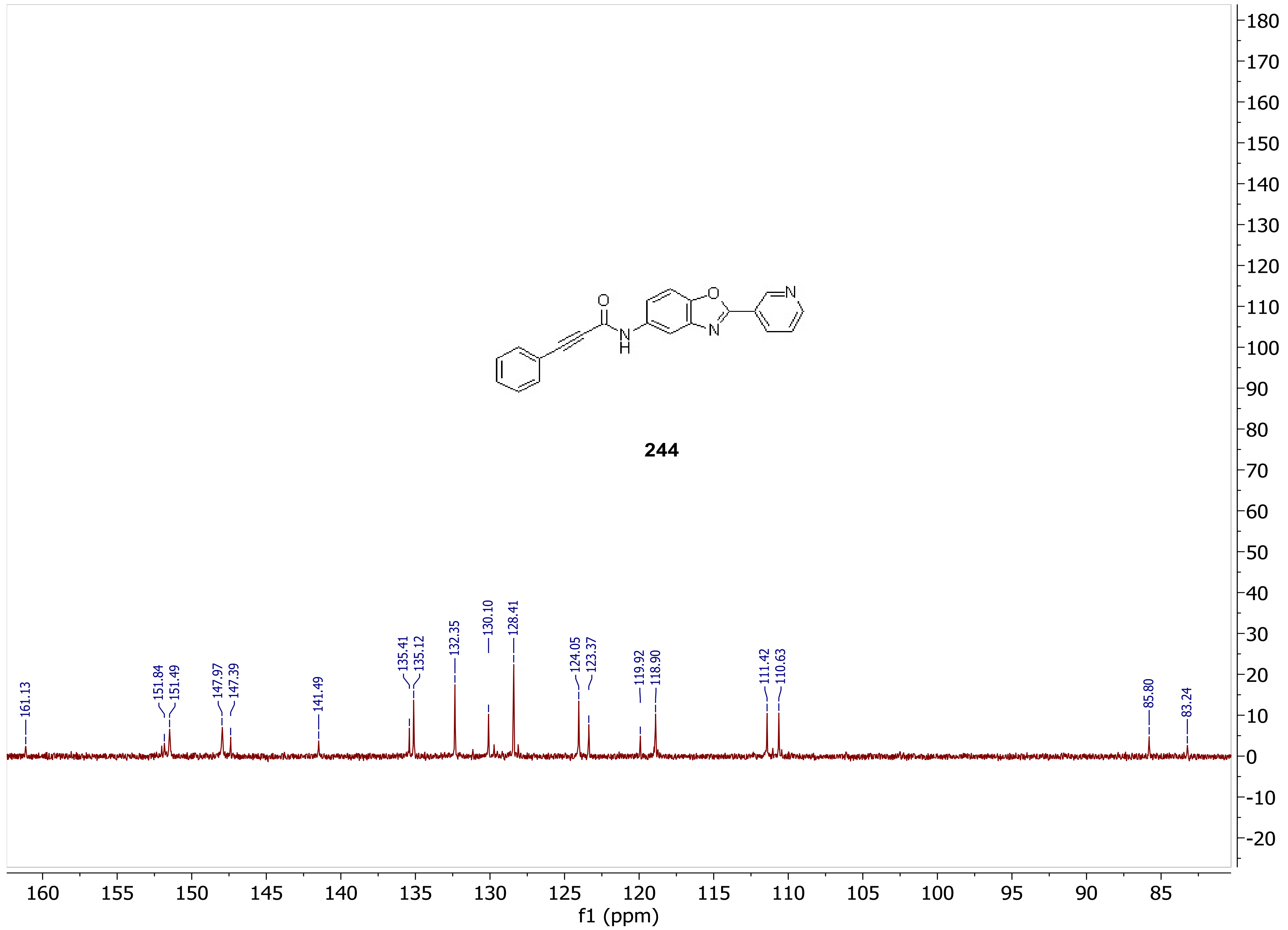

244
161.13
151.84
151.49
147.97
147.39
141.49
135.41
135.12
132.35
130.10
128.41
124.05
123.37
119.92
118.90
111.42
110.63
85.80
83.24
f1 (ppm)

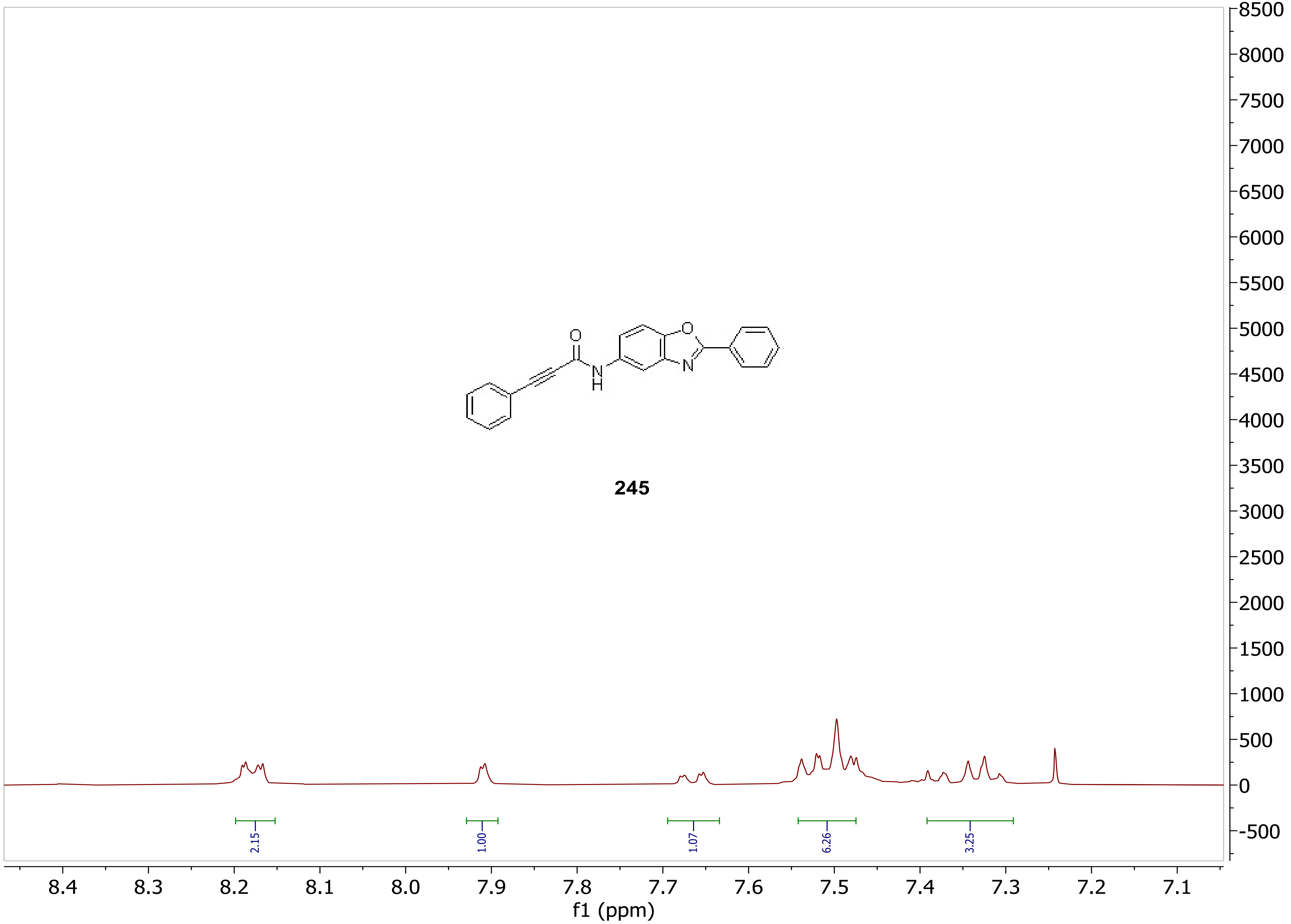
245
2.15
1.00
1.07
6.26
3.25
8.4
8.3
8.2
8.1
8.0
7.9
7.8
7.7
7.6
7.5
7.4
7.3
7.2
7.1
f1 (ppm)
8500
8000
7500
7000
6500
6000
5500
5000
4500
4000
3500
3000
2500
2000
1500
1000
500
0
-500

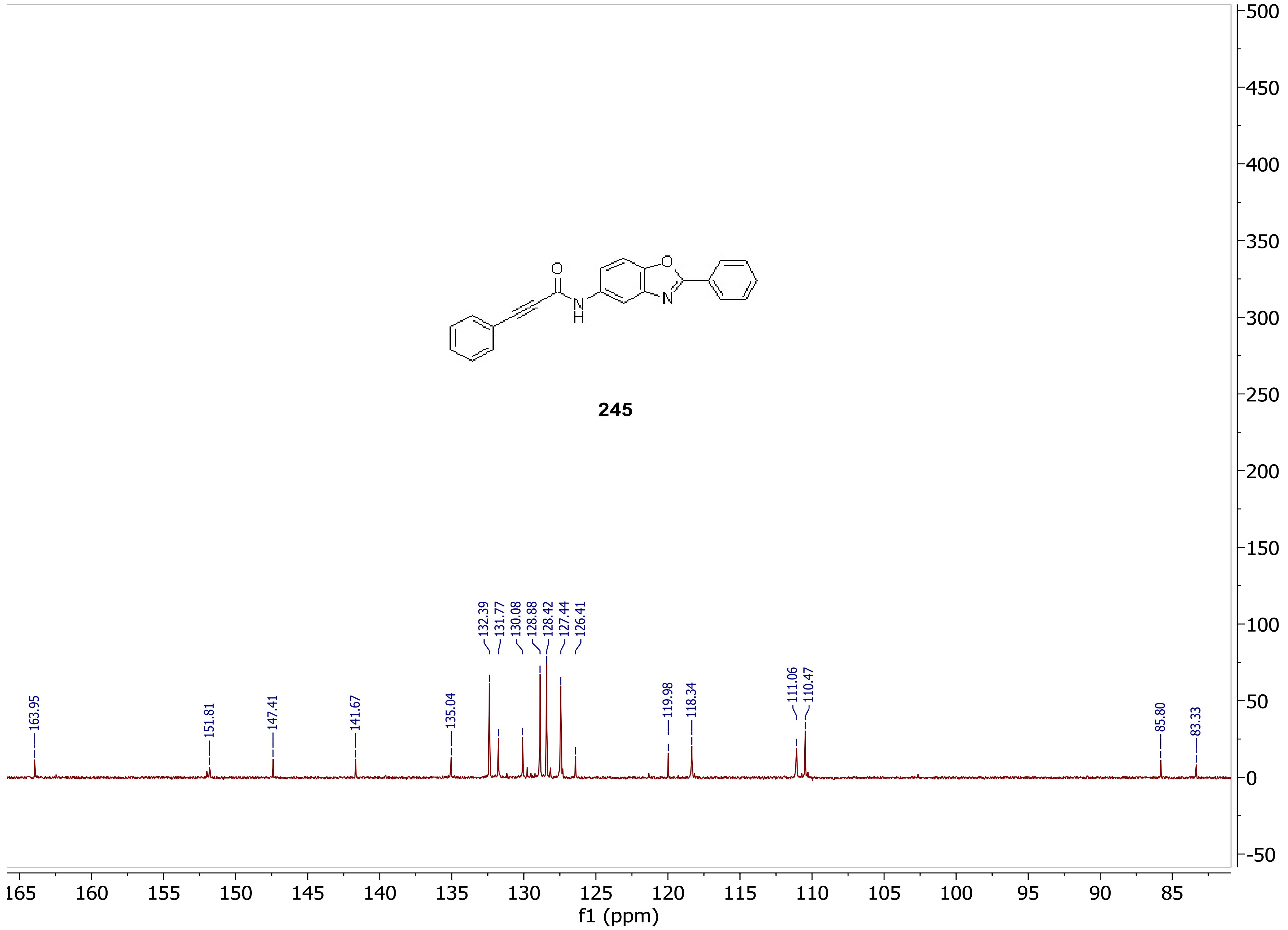
245
163.95
151.81
147.41
141.67
135.04
132.39
131.77
130.08
128.88
128.42
127.44
126.41
119.98
118.34
111.06
110.47
85.80
83.33
f1 (ppm)
165 160 155 150 145 140 135 130 125 120 115 110 105 100 95 90 85
500 450 400 350 300 250 200 150 100 50 0 -50

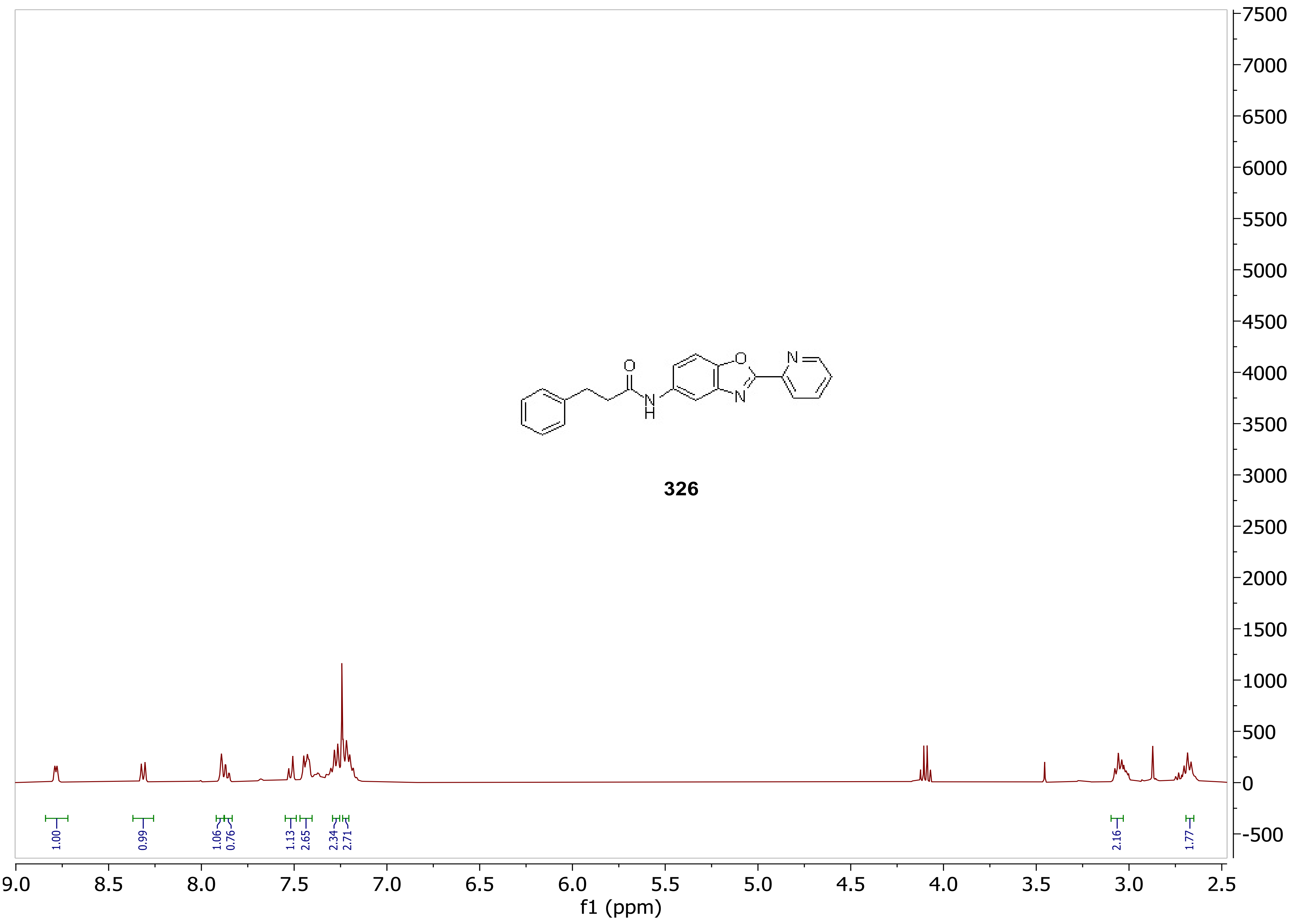
326
1.00
0.99
1.06
0.76
1.13
2.65
2.34
2.71
2.16
1.77
9.0
8.5
8.0
7.5
7.0
6.5
6.0
5.5
5.0
4.5
4.0
3.5
3.0
2.5
f1 (ppm)
7500
7000
6500
6000
5500
5000
4500
4000
3500
3000
2500
2000
1500
1000
500
0
-500

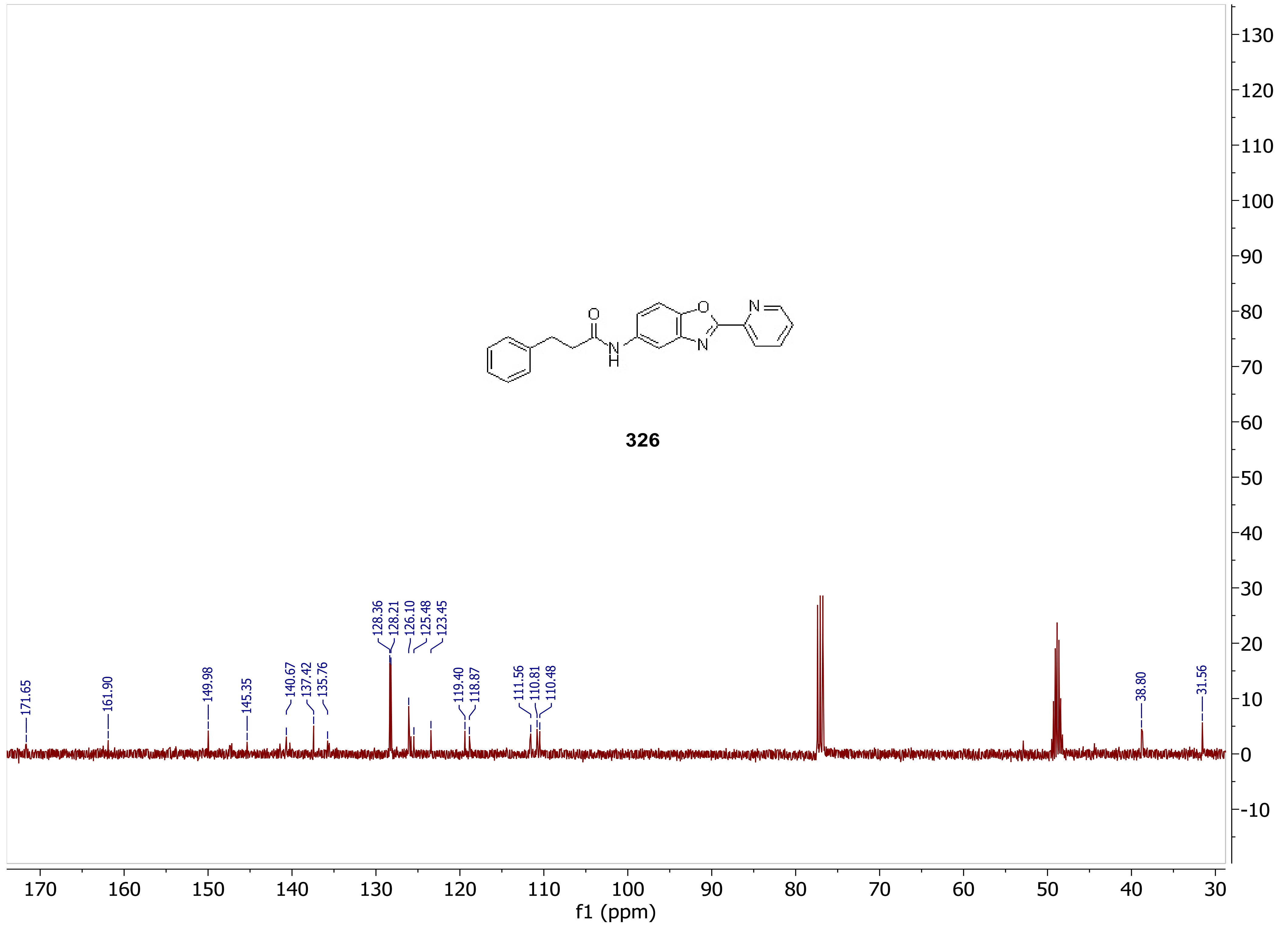

O
N
H
O
N
N
326
171.65
161.90
149.98
145.35
140.67
137.42
135.76
128.36
128.21
126.10
125.48
123.45
119.40
118.87
111.56
110.81
110.48
38.80
31.56
170 160 150 140 130 120 110 100 90 80 70 60 50 40 30
f1 (ppm)
130 120 110 100 90 80 70 60 50 40 30 20 10 0 -10

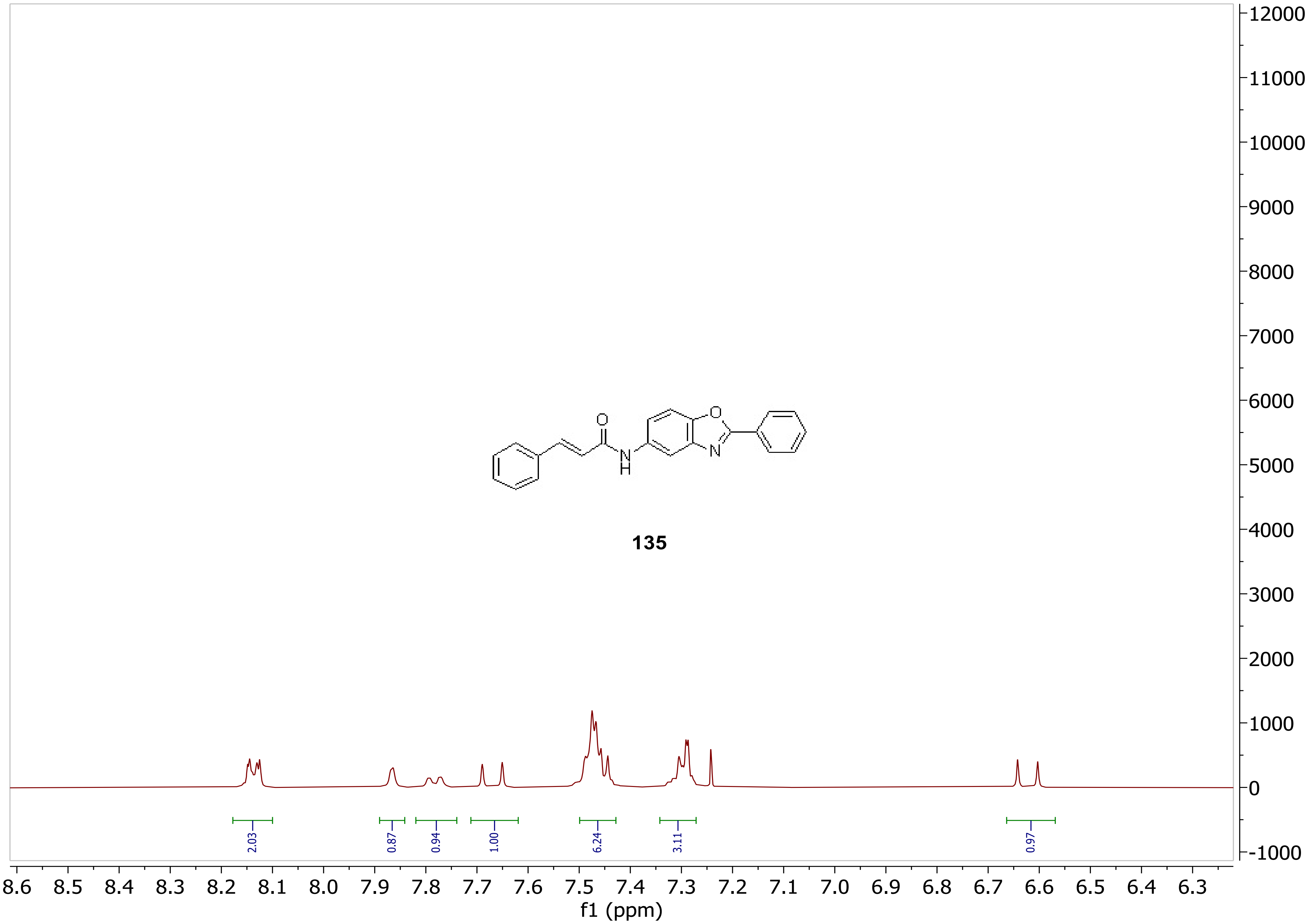

135
2.03
0.87
0.94
1.00
6.24
3.11
0.97
12000
11000
10000
9000
8000
7000
6000
5000
4000
3000
2000
1000
0
-1000
8.6 8.5 8.4 8.3 8.2 8.1 8.0 7.9 7.8 7.7 7.6 7.5 7.4 7.3 7.2 7.1 7.0 6.9 6.8 6.7 6.6 6.5 6.4 6.3
f1 (ppm)
O
N
H

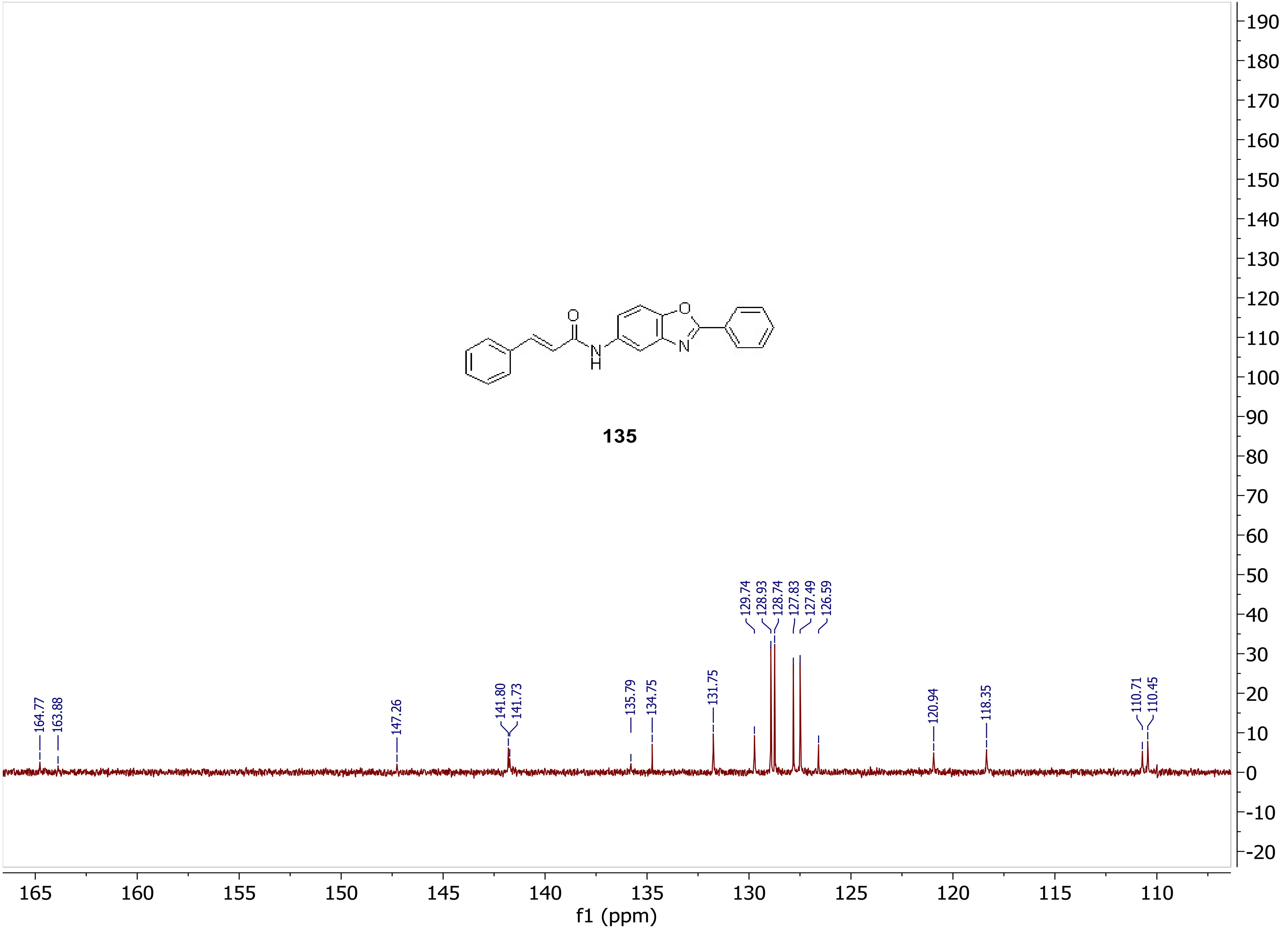

135
164.77
163.88
147.26
141.80
141.73
135.79
134.75
131.75
129.74
128.93
128.74
127.83
127.49
126.59
120.94
118.35
110.71
110.45
f1 (ppm)

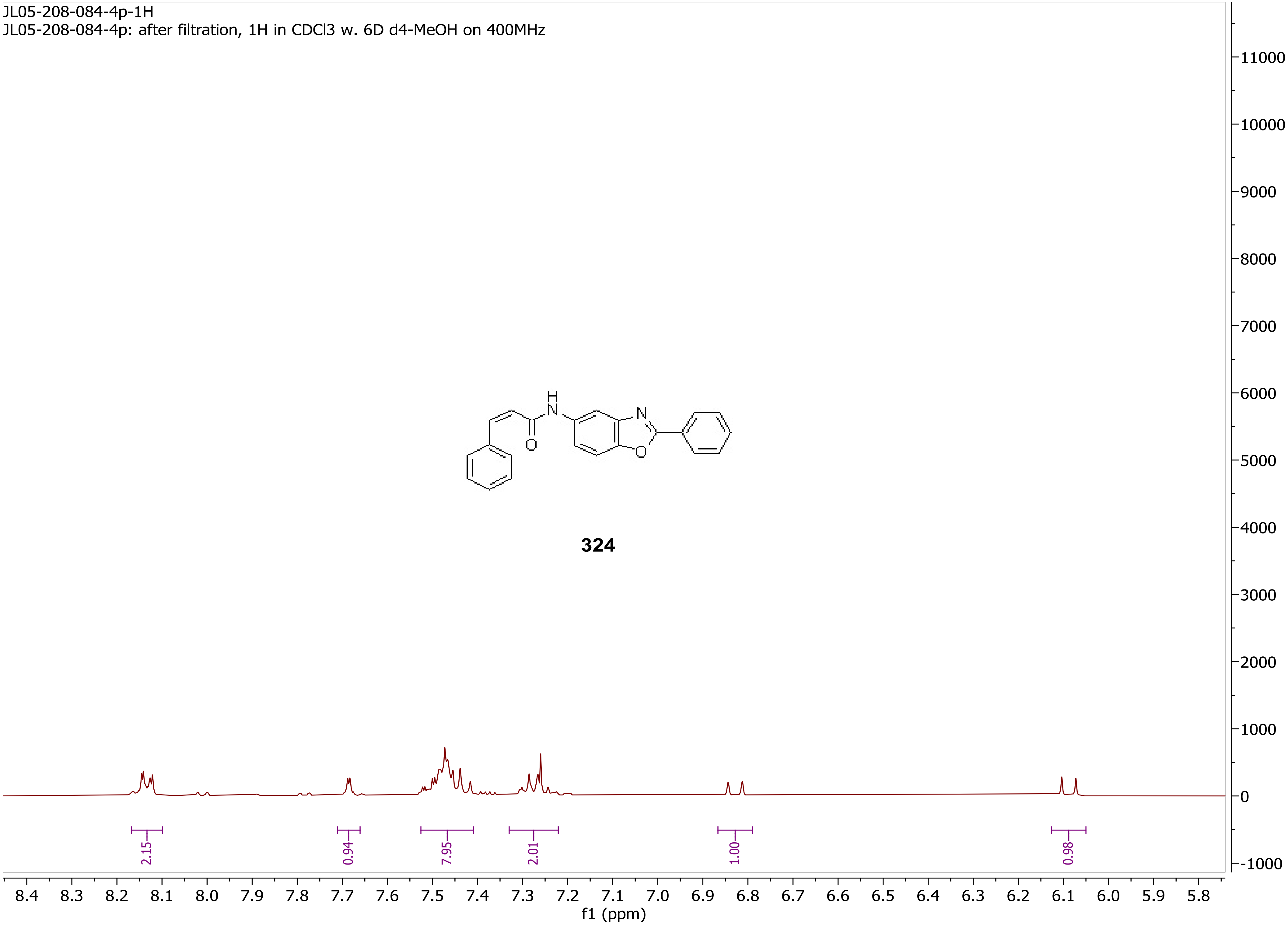
JL05-208-084-4p-1H
JL05-208-084-4p: after filtration, 1H in CDCl3 w. 6D d4-MeOH on 400MHz
324
2.15
0.94
7.95
2.01
1.00
0.98
f1 (ppm)

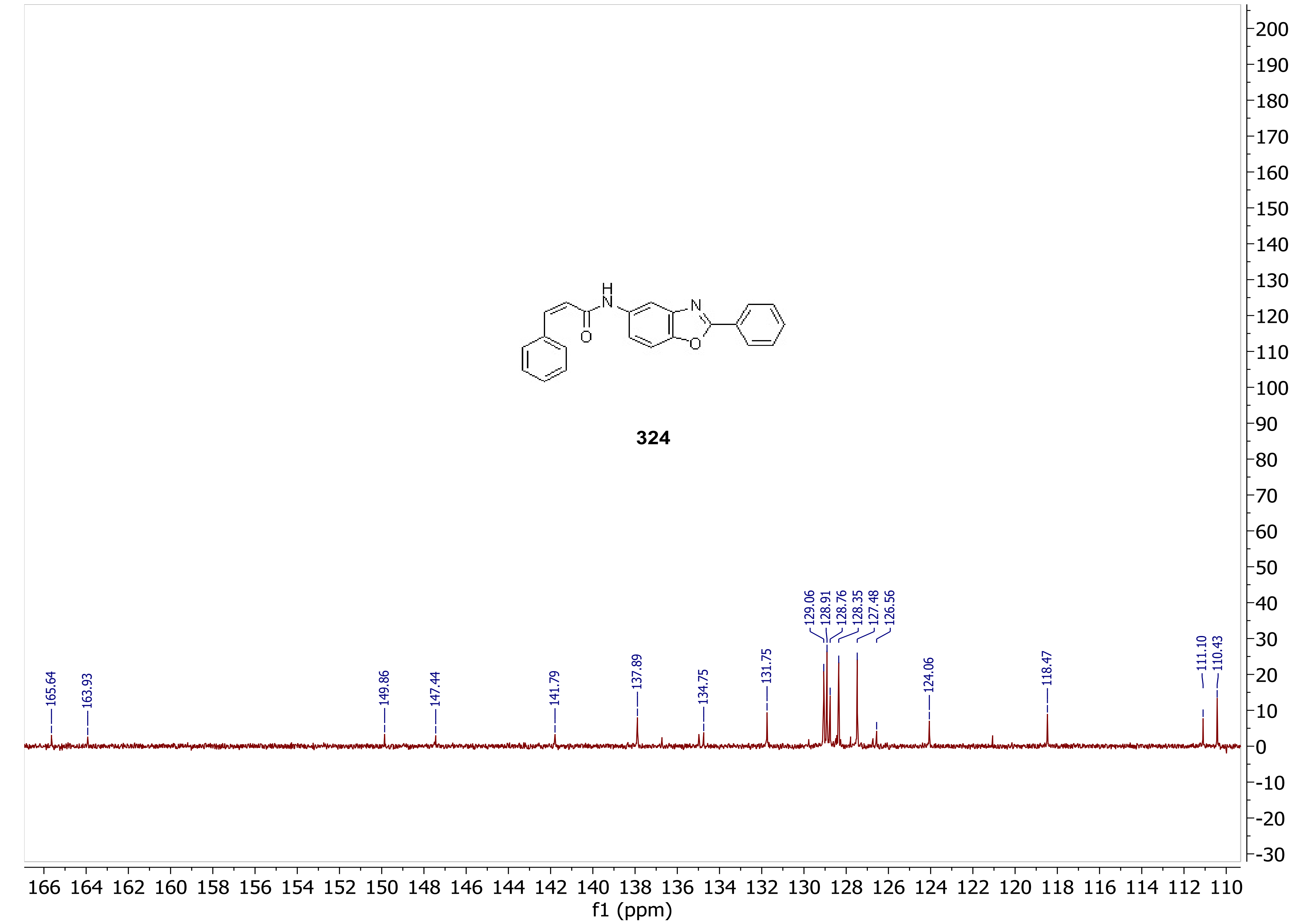
324
165.64
163.93
149.86
147.44
141.79
137.89
134.75
131.75
129.06
128.91
128.76
128.35
127.48
126.56
124.06
118.47
111.10
110.43
f1 (ppm)

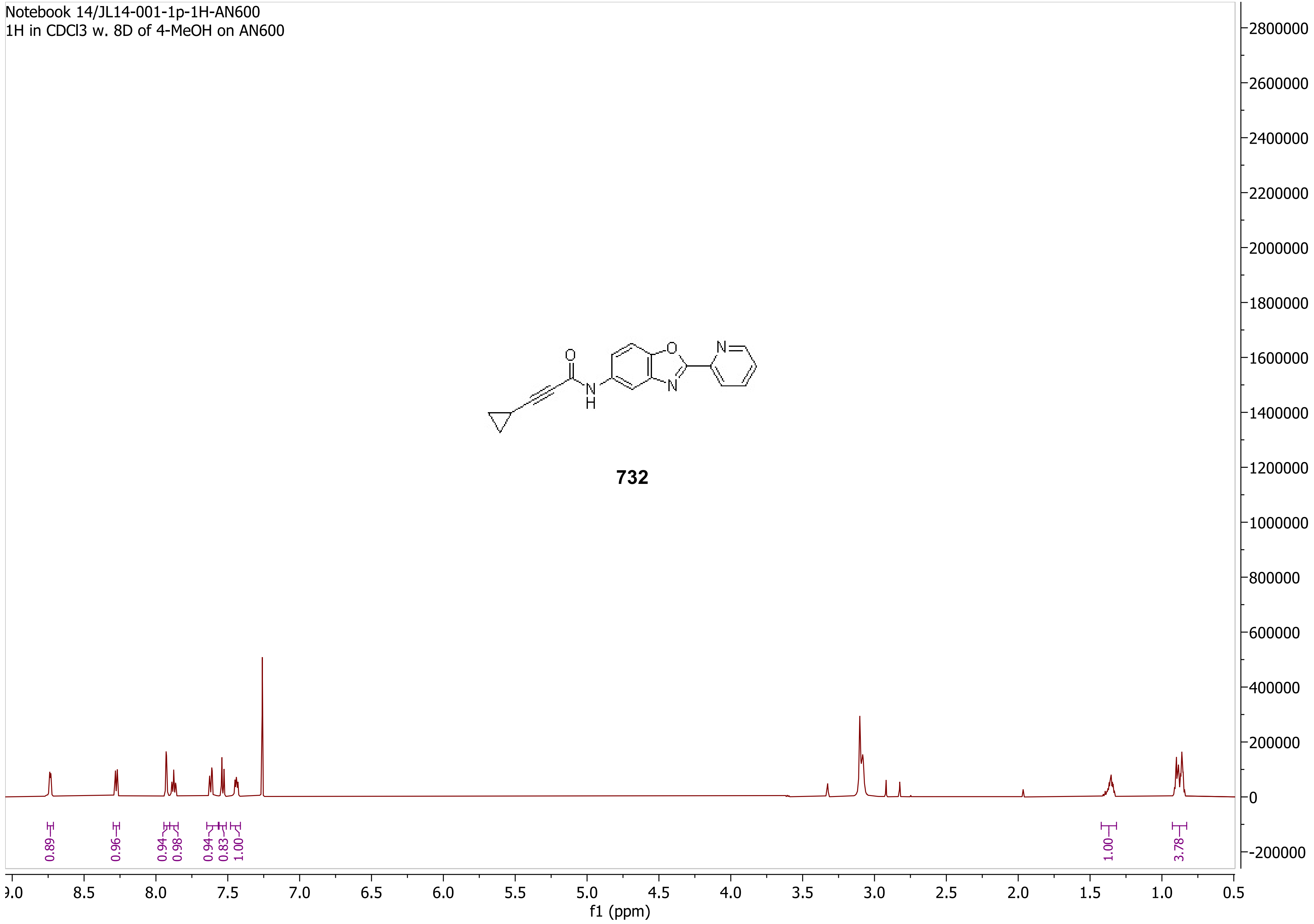
Notebook 14/JL14-001-1p-1H-AN600
1H in CDCl3 w. 8D of 4-MeOH on AN600
O
N
H
O
N
N
732
0.89
0.96
0.94
0.98
0.94
0.83
1.00
1.00
3.78
2800000
2600000
2400000
2200000
2000000
1800000
1600000
1400000
1200000
1000000
800000
600000
400000
200000
0
-200000
9.0
8.5
8.0
7.5
7.0
6.5
6.0
5.5
5.0
4.5
4.0
3.5
3.0
2.5
2.0
1.5
1.0
0.5
f1 (ppm)

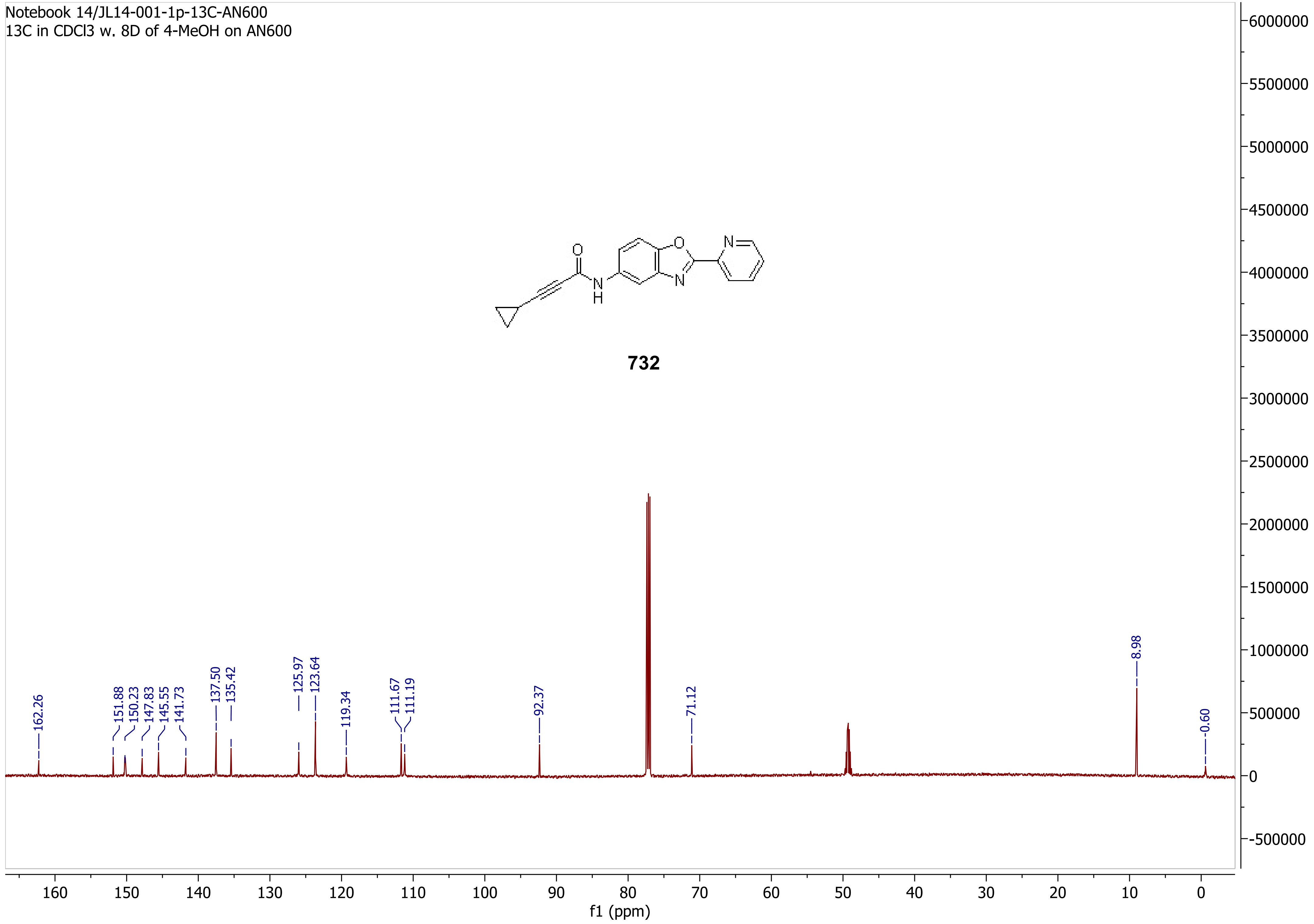
Notebook 14/JL14-001-1p-13C-AN600
13C in CDCl3 w. 8D of 4-MeOH on AN600
O
N
H
O
N
N
732
162.26
151.88
150.23
147.83
145.55
141.73
137.50
135.42
125.97
123.64
119.34
111.67
111.19
92.37
71.12
8.98
-0.60
6000000
5500000
5000000
4500000
4000000
3500000
3000000
2500000
2000000
1500000
1000000
500000
0
-500000
160
150
140
130
120
110
100
90
80
70
60
50
40
30
20
10
0
f1 (ppm)

Notebook 14/JL14-001-2p-1H-AN600
off-white solid after water wash, 1H in CDCl3 w. 8D of d4-MeOH on AN600

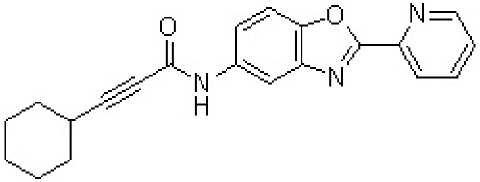


**733**

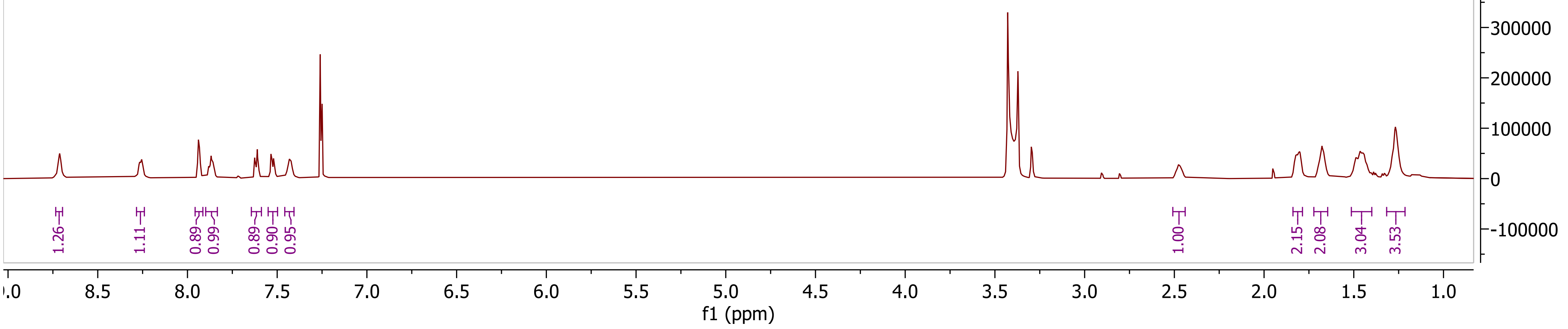

Notebook 14/JL14-001-2p-13C-AN600
off-white solid after water wash, 13C in CDCl3 w. 8D of d4-MeOH on AN600

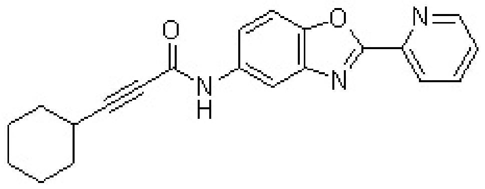

**733**

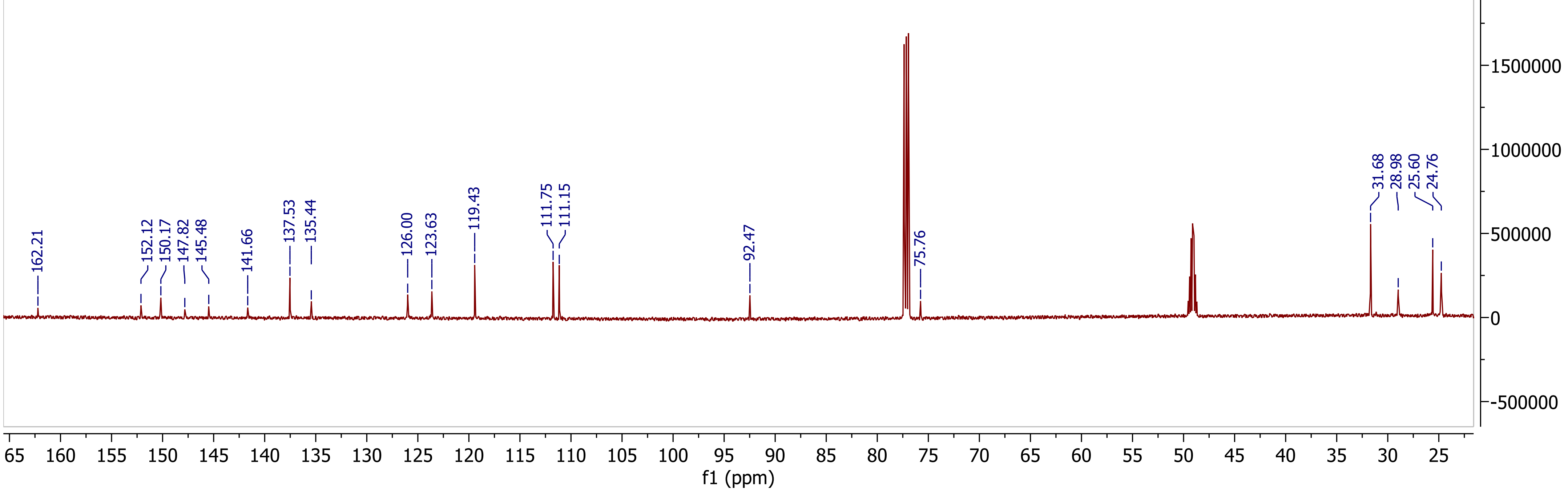

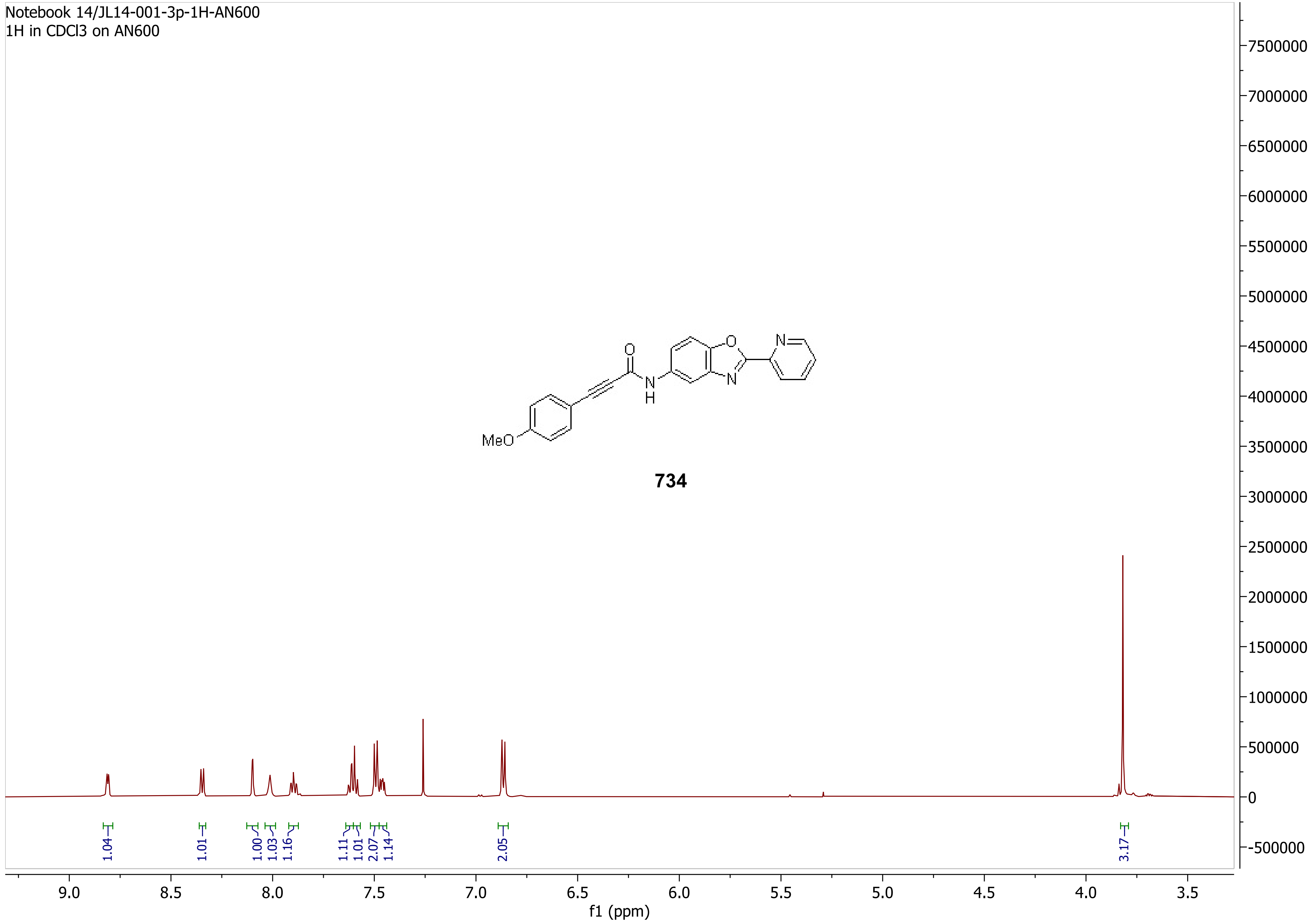
Notebook 14/JL14-001-3p-1H-AN600
1H in CDCl3 on AN600
MeO
O
N
H
N
734
1.04
1.01
1.00
1.03
1.16
1.11
1.01
2.07
1.14
2.05
3.17
9.0
8.5
8.0
7.5
7.0
6.5
6.0
5.5
5.0
4.5
4.0
3.5
f1 (ppm)
7500000
7000000
6500000
6000000
5500000
5000000
4500000
4000000
3500000
3000000
2500000
2000000
1500000
1000000
500000
0
-500000

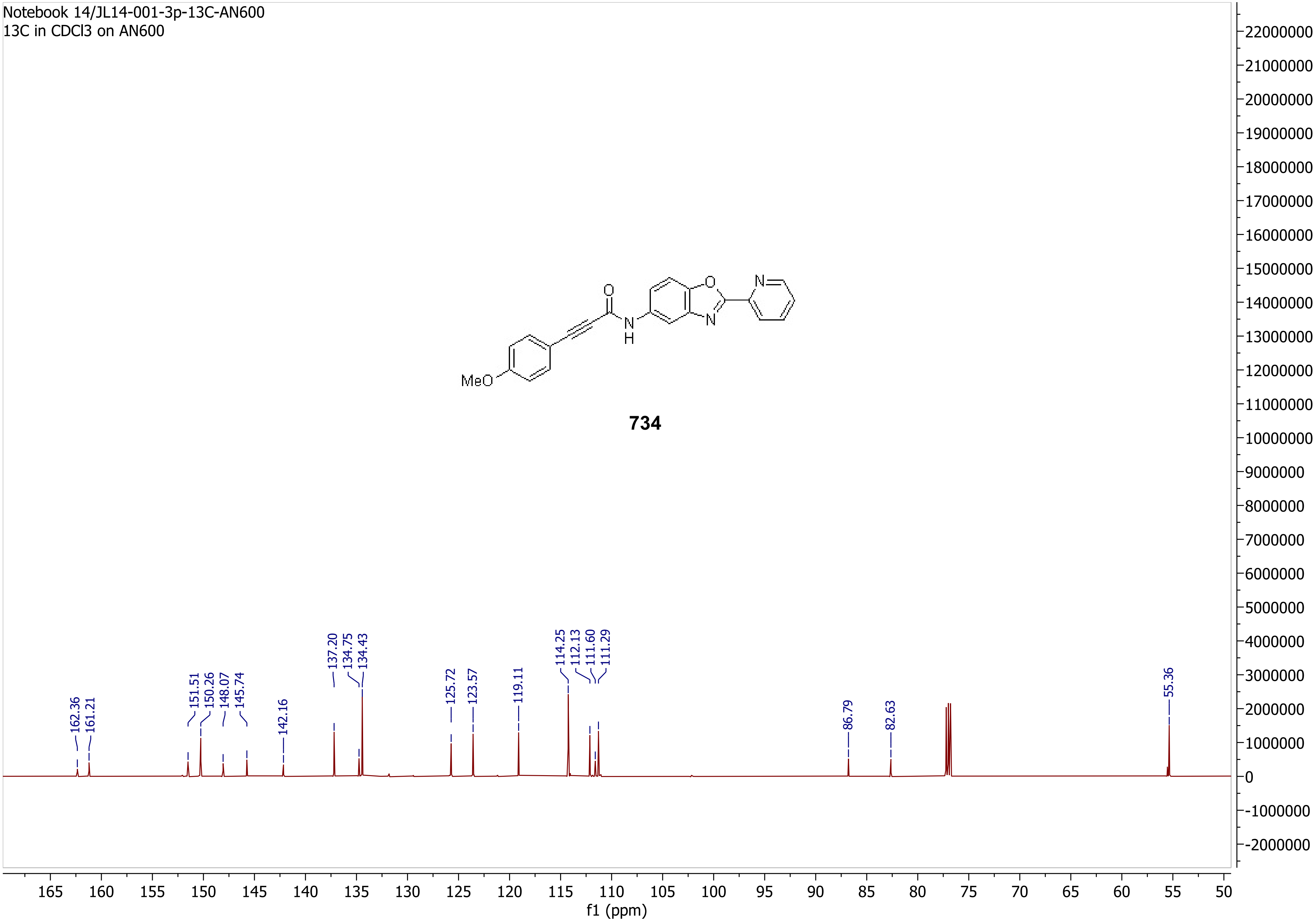
Notebook 14/JL14-001-3p-13C-AN600
13C in CDCl3 on AN600
MeO
O
N
H
N
O
N
734
162.36
161.21
151.51
150.26
148.07
145.74
142.16
137.20
134.75
134.43
125.72
123.57
119.11
114.25
112.13
111.60
111.29
86.79
82.63
55.36
f1 (ppm)

Notebook 14/JL14-002-1p-1H-AN600
off-white solid after 1) plug; 2) trituration, 1H in CDCl3 on AN600

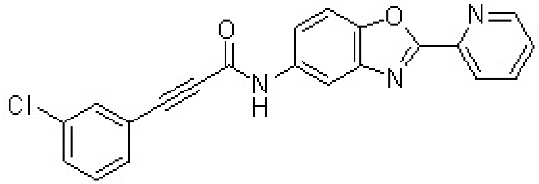


**735**

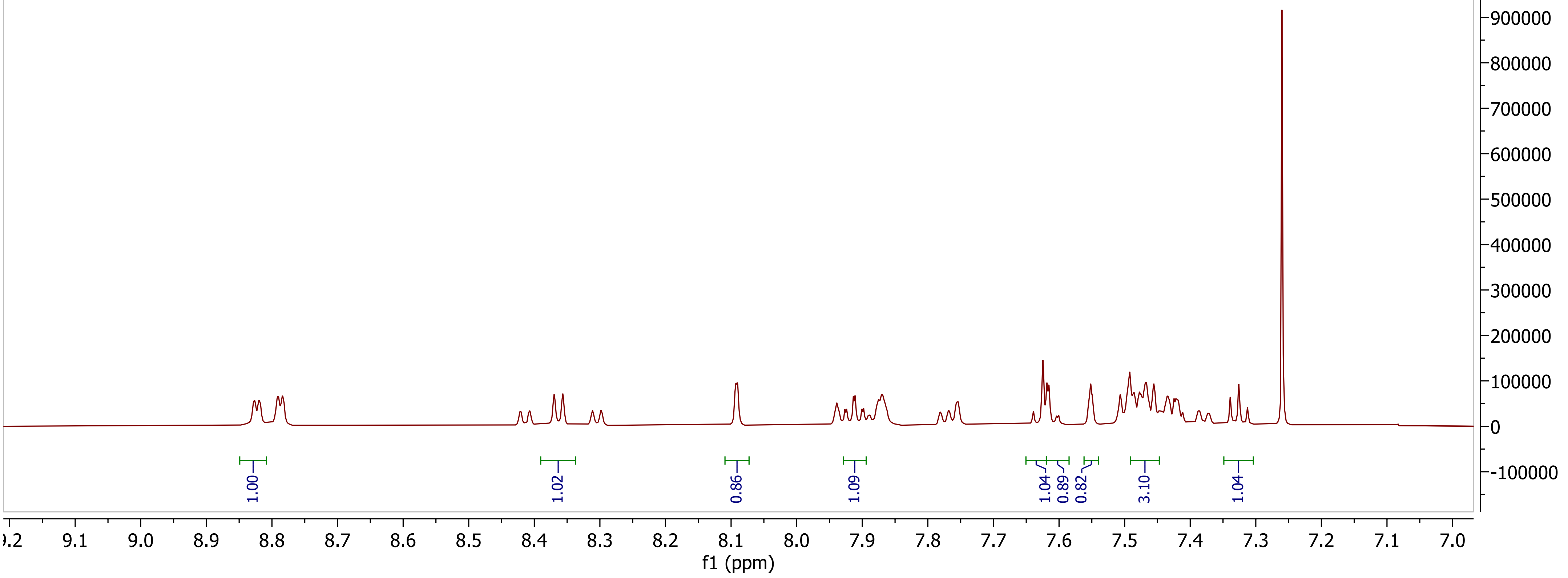

Notebook 14/JL14-002-1p-13C-AN600
off-white solid after 1) plug; 2) trituration, 13C in CDCl3 on AN600

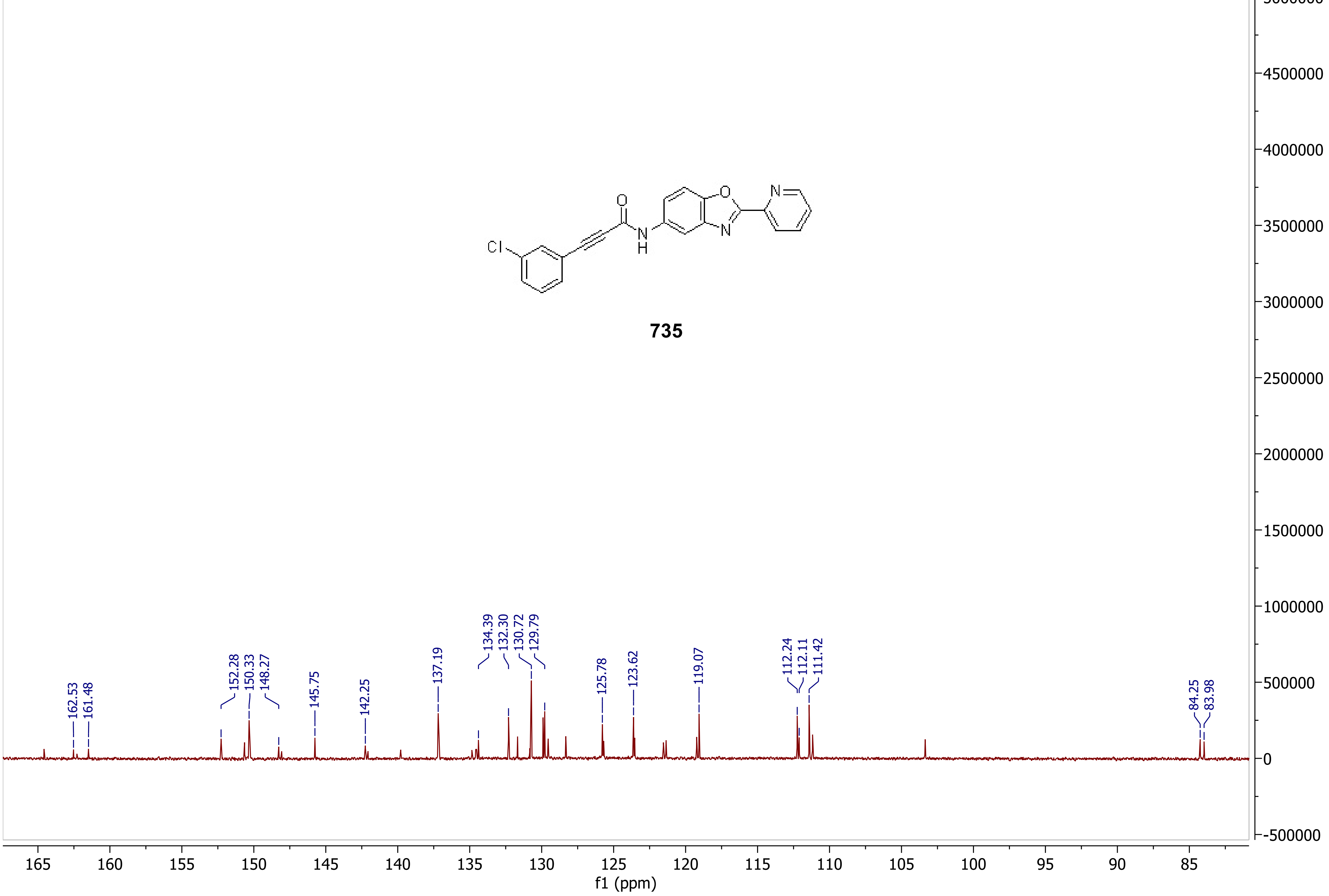


**735**

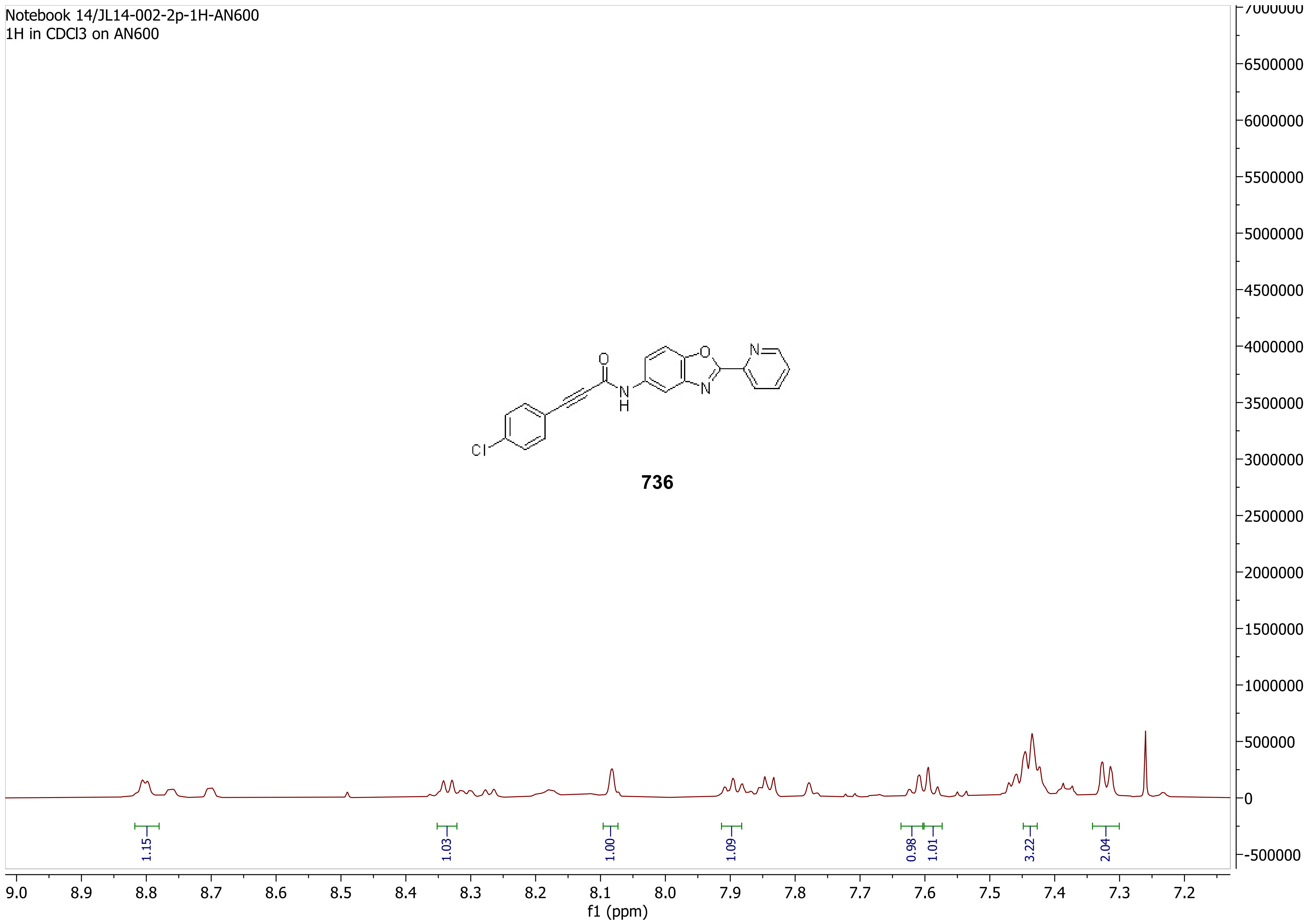
Notebook 14/JL14-002-2p-1H-AN600
1H in CDCl3 on AN600
736
1.15
1.03
1.00
1.09
0.98
1.01
3.22
2.04
9.0 8.9 8.8 8.7 8.6 8.5 8.4 8.3 8.2 8.1 8.0 7.9 7.8 7.7 7.6 7.5 7.4 7.3 7.2
f1 (ppm)
7000000 6500000 6000000 5500000 5000000 4500000 4000000 3500000 3000000 2500000 2000000 1500000 1000000 500000 0 -500000

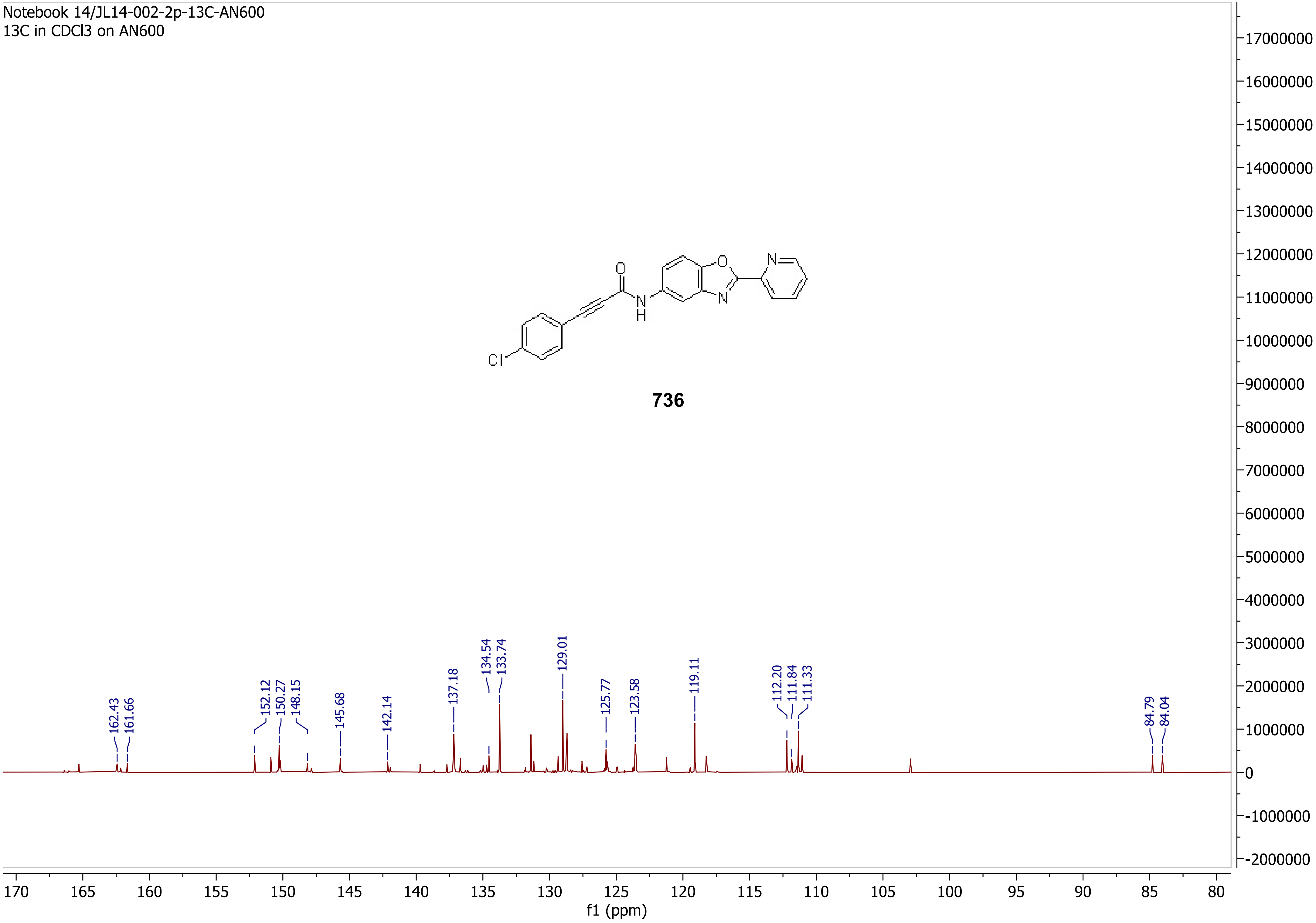
Notebook 14/JL14-002-2p-13C-AN600
13C in CDCl3 on AN600
O
N
H
O
N
N
Cl
736
162.43
161.66
152.12
150.27
148.15
145.68
142.14
137.18
134.54
133.74
129.01
125.77
123.58
119.11
112.20
111.84
111.33
84.79
84.04
17000000
16000000
15000000
14000000
13000000
12000000
11000000
10000000
9000000
8000000
7000000
6000000
5000000
4000000
3000000
2000000
1000000
0
-1000000
-2000000
170
165
160
155
150
145
140
135
130
125
120
115
110
105
100
95
90
85
80
f1 (ppm)

Notebook 14/JL14-004-1p-1H-AN600-1
1H in CDCl3 on AN600

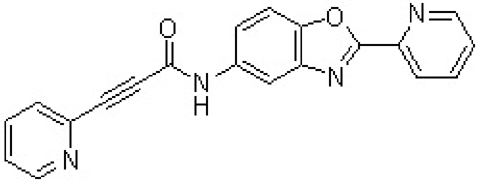

**737**

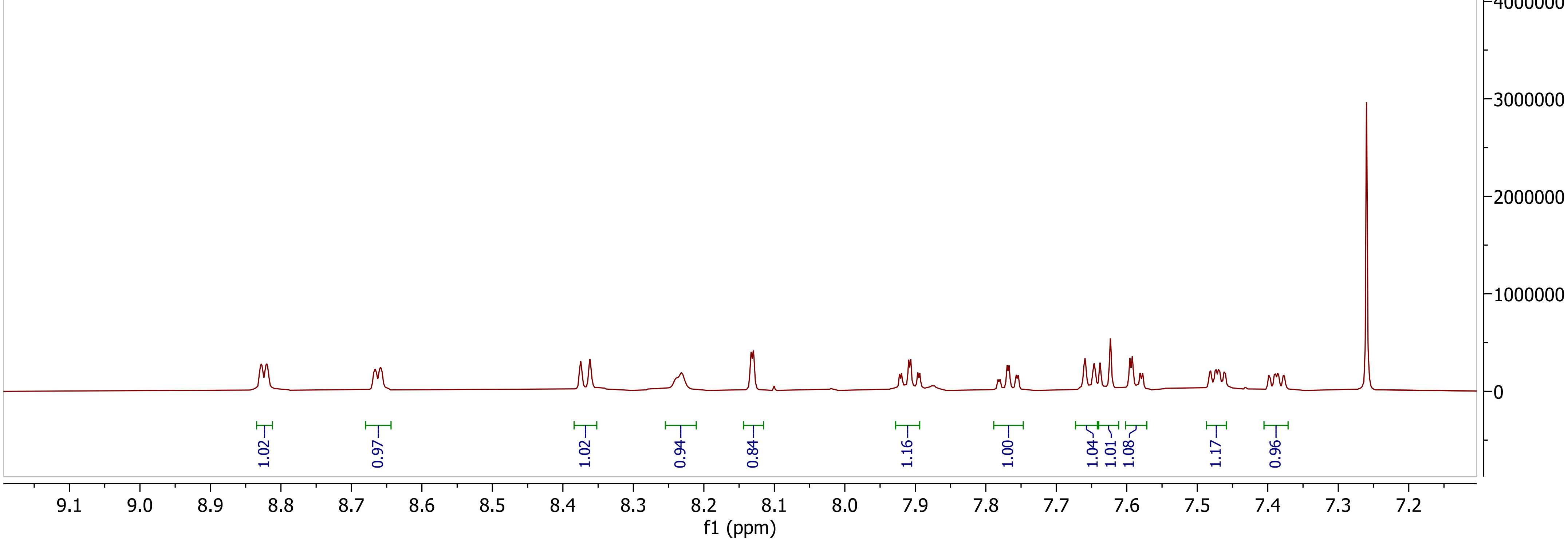

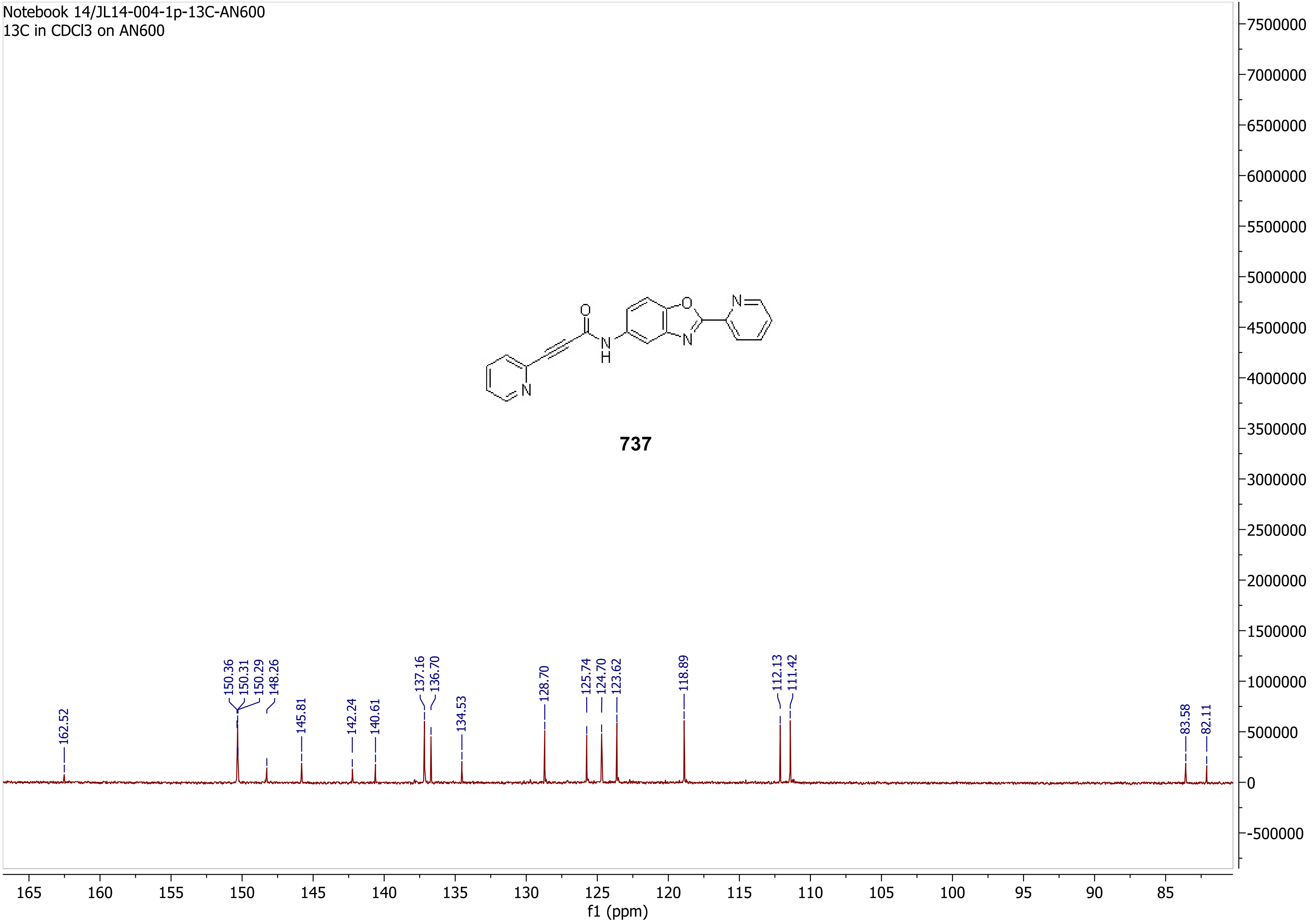
Notebook 14/JL14-004-1p-13C-AN600
13C in CDCl3 on AN600
737
162.52
150.36
150.31
150.29
148.26
145.81
142.24
140.61
137.16
136.70
134.53
128.70
125.74
124.70
123.62
118.89
112.13
111.42
83.58
82.11
165 160 155 150 145 140 135 130 125 120 115 110 105 100 95 90 85
f1 (ppm)
7500000
7000000
6500000
6000000
5500000
5000000
4500000
4000000
3500000
3000000
2500000
2000000
1500000
1000000
500000
0
-500000

Notebook 14/JL14-004-2p-1H-AN600
white ppt, poor solubility, 1H in d4-MeOH w. 12D CDCl3 w. heat, on AN600

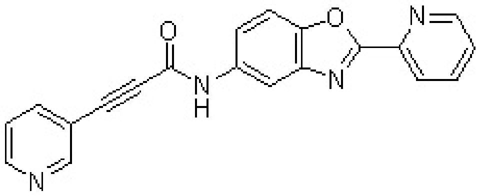

**738**

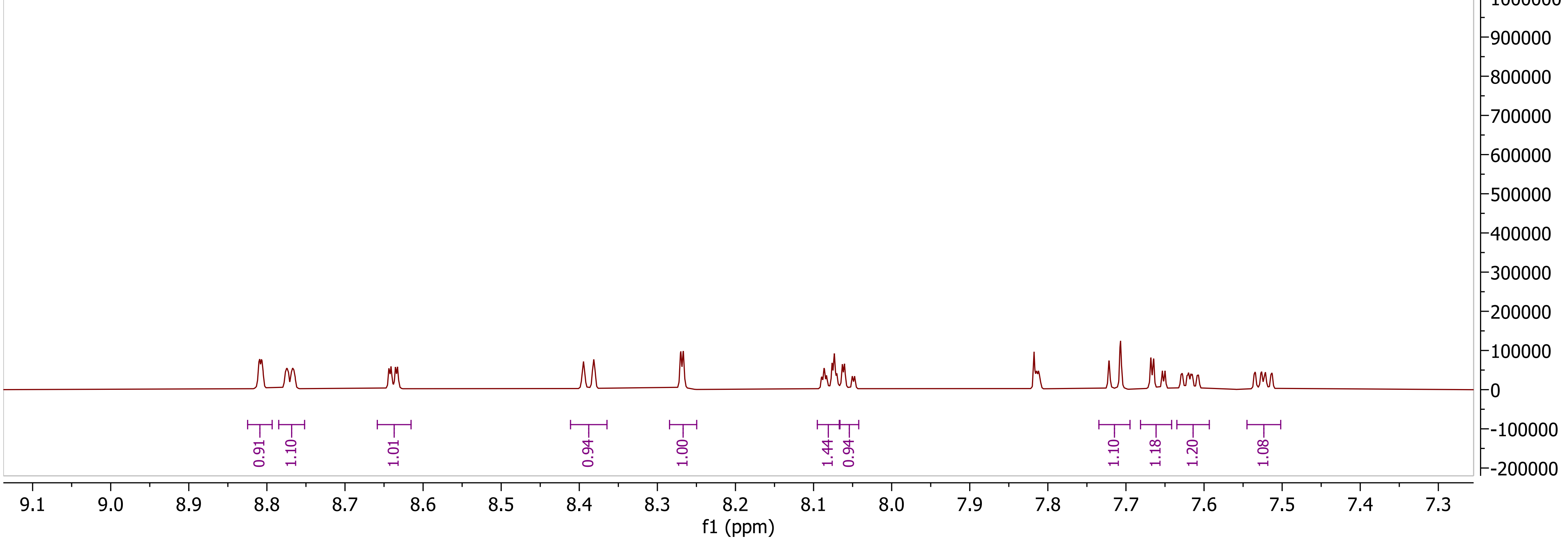

Notebook 14/JL14-004-2p-13C-AN600
white ppt, poor solubility, 13C in d4-MeOH w. 12D CDCl3 w. heat, on AN600

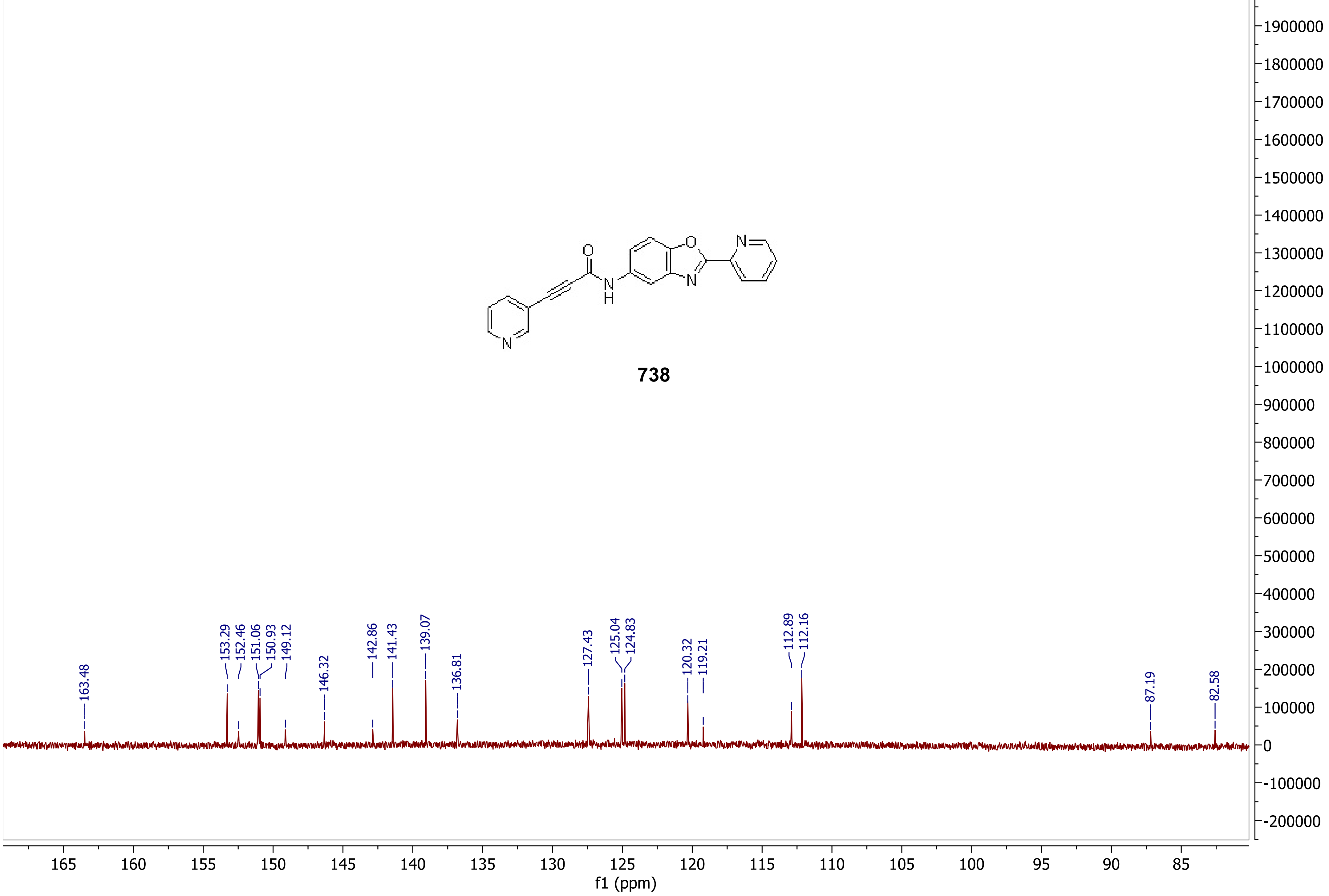

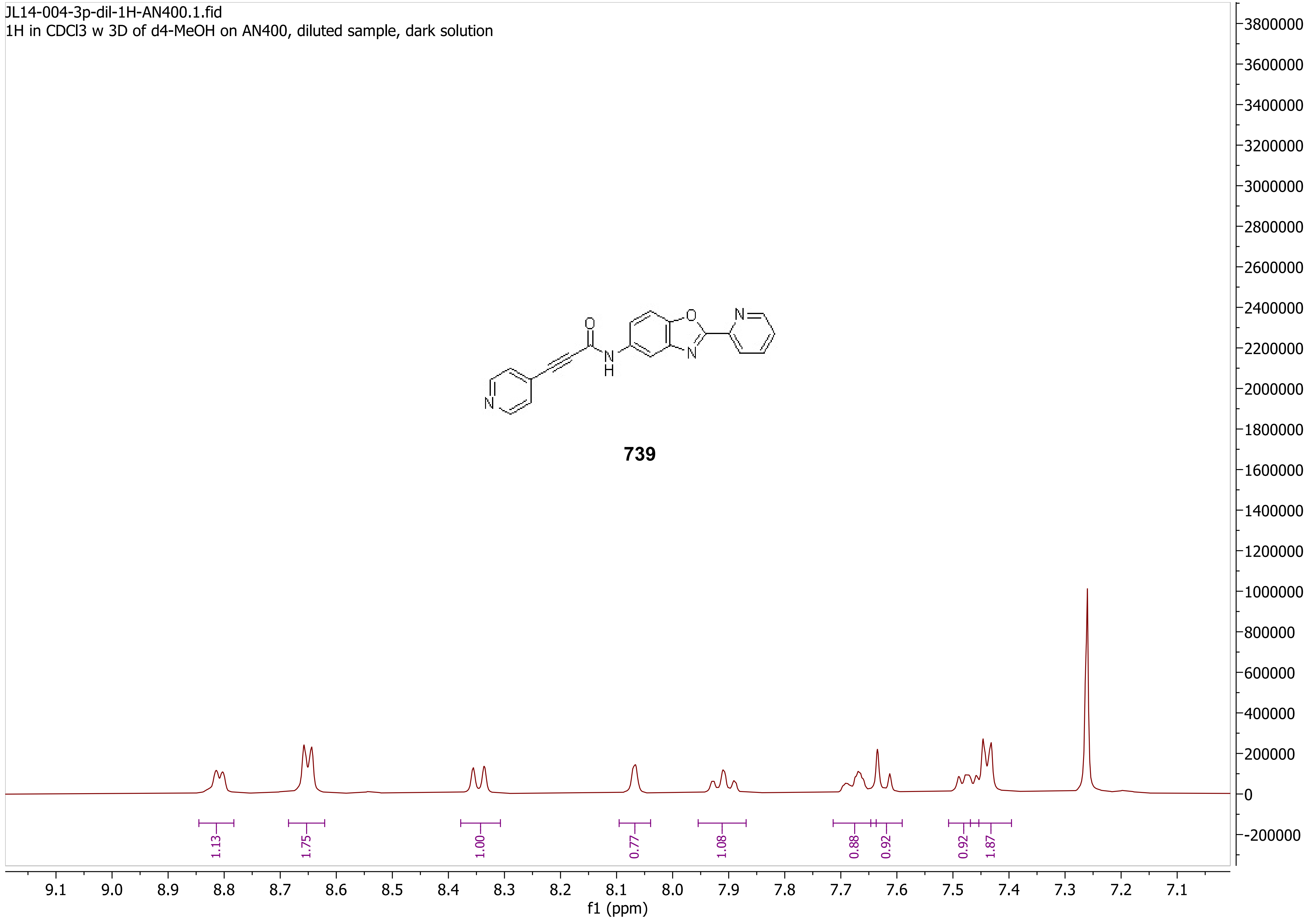
JL14-004-3p-dil-1H-AN400.1.fid
1H in CDCl3 w 3D of d4-MeOH on AN400, diluted sample, dark solution
739
1.13
1.75
1.00
0.77
1.08
0.88
0.92
0.92
1.87
f1 (ppm)

Notebook 14/JL14-004-3p-b-13C-AN600
light-brown solution quickly turned to greenish, 13C in CDCl3 w 8D d4-MeOH on AN600

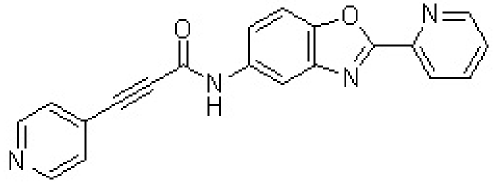

**739**

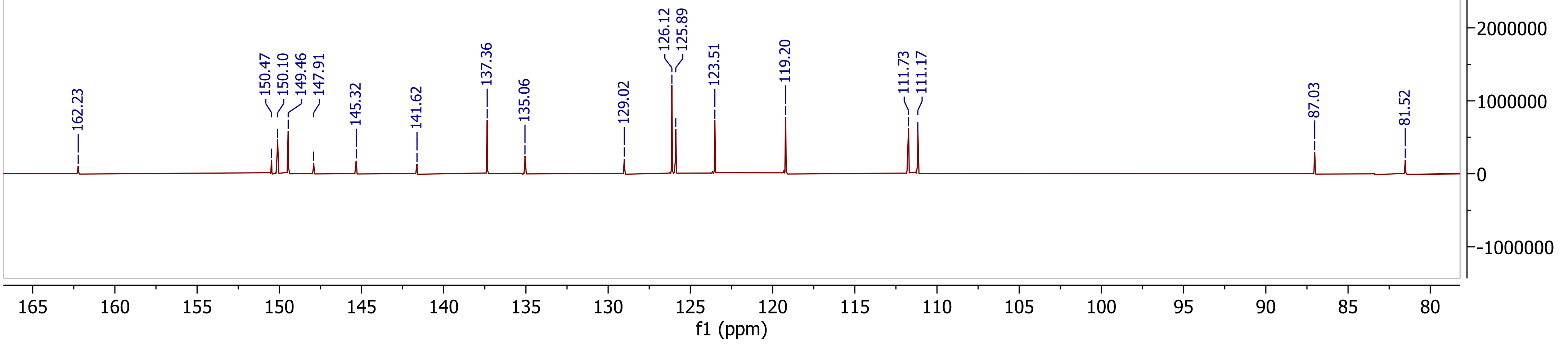

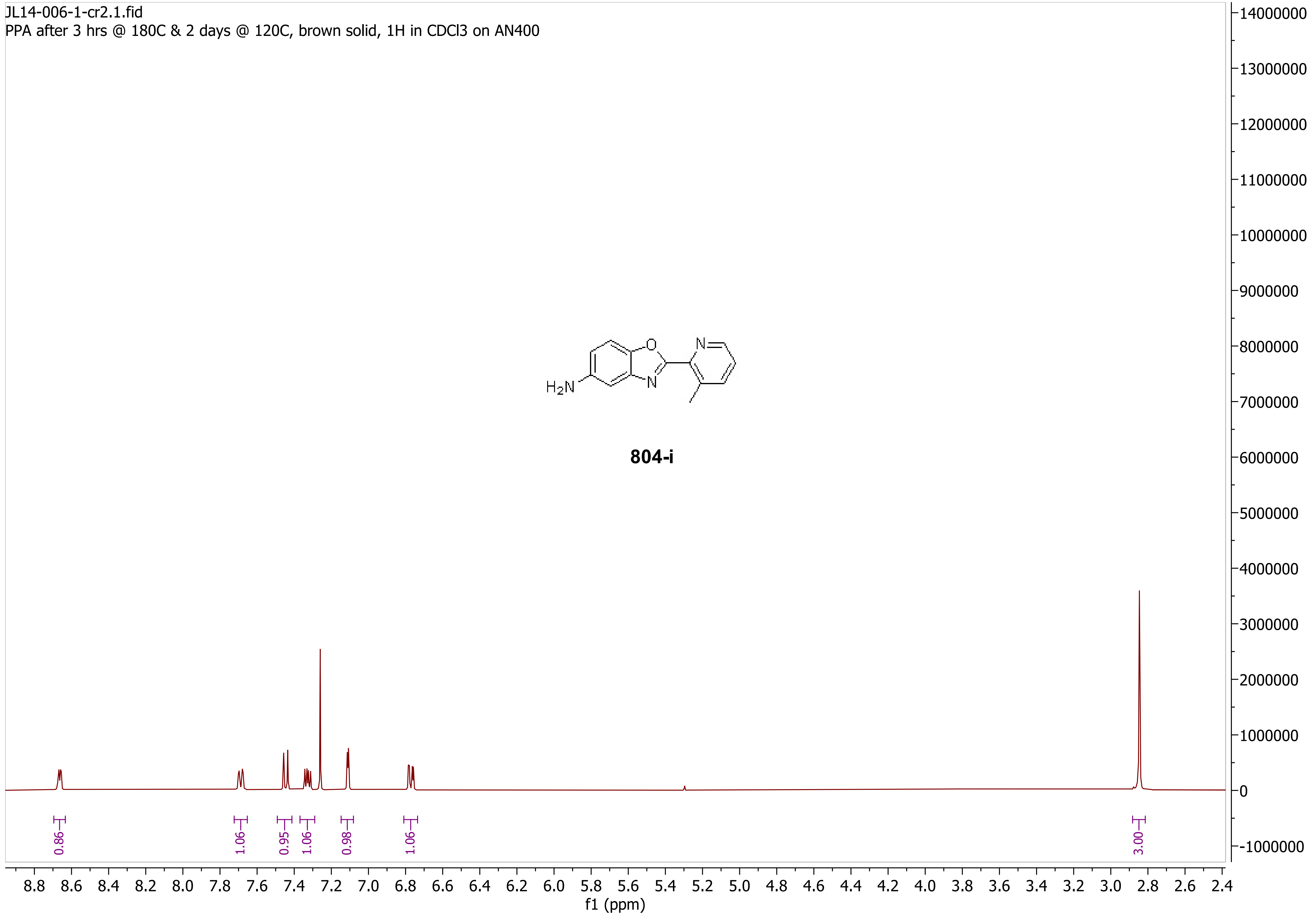
JL14-006-1-cr2.1.fid
PPA after 3 hrs @ 180C & 2 days @ 120C, brown solid, 1H in CDCl3 on AN400
H2N
O
N
N
804-i
0.86
1.06
0.95
1.06
0.98
1.06
3.00
f1 (ppm)
8.8 8.6 8.4 8.2 8.0 7.8 7.6 7.4 7.2 7.0 6.8 6.6 6.4 6.2 6.0 5.8 5.6 5.4 5.2 5.0 4.8 4.6 4.4 4.2 4.0 3.8 3.6 3.4 3.2 3.0 2.8 2.6 2.4
14000000
13000000
12000000
11000000
10000000
9000000
8000000
7000000
6000000
5000000
4000000
3000000
2000000
1000000
0
-1000000

Notebook 14/JL14-006-1p-13C-AN600
conc. brown solution, 13C in CDCl3 on AN600

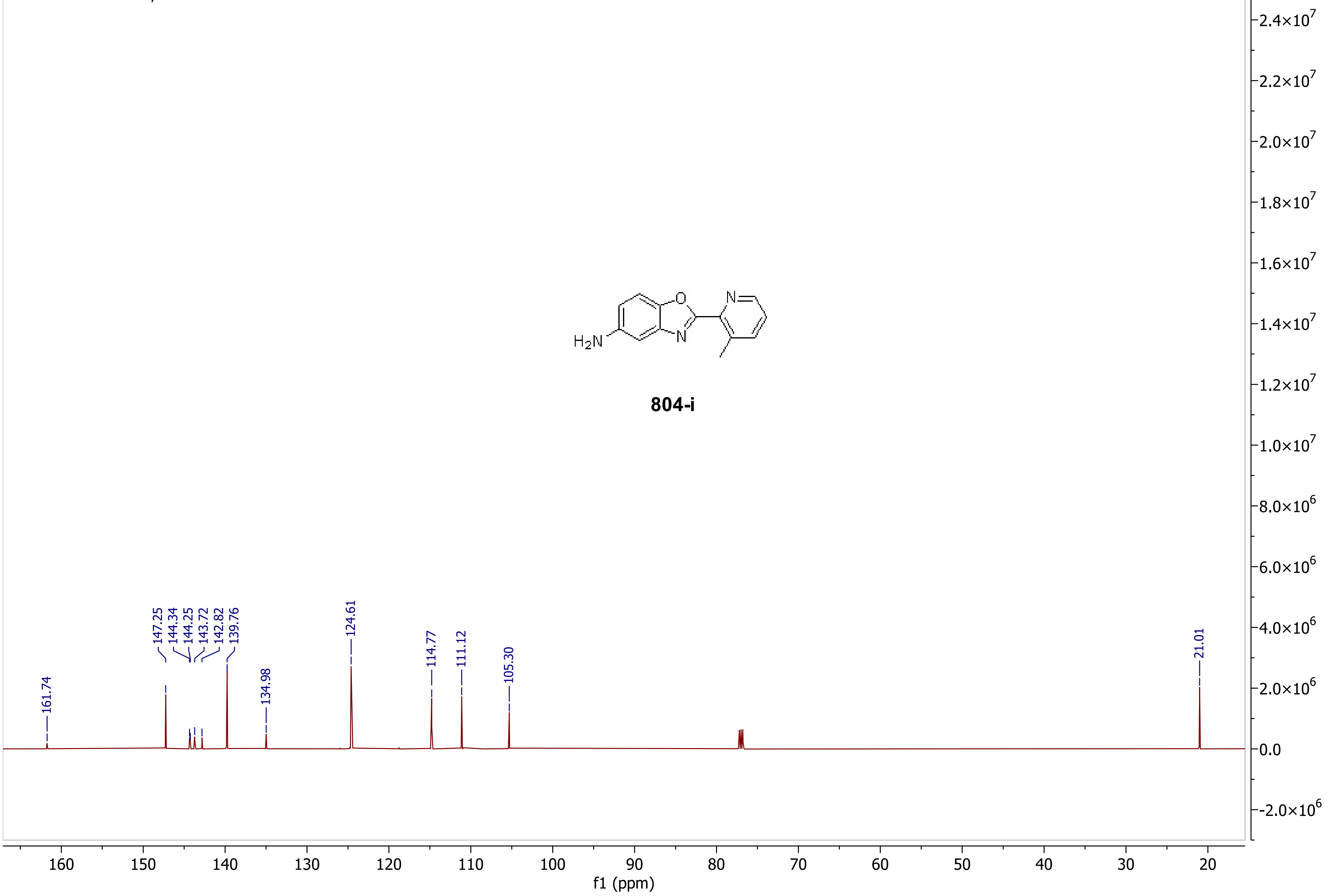

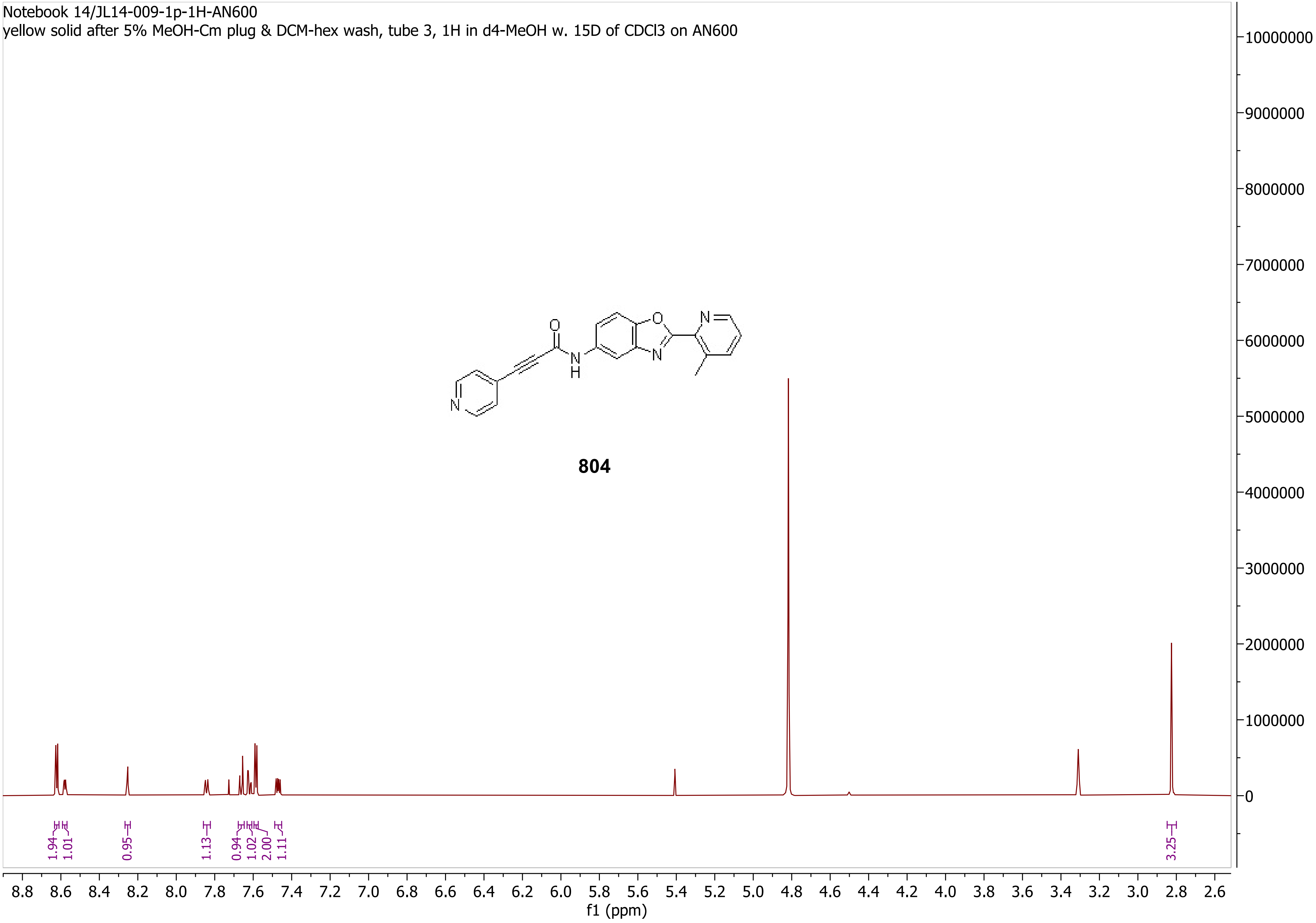
Notebook 14/JL14-009-1p-1H-AN600
yellow solid after 5% MeOH-Cm plug & DCM-hex wash, tube 3, 1H in d4-MeOH w. 15D of CDCl3 on AN600
804
1.94
1.01
0.95
1.13
0.94
1.02
2.00
1.11
3.25
8.8 8.6 8.4 8.2 8.0 7.8 7.6 7.4 7.2 7.0 6.8 6.6 6.4 6.2 6.0 5.8 5.6 5.4 5.2 5.0 4.8 4.6 4.4 4.2 4.0 3.8 3.6 3.4 3.2 3.0 2.8 2.6
f1 (ppm)
10000000
9000000
8000000
7000000
6000000
5000000
4000000
3000000
2000000
1000000
0

Notebook 14/JL14-009-1p-13C-AN600
yellow solid after 5% MeOH-Cm plug & DCM-hex wash, tube 3, 13C in d4-MeOH w. 15D of CDCl3 on AN600

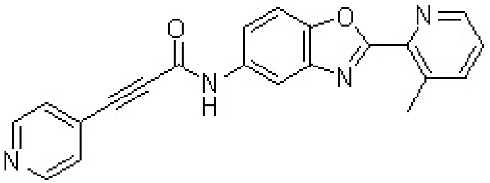

**804**

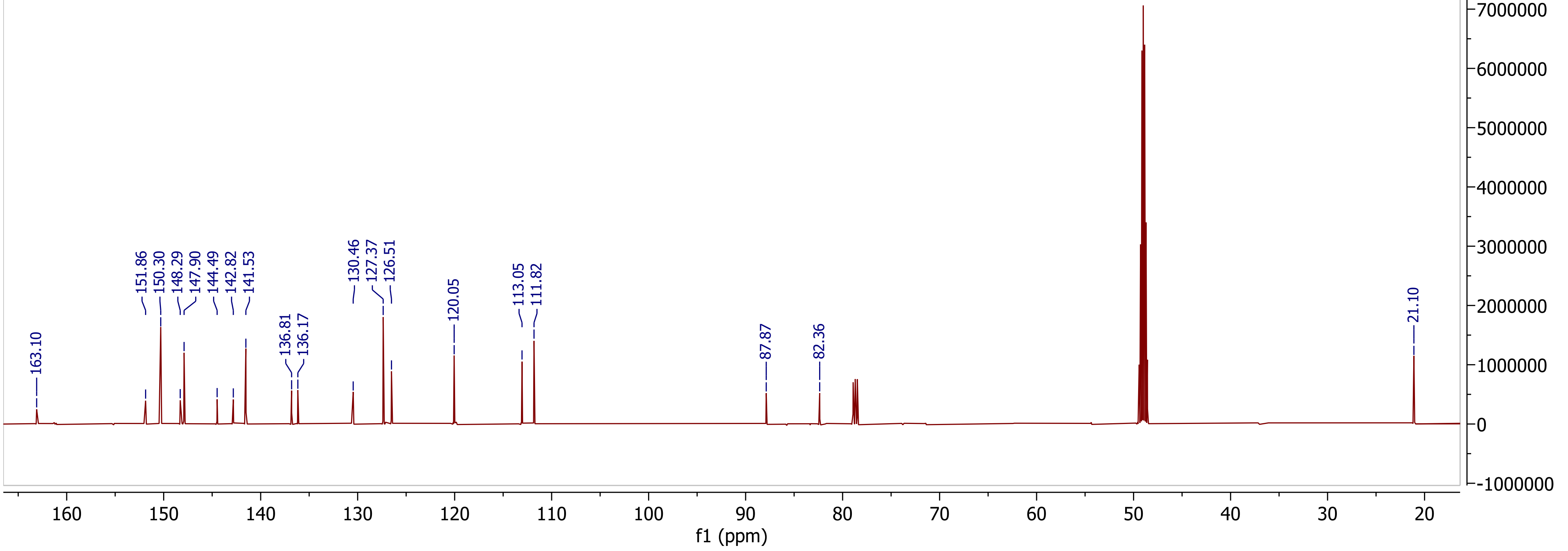

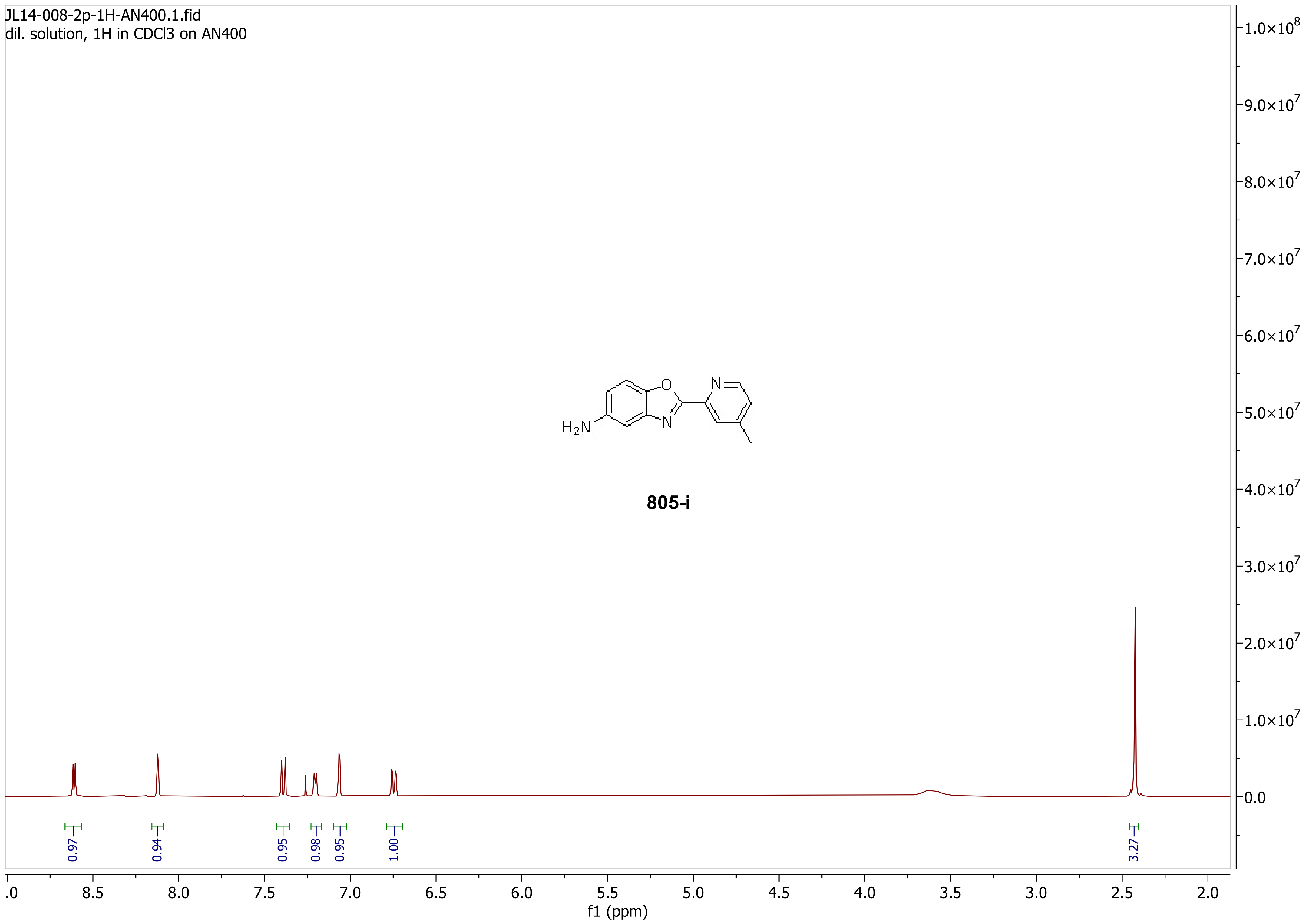
JL14-008-2p-1H-AN400.1.fid
dil. solution, 1H in CDCl3 on AN400
H2N
O
N
N
805-i
0.97
0.94
0.95
0.98
0.95
1.00
3.27
f1 (ppm)
1.0×10^8
9.0×10^7
8.0×10^7
7.0×10^7
6.0×10^7
5.0×10^7
4.0×10^7
3.0×10^7
2.0×10^7
1.0×10^7
0.0

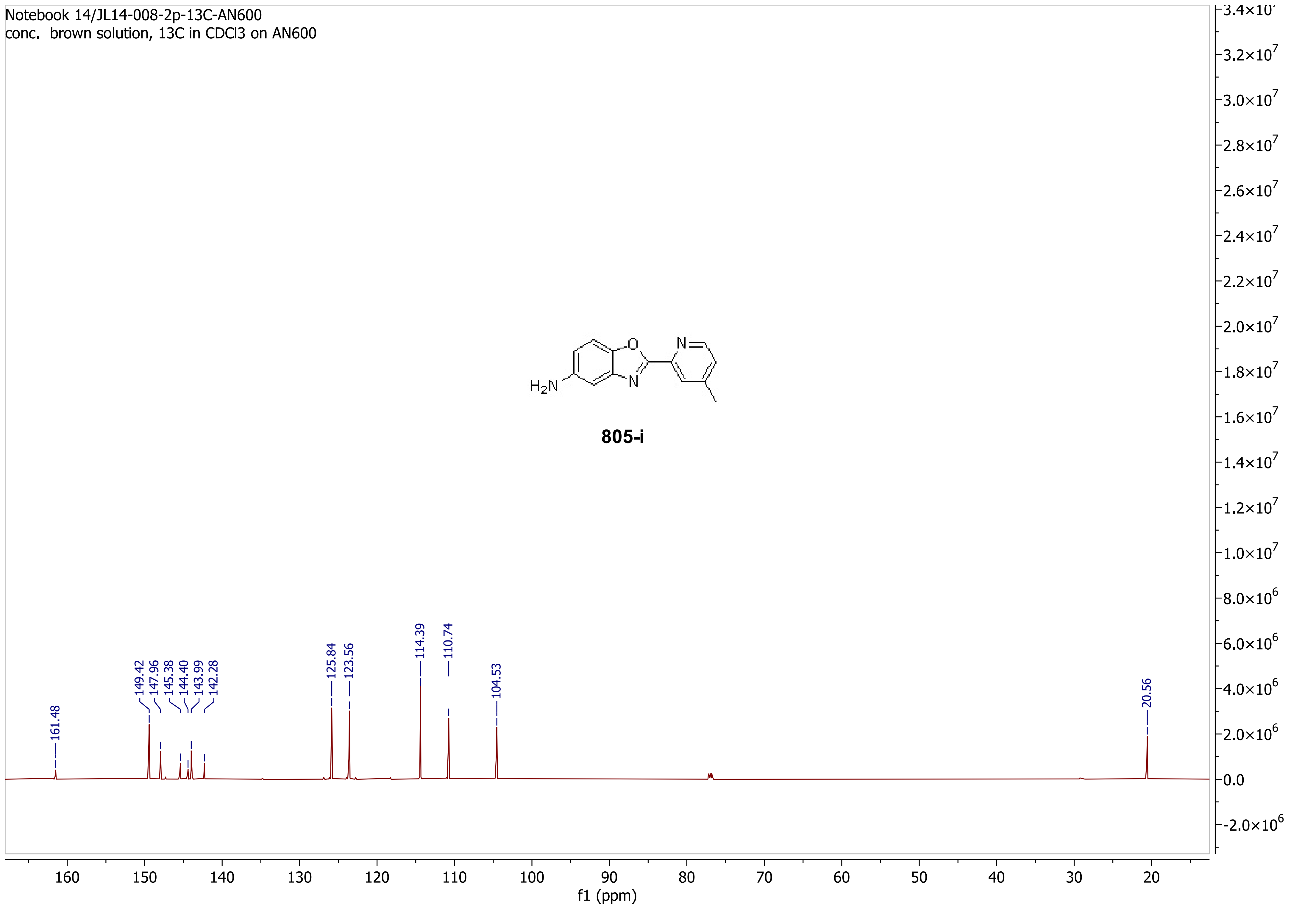
Notebook 14/JL14-008-2p-13C-AN600
conc. brown solution, 13C in CDCl3 on AN600
H2N
O
N
N
805-i
161.48
149.42
147.96
145.38
144.40
143.99
142.28
125.84
123.56
114.39
110.74
104.53
20.56
f1 (ppm)

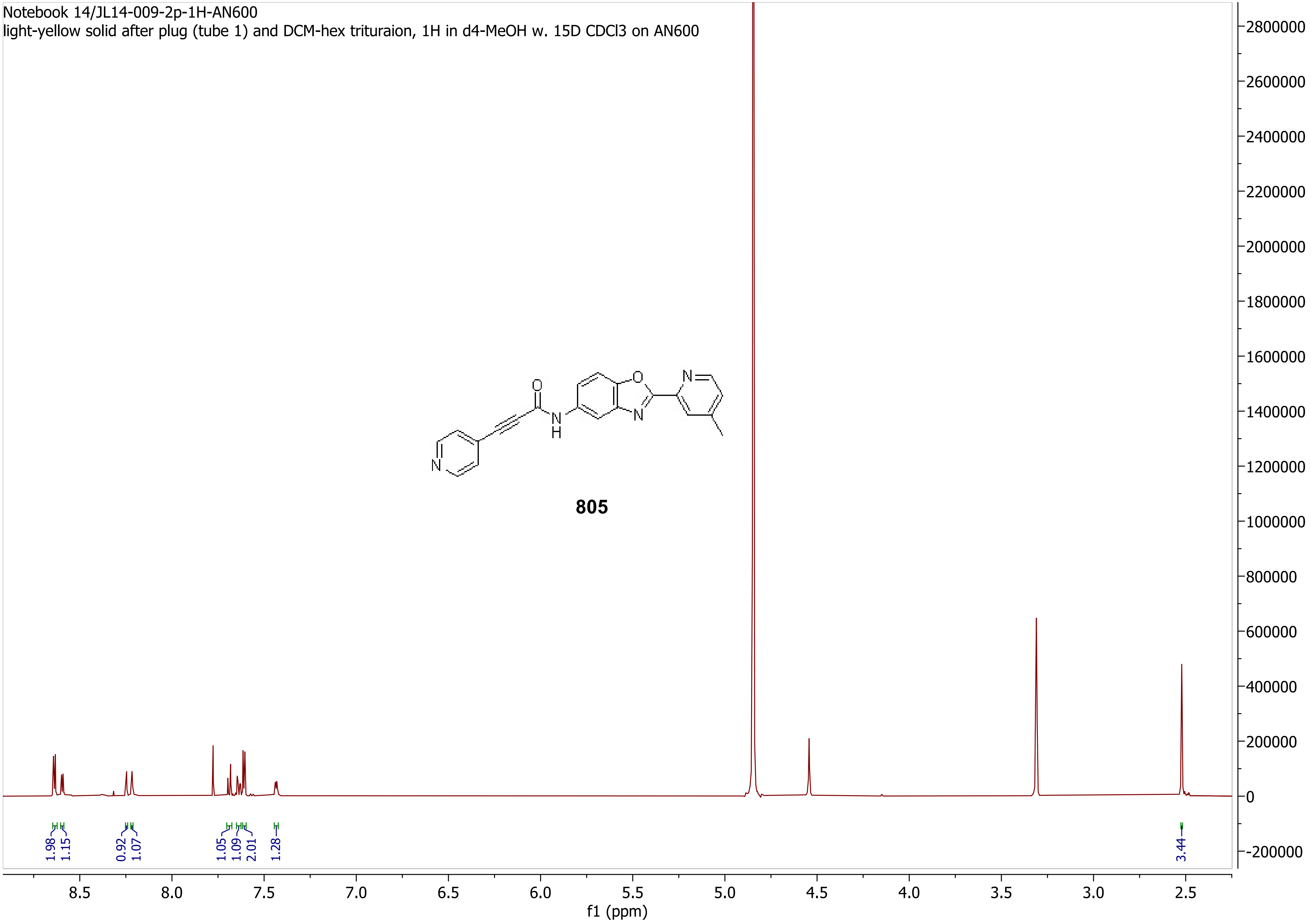
Notebook 14/JL14-009-2p-1H-AN600
light-yellow solid after plug (tube 1) and DCM-hex trituraion, 1H in d4-MeOH w. 15D CDCl3 on AN600
805
1.98
1.15
0.92
1.07
1.05
1.09
2.01
1.28
3.44
f1 (ppm)

Notebook 14/JL14-009-2p-13C-AN600
light-yellow solid after plug (tube 1) and DCM-hex trituraion, 1H in d4-MeOH w. 15D CDCl3 on AN600

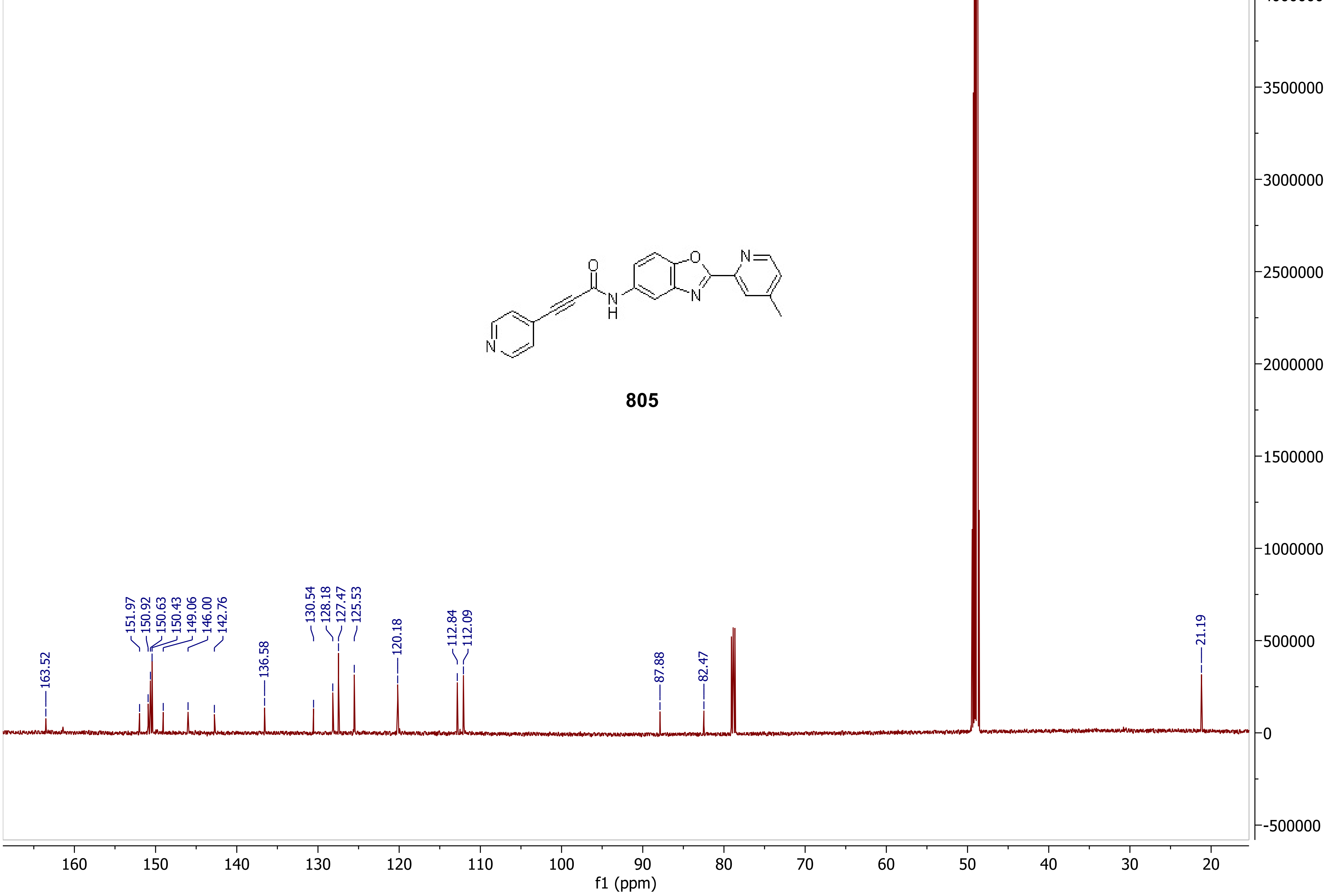

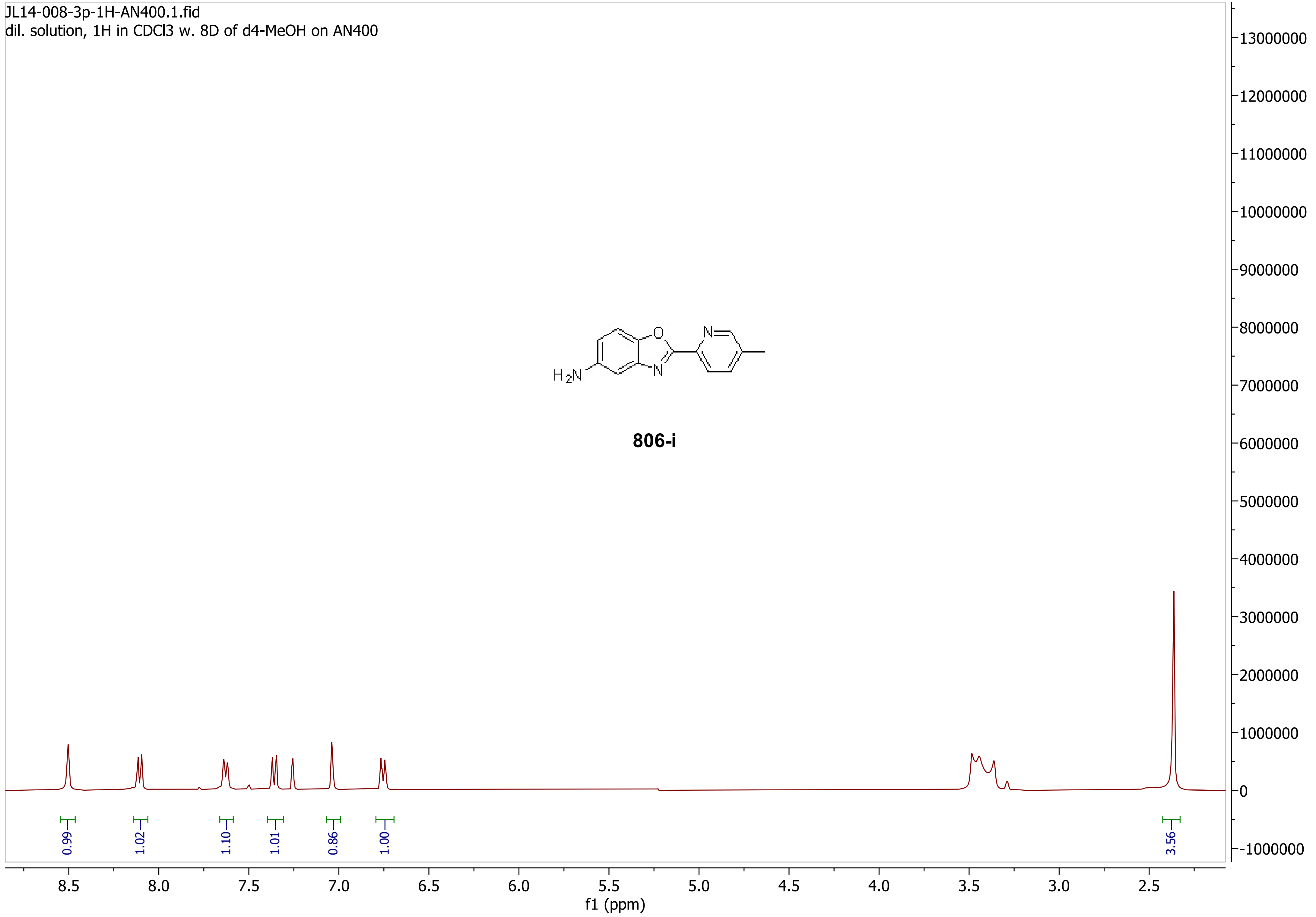
JL14-008-3p-1H-AN400.1.fid
dil. solution, 1H in CDCl3 w. 8D of d4-MeOH on AN400
H2N
O
N
N
806-i
0.99
1.02
1.10
1.01
0.86
1.00
3.56
8.5
8.0
7.5
7.0
6.5
6.0
5.5
5.0
4.5
4.0
3.5
3.0
2.5
f1 (ppm)
13000000
12000000
11000000
10000000
9000000
8000000
7000000
6000000
5000000
4000000
3000000
2000000
1000000
0
-1000000

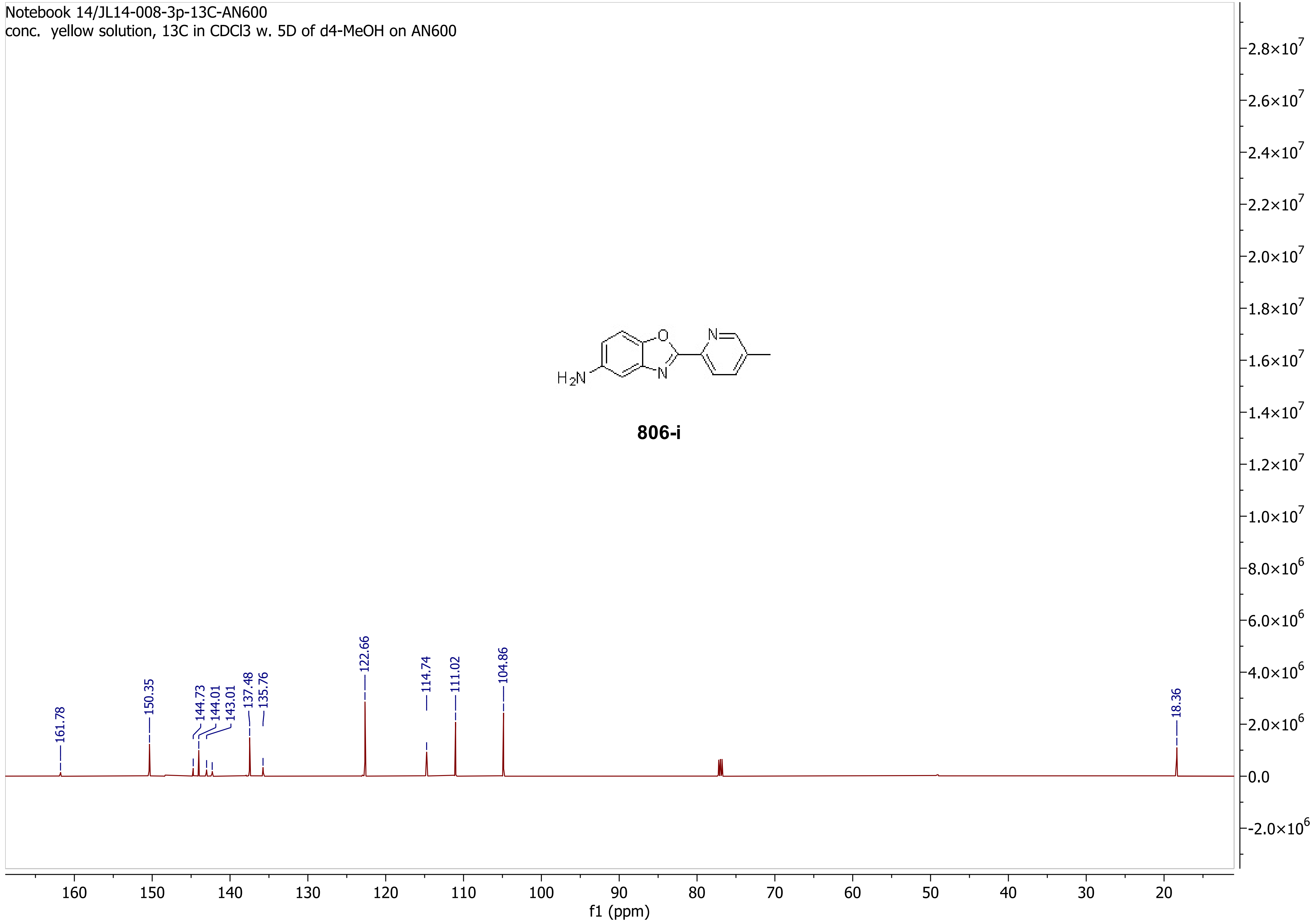
Notebook 14/JL14-008-3p-13C-AN600
conc. yellow solution, 13C in CDCl3 w. 5D of d4-MeOH on AN600
806-i
161.78
150.35
144.73
144.01
143.01
137.48
135.76
122.66
114.74
111.02
104.86
18.36
f1 (ppm)

Notebook 14/JL14-009-3p-1H-AN600
light-yellow solid after 5% MeOH-DCM plug, tube 3, 1H in d4-MeOH w. 15D CDCl3 on AN600

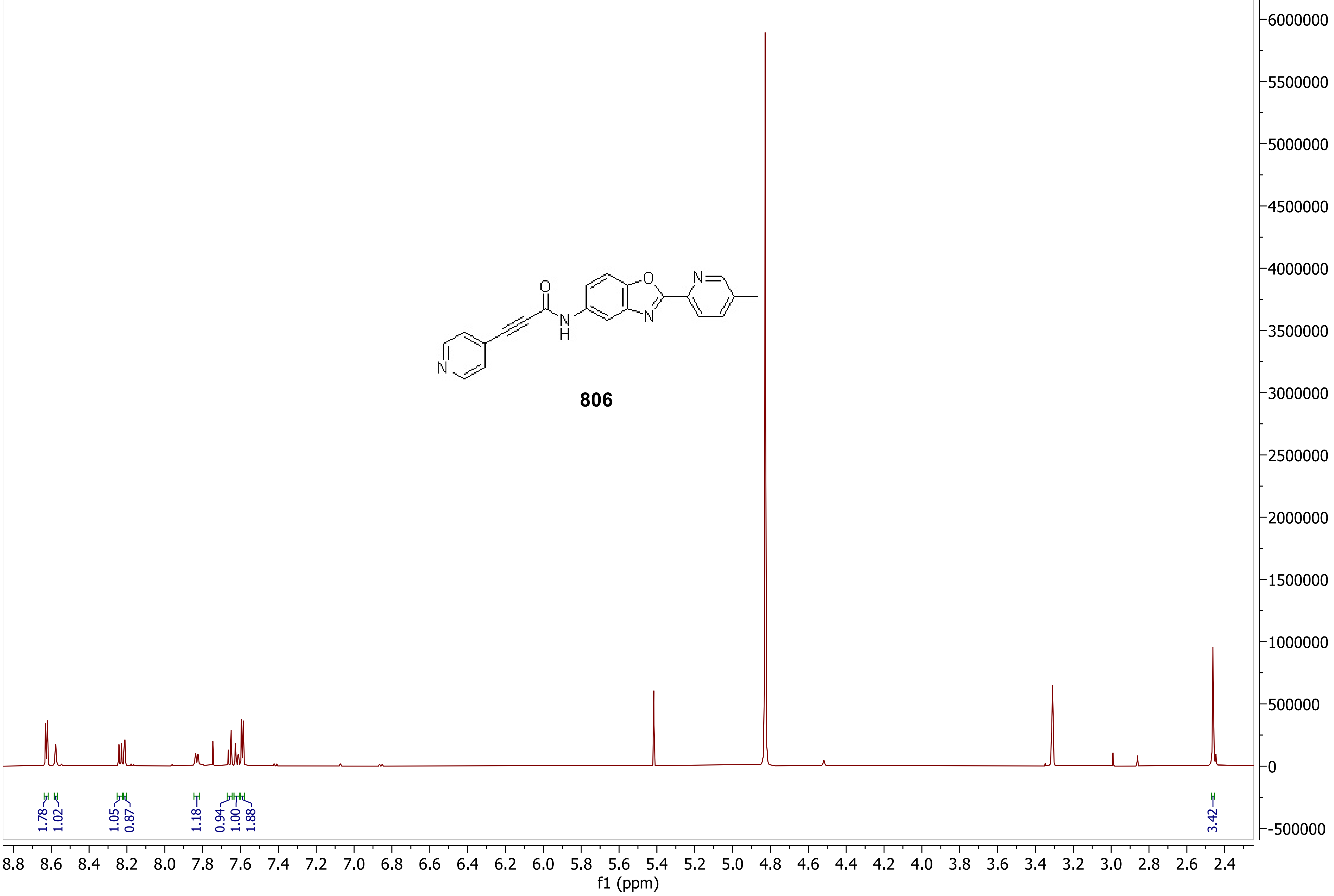

Notebook 14/JL14-009-3p-13C-AN600
light-yellow solid after 5% MeOH-DCM plug, tube 3, 13C in d4-MeOH w. 15D CDCl3 on AN600

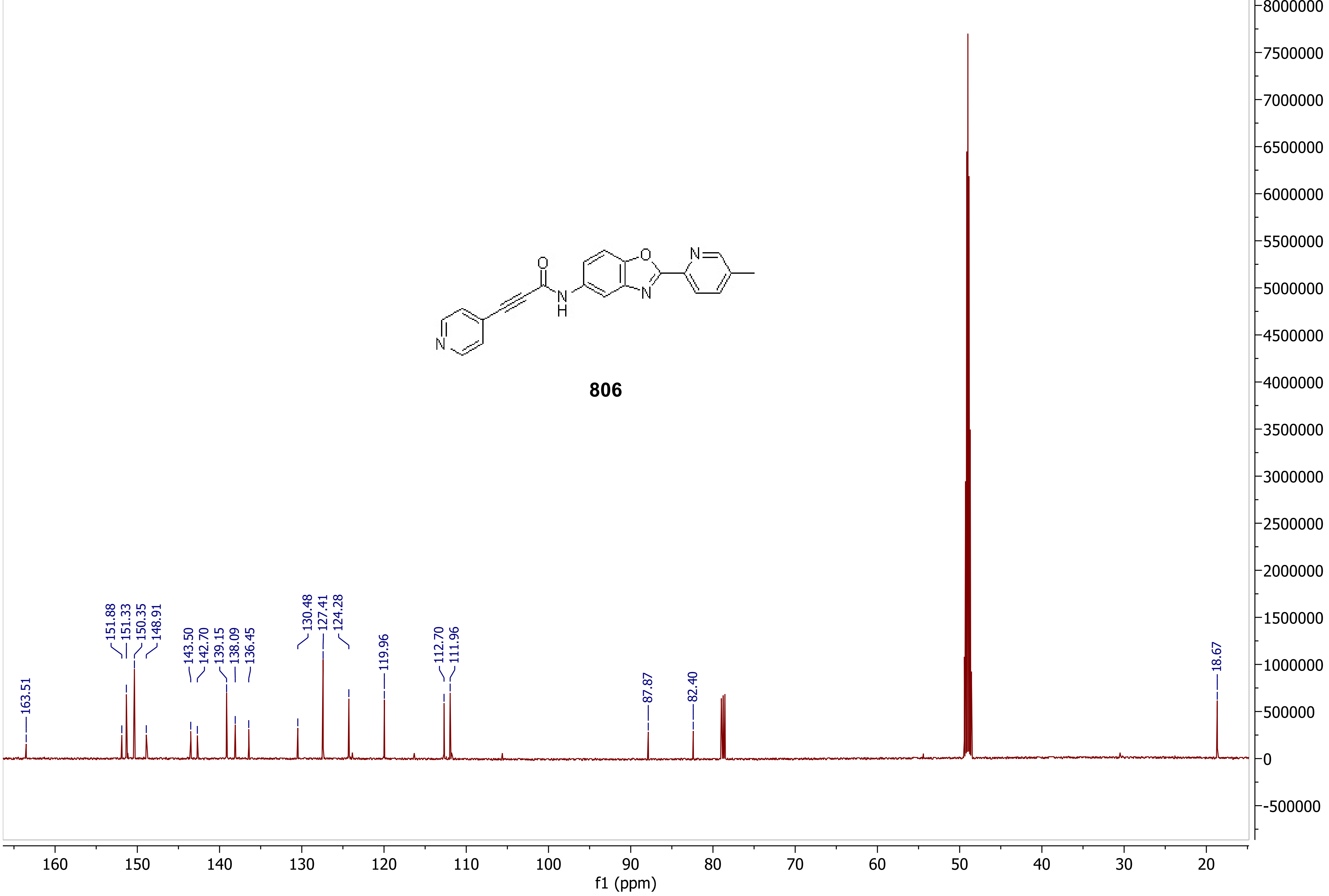

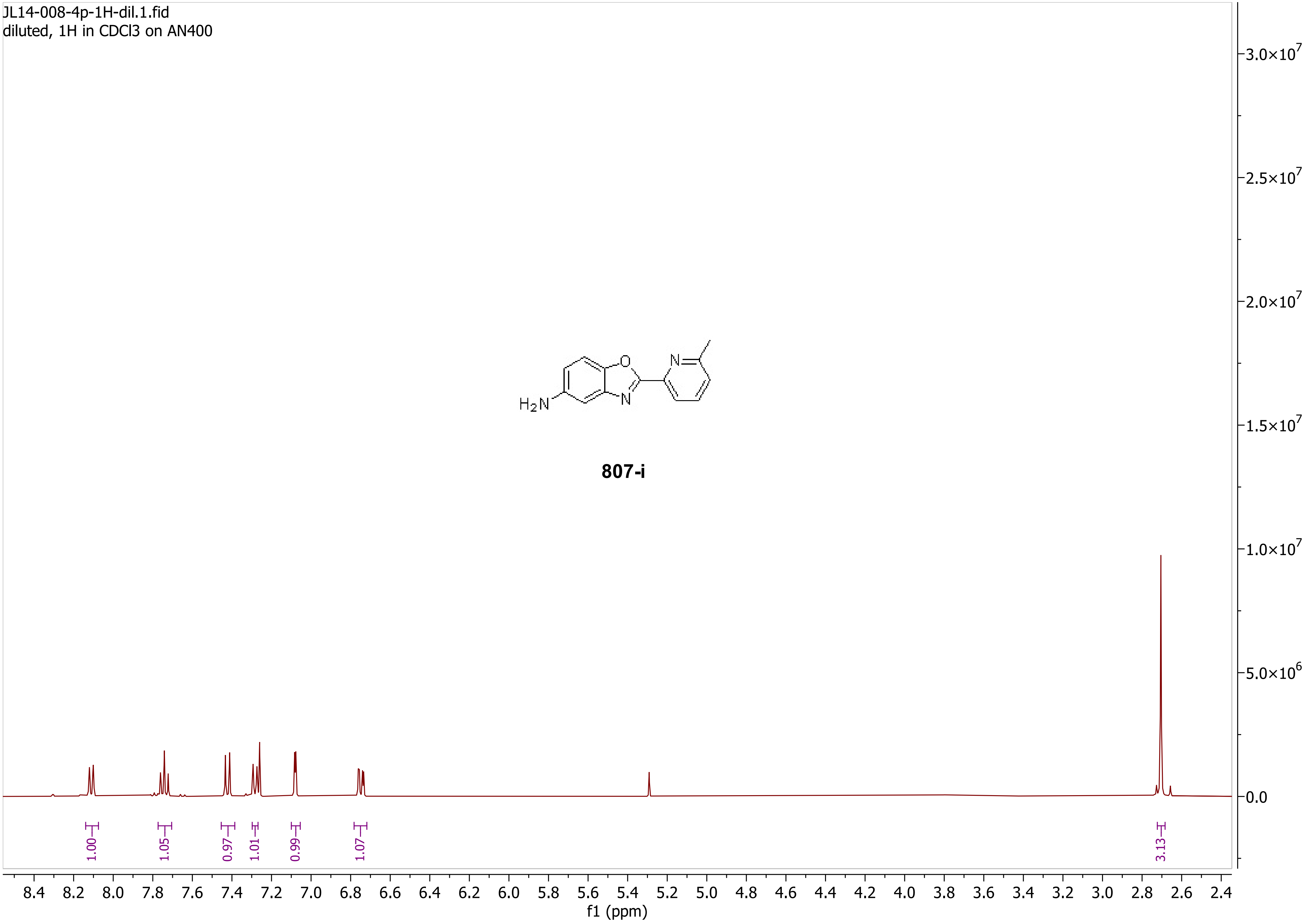
JL14-008-4p-1H-dil.1.fid
diluted, 1H in CDCl3 on AN400
H2N
O
N
N
807-i
1.00
1.05
0.97
1.01
0.99
1.07
3.13
8.4 8.2 8.0 7.8 7.6 7.4 7.2 7.0 6.8 6.6 6.4 6.2 6.0 5.8 5.6 5.4 5.2 5.0 4.8 4.6 4.4 4.2 4.0 3.8 3.6 3.4 3.2 3.0 2.8 2.6 2.4
f1 (ppm)
3.0×10^7
2.5×10^7
2.0×10^7
1.5×10^7
1.0×10^7
5.0×10^6
0.0

JL14-008-4p-13C-AN400.3.fid
orange solid after extraction & 5% MeOH-DCM plug, 13C in CDCl3 on AN400

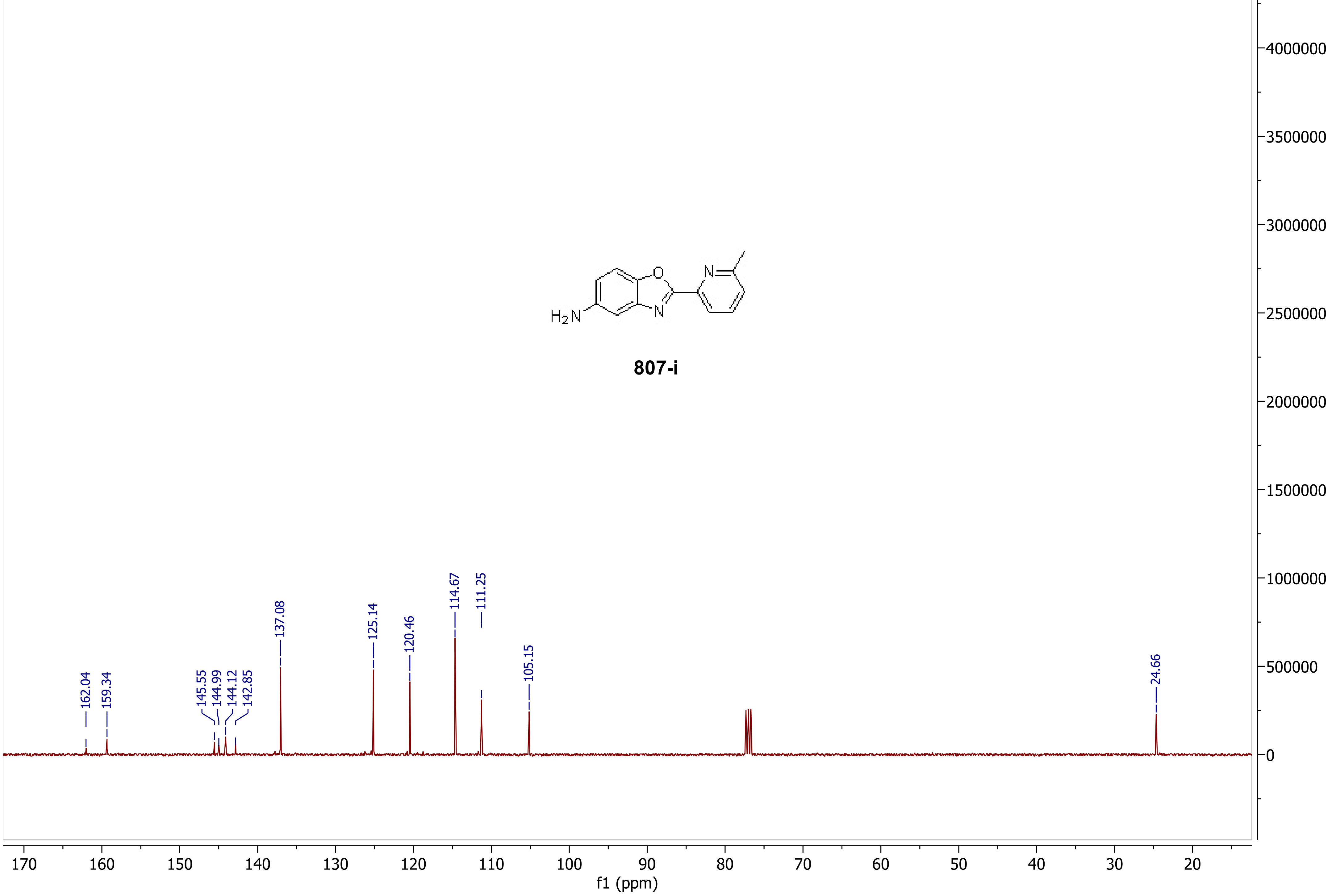

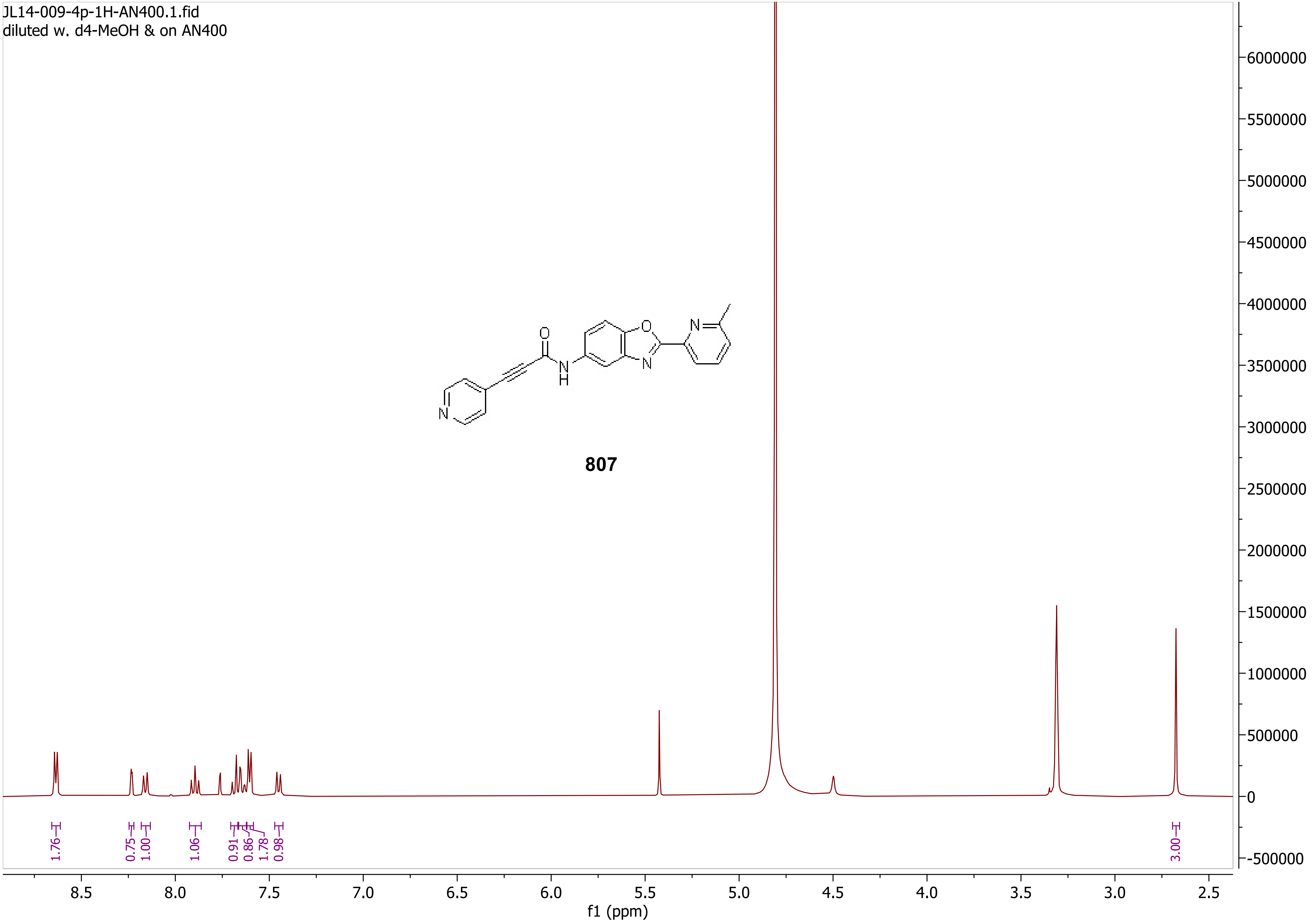
JL14-009-4p-1H-AN400.1.fid
diluted w. d4-MeOH & on AN400
807
1.76
0.75
1.00
1.06
0.91
0.86
1.78
0.98
3.00
8.5
8.0
7.5
7.0
6.5
6.0
5.5
5.0
4.5
4.0
3.5
3.0
2.5
f1 (ppm)
6000000
5500000
5000000
4500000
4000000
3500000
3000000
2500000
2000000
1500000
1000000
500000
0
-500000

Notebook 14/JL14-009-4p-13C-AN600-t2
light-yellow solid after 5% MeOH-DCM plug & DCM-hex trituration, tube 2, 13C in d4-MeOH w. 15D CDCl3 on AN600

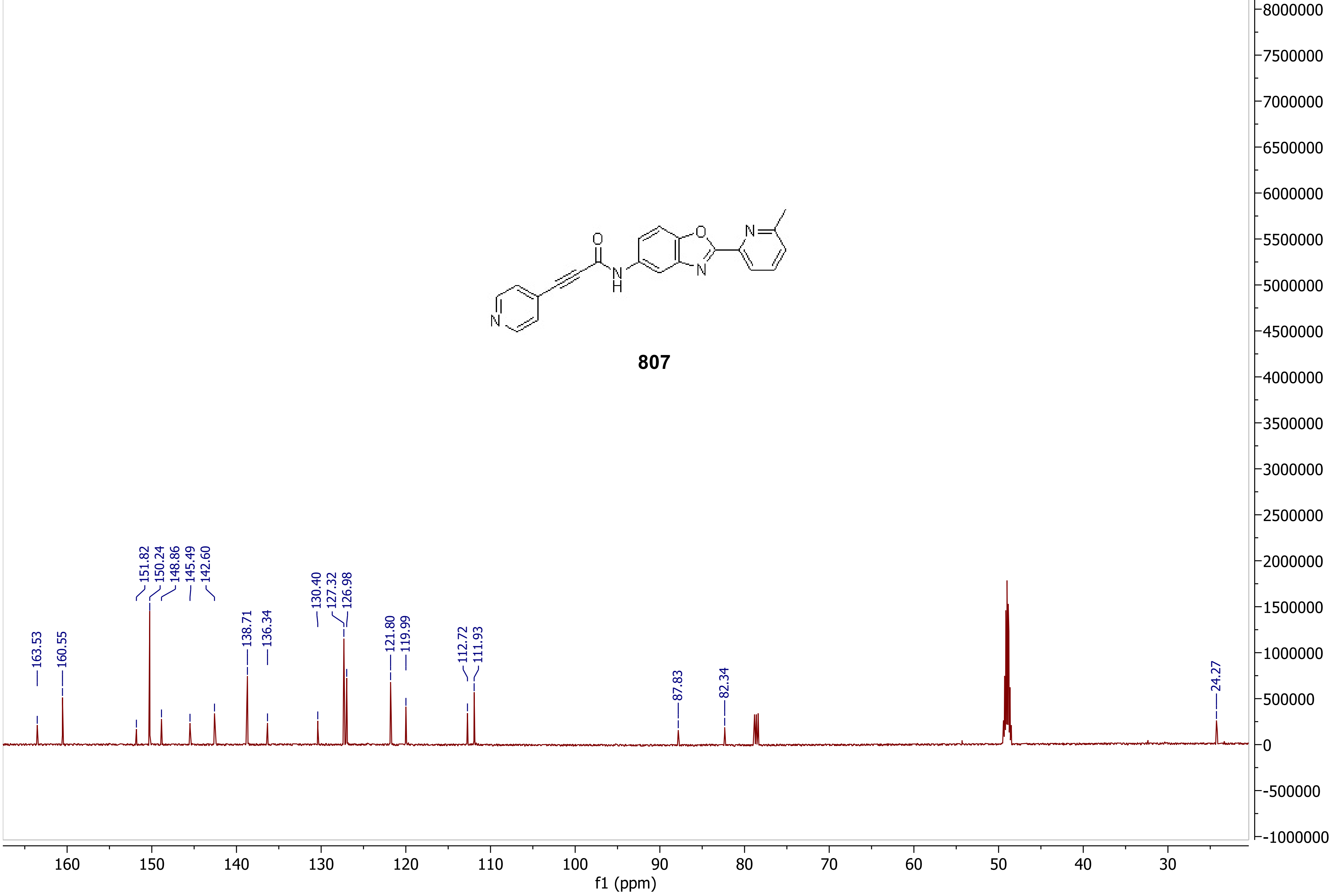

Notebook 14/JL14-008-5p-cr-1H-AN600
yellow solid after DCM extraction and triuration in DCM-hex, supernatant, 1H in CDCl3 on AN600

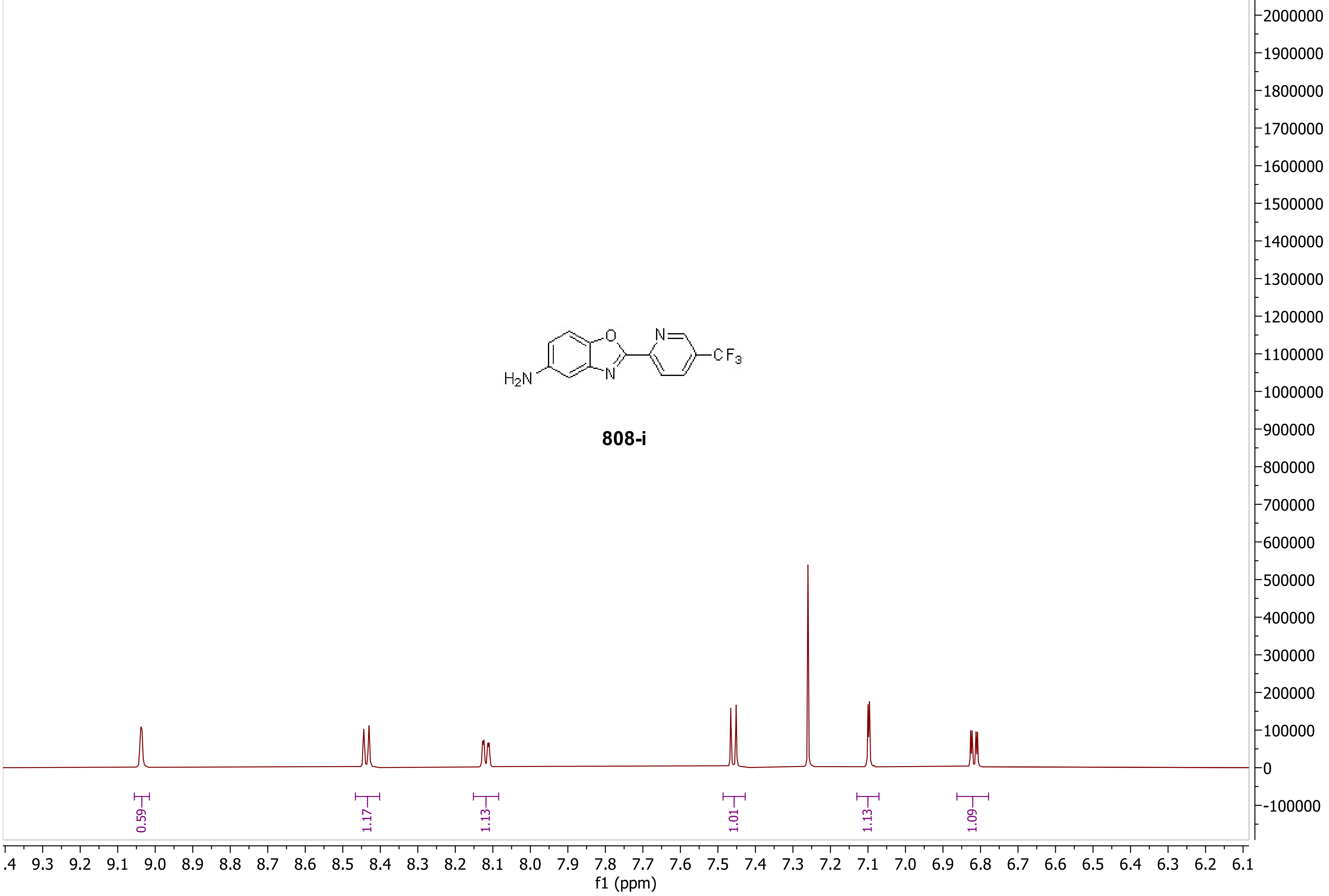

Notebook 14/JL14-008-5p-cr-13C-AN600
yellow solid after DCM extraction and triuration in DCM-hex, supernatant, 13C in CDCl3 on AN600

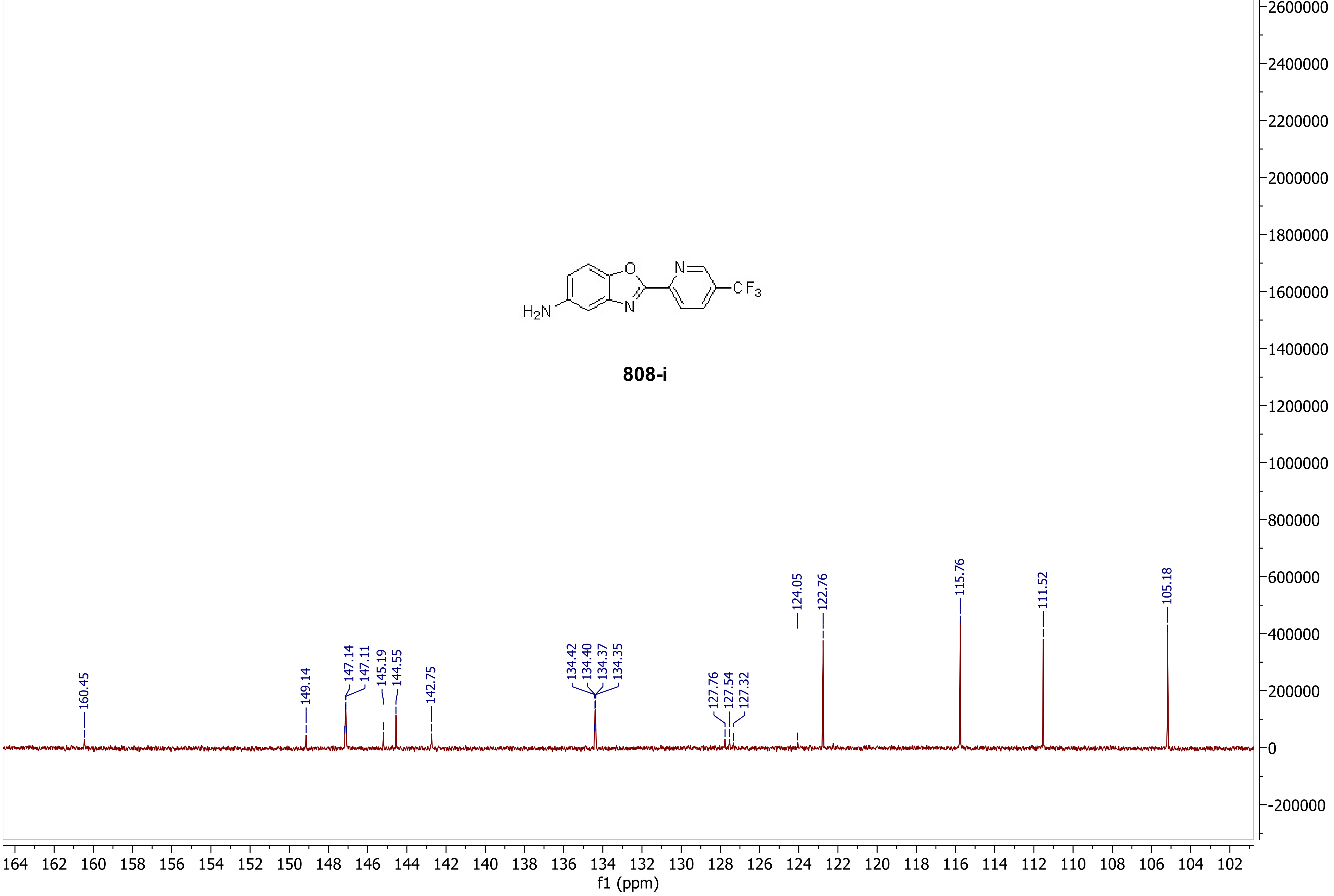

Notebook 14/JL14-009-5p-1H-MeOH
1H in d4-MeOH w. 20D CDCl3 on AN600

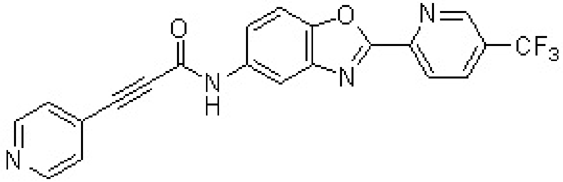


**808**

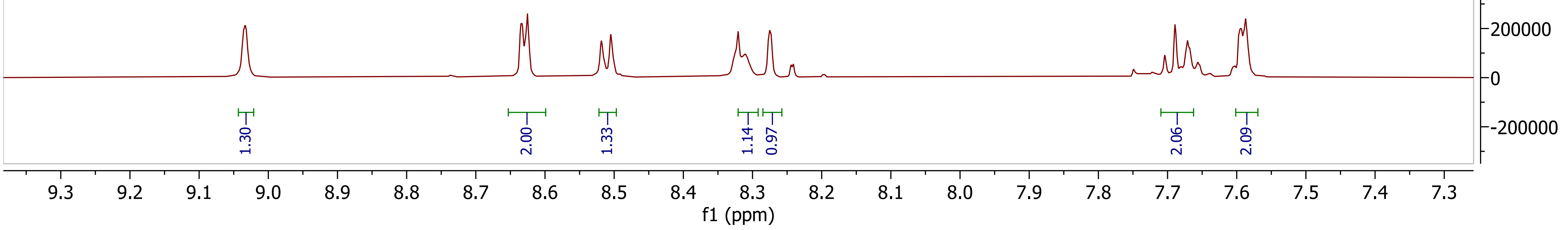

Notebook 14/JL14-009-5p-13C-MeOH
13C in d4-MeOH w. 20D CDCl3 on AN600

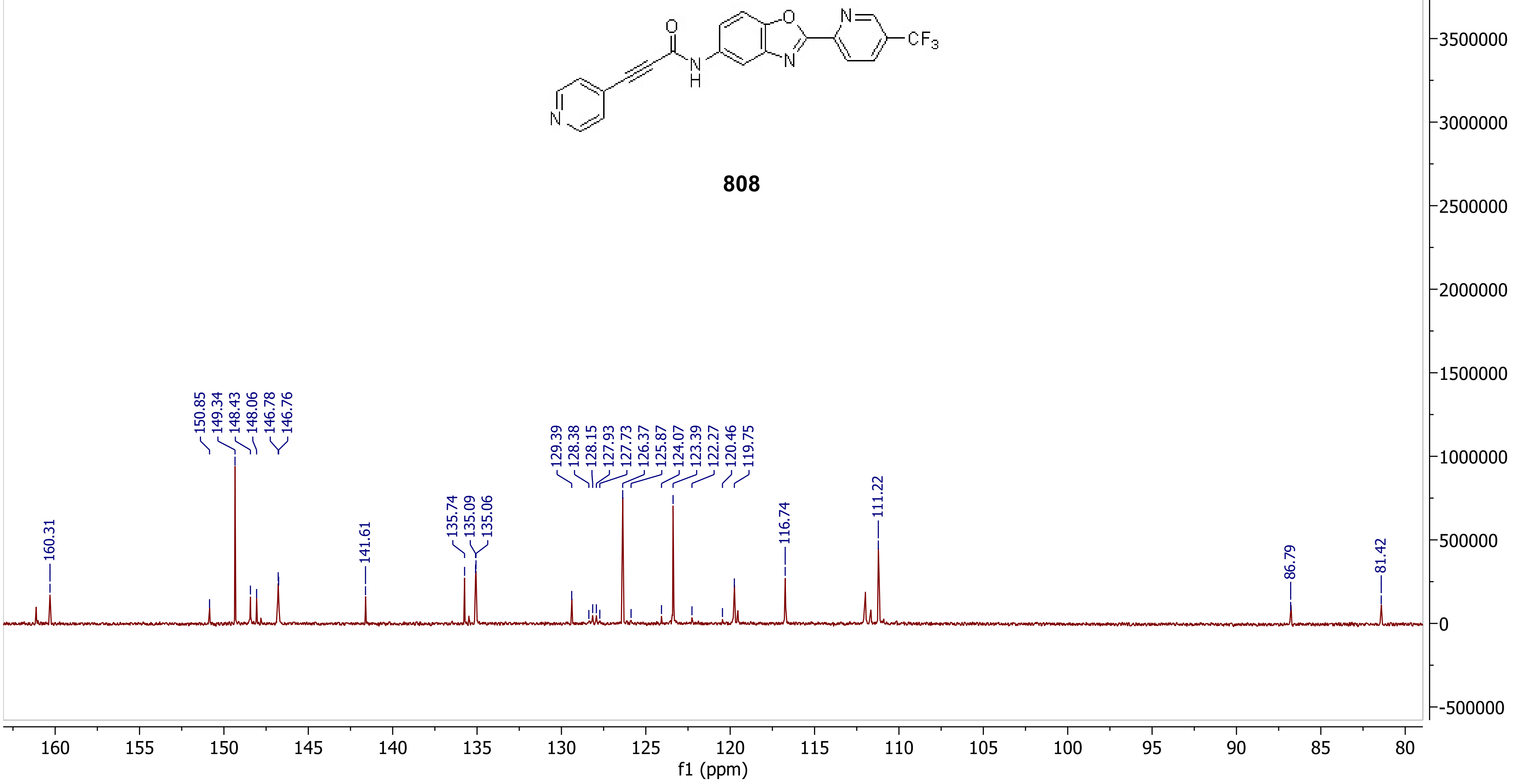
